# Myosin VIII and XI isoforms interact with *Agrobacterium* VirE2 protein and help direct transport from the plasma membrane to the perinuclear region during plant transformation

**DOI:** 10.1101/2023.03.06.531343

**Authors:** Nana Liu, Lan-Ying Lee, Yanjun Yu, Stanton B. Gelvin

**Affiliations:** Department of Biological Sciences, Purdue University, West Lafayette, IN 47907

## Abstract

Virulent *Agrobacterium* strains transfer single-strand T-DNA (T-strands) and virulence effector proteins into plant cells. VirE2, one of these virulence effectors, enters the plant cell and is thought to bind T-strands, protecting them from nuclease degradation and helping guide them to the nucleus. How VirE2 is trafficked inside the plant cell is not fully understood. Using bimolecular fluorescence complementation, *in vitro* pull-down, yeast two-hybrid, and *in vivo* co-immunoprecipitation assays, we found that VirE2 binds directly to the cargo binding domains of several myosin VIII family members, and to myosin XI-K. We observed reduced susceptibility of several *Arabidopsis* actin mutants and a myosin *VIII-1/2/a/b* quadruple mutant to *Agrobacterium*-mediated transformation. Expression of cargo binding domains of myosin VIII-1, VIII-2, VIII-A, or VIII-B in transgenic plants inhibits *Arabidopsis* root transformation. However, none of the myosin VIII proteins contribute to the intracellular trafficking of VirE2. Expression of myosin *VIII-2*, *-A*, *-B*, but not *VIII-1*, cDNAs in the myosin *VIII-1/2/a/b* mutant partially restored transformation. Furthermore, functional fluorescently-tagged VirE2, synthesized in plant cells, relocalized from the cellular periphery into the cytoplasm after delivery of T-strands from *Agrobacterium*. Surprisingly, mutation of myosin *XI-k* and expression of the myosin XI-K cargo binding domain had no effect on transformation, although it blocked VirE2 movement along actin filaments. We hypothesize that myosin VIII proteins facilitate VirE2 tethering to the plasma membrane and are required for efficient localization of VirE2 to membrane sites from which they bind incoming T-strands. Myosin XI-K is important for VirE2 movement through the cytoplasm towards the nucleus.

## INTRODUCTION

The soil-borne phytopathogen *Agrobacterium tumefaciens* is the causative agent of crown gall disease in plants (Gelvin, 2000; Tzfira et al., 2004). The ability of *Agrobacterium* to genetically transform plant cells has made *Agrobacterium*-mediated transformation the most widely used platform for generating transgenic plants (Gelvin, 2017). During transformation, T-DNA, a defined segment of the resident tumor inducing (Ti) plasmid, is transferred to the plant cell along with several virulence (Vir) effector proteins (Bevan and Chilton, 1982). Virulence effector proteins VirD5, VirE2, VirE3, and VirF, along with VirD2, which covalently links to T-strands (single-strand T-DNA molecules), are transferred from *Agrobacterium* to the plant through a type IV secretion system (T4SS) (Cascales and Christie, 2004; Christie et al., 2005).

VirE2, the most abundant of the virulence effector proteins (Engstrom et al., 1987), is important for efficient plant transformation as *virE2* mutant *Agrobacterium* strains are highly attenuated in virulence (Stachel and Nester, 1986; Rossi et al., 1996). *In vitro*, VirE2 binds to single-strand (ss)DNA in a sequence-independent manner and has been proposed to interact with single-strand T-strands, linked to VirD2, in the plant cytoplasm, forming a “T-complex” and protecting T-strands from nuclease degradation (Citovsky et al., 1989; Sen et al., 1989; Howard et al., 1990; Yusibov et al., 1994; Rossi et al., 1996; Gelvin, 1998). VirE2 does not interact with T-strands in *Agrobacterium* (Cascales and Christie, 2004) and its interaction with T-strands to form a T-complex *in planta* has only been inferred. In addition to its *in vitro* DNA binding activity, VirE2 can form membrane channels which permit the transport of ssDNA through artificial membranes (Dumas et al., 2001; Duckely et al., 2005). Recent research showed that VirE2 is anchored on the host plasma membrane through interaction with VirE3 at *Agrobacterium*-host contact sites. Such membrane associations may facilitate the proposed interactions between VirE2 and T-strands as the T-strands enter the plant cell (Li et al., 2018).

Efficient plant transformation by a *virE2* mutant *Agrobacterium* strain (the “T-DNA donor”) can be restored by co-inoculation with a bacterial strain lacking T-DNA but expressing VirE2 (the “VirE2 donor”; Otten et al., 1984; Citovsky et al., 1992; Simone et al., 2001), and a *virE2* mutant *Agrobacterium* strain can be complemented to full virulence by expression of a *VirE2* transgene *in planta* (Citovsky et al., 1992; Lapham et al., 2021), indicating that VirE2 is functional in plants during transformation. However, the subcellular localization of VirE2 remains controversial. Early reports demonstrated nuclear localization of VirE2 (Citovsky et al., 1992), but other studies from several laboratories indicated that VirE2 localizes in the cytoplasm (Gelvin, 2010). A recent report claimed that a small amount of VirE2 can enter the nucleus, but only in the presence of VirD2 and T-strands (Li et al., 2020). How VirE2, and potentially associated T-strands, are trafficked from their entry point at the cell periphery to the nucleus remains unknown.

Myosins, motor proteins that convert chemical energy into directed movement of cargo proteins along actin filaments, are key players in trafficking proteins and organelles in plants (Vale, 2003; Richards and Cavalier-Smith, 2005). Two classes of myosins, myosin VIII and XI, are encoded by plant genomes. *Arabidopsis thaliana* encodes 13 myosin XI group members and four myosin VIII group members (Bezanilla et al., 2003; Foth et al., 2006). Generally, myosin proteins contain a highly conserved N-terminal motor domain for actin binding, a neck domain with a number of IQ repeats for light-chain binding, a coiled-coil domain that is responsible for myosin protein dimerization, and a specific C-terminal tail domain that binds cargo (Trybus, 2008). Class XI myosins, known as organelle transporters, colocalize with organelles and are involved in the rapid trafficking of Golgi stacks, streaming of endoplasmic reticulum (ER), and cellular remodeling (Lee and Liu, 2004; Reisen and Hanson, 2007; Avisar et al., 2008b; Sparkes et al., 2009; Ueda et al., 2010). In addition, myosins XI-K and XI-2 play major and overlapping roles in root hair development (Peremyslov et al., 2008). In contrast to class XI myosins, considerably less is known about the intracellular functions of class VIII myosins. Class VIII myosins colocalize with plasmodesmata, the endoplasmic reticulum, and the plasma membrane (Golomb et al., 2008) and may function in intercellular protein and RNA delivery to the plasmodesmata, in endocytosis, and in anchoring actin filaments at plasmodesmata sites (Reichelt et al., 1999; Wojtaszek et al., 2005; Avisar et al., 2008a; Golomb et al., 2008; Sattarzadeh et al., 2008). Moreover, myosin VIII is important for the cell-to-cell movement of virus proteins, and is required for viral movement protein targeting to and virus trafficking through plasmodesmata (Amari et al., 2014).

The actin cytoskeleton functions to organize the endomembrane system and trafficking patterns within the cell (Smith and Oppenheimer, 2005; Hussey et al., 2006). Cargo-binding and intracellular transport by myosin is proposed to exert force on the actin filaments (Staiger et al., 2009). Yang et al. (2017) reported that trafficking of VirE2 inside plant cells is powered by myosin XI-K via the endoplasmic reticulum and F-actin filaments. However, our knowledge of the roles of plant myosins in specific steps of VirE2 trafficking and transformation remains sparse. In the present study, we investigated the roles of myosin VIII family members and myosin XI-K in *Agrobacterium*-mediated transformation and in VirE2 intracellular movement. Using a dominant-negative approach by overexpressing individual myosin cargo-binding domains, and by overexpressing various individual full-length myosin cDNAs, we show that *Arabidopsis* myosins VIII-2, VIII-A, and VIII-B, but not XI-K, are important for transformation. These three myosin VIII isoforms interact with VirE2 and retain it at the periphery of root cells. When T-DNA enters the plant cell, a portion of peripherally localized VirE2 relocalizes into the cytosol, especially the perinuclear region. However, class VIII myosins do not affect the movement of VirE2 along actin filaments within the cytoplasm. In contrast, myosin XI-K supports the intracellular movement of VirE2. We propose that myosins VIII-2, VIII-A, and VIII-B tether VirE2 to the plasma membrane. Entry of VirD2/T-strands into the plant cell releases VirE2 from the membrane, following which VirE2 translocates to the perinuclear region along actin filaments, powered by myosin XI-K.

## RESULTS

### The actin cytoskeleton is important for *Agrobacterium*-mediated transformation

The actin cytoskeleton is conserved in diverse eukaryotic organisms and is important for a variety of plant developmental and other processes, including cell growth, division, and expansion; organelle motility; organization of the cellular interior; endomembrane trafficking; and host defense responses (Smith and Oppenheimer, 2005; Embley and Martin, 2006; Hussey et al., 2006; Szymanski and Cosgrove, 2009; Peremyslov et al., 2010). To test whether the actin cytoskeleton plays a role in *Agrobacterium*-mediated transformation, we conducted transient and stable *Arabidopsis* root transformation assays using wild-type plants and plants individually mutant in two root-expressed actin genes, *actin2* (*act2*) and *actin7* (*act7*). These actin mutant plants showed substantial reductions (three-to five-fold) in transformation. Introduction of a wild-type *Act7* cDNA into the *act7-4* mutant restored both transient and stable transformation (Figure 1A). These results show that root-expressed actin genes are important for transformation of *Arabidopsis* roots. We also conducted root transformation assays using a mutant of the pollen-expressed *actin12* gene (*act12*) and the *botero* (*bot1*) mutant that shows disorganization of the root cortical microtubule network (Bichet et al., 2001). These mutants showed wild-type levels of transformation (Figure 1A), indicating that neither a pollen-expressed actin gene nor the correct organization of root microtubules is essential for *Agrobacterium*-mediated root transformation.

**Figure 1.**
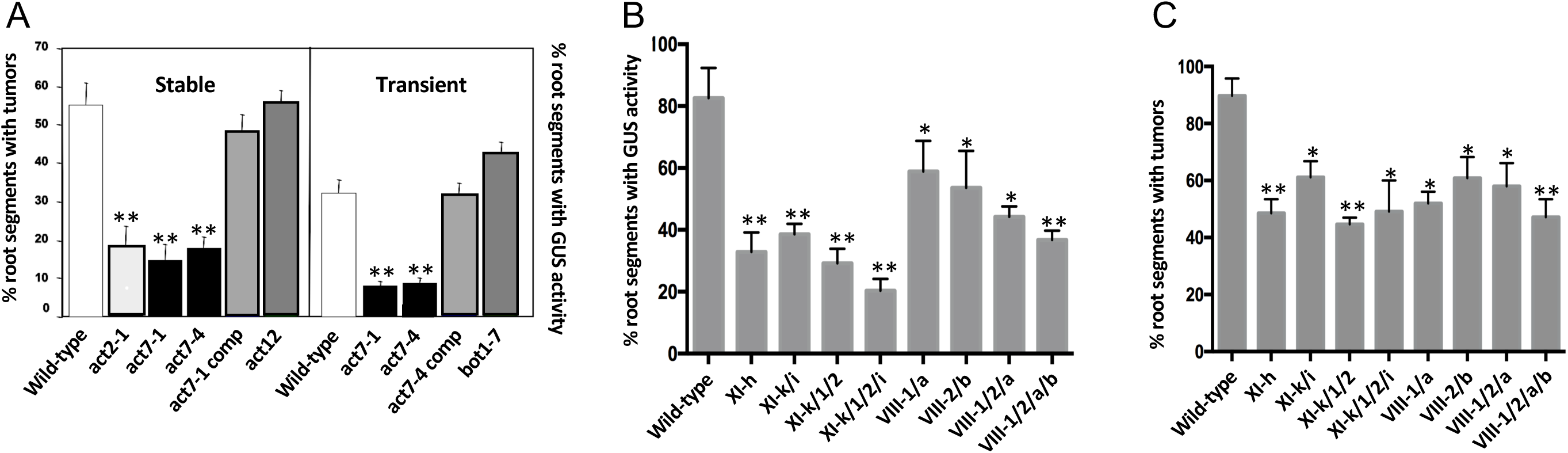
Some *Arabidopsis* actin and myosin genes are important for *Agrobacterium*-mediated transformation. (A) Root segments from *Arabidopsis* wild-type and *act2-1*, *act7-1*, *act7-4*, *act12*, *act7-*4 comp (complemented), or *bot1* mutants were infected with *A. tumefaciens* A208 (tumorigenic strain for stable transformation; left panel) or *A. tumefaciens* At849 (T-DNA contains a plant-active *gusA*-intron gene for transient transformation; right panel). After two days, the root segments were moved to medium (MS lacking phytohormones for stable transformation, CIM for transient transformation) containing timentin to kill the bacteria. For stable transformation, the presence of tumors was scored one month after infection. For transient transformation, root segments were stained with X-gluc after an additional four days, and the percentage of roots showing GUS activity was calculated. *Act7-4* comp, *act7-4* mutant complemented with an *ACT7* cDNA. **(B and C)** Root segments from *Arabidopsis* wild-type and single or higher order *myosin VIII* or *XI* mutants were infected with *A. tumefaciens* At849 transiently or *A. tumefaciens* A208 stably. A total of 10-15 plants and >100 segments per plant were tested for stable transformation. Values given are means ± SE. Asterisks indicate significant differences compared to wild-type plants. [*t-test*, \**P* < 0.05; \*\**P* < 0.01].

Yang et al. (2017) showed that the *Agrobacterium* virulence effector protein VirE2 uses myosin XI-K to traffic through the tobacco cytoplasm. Given the importance of VirE2 for transformation, we individually tested *Arabidopsis* mutants in each of the 13 myosin XI and four myosin VIII gene family members for their susceptibility to transient and stable *Agrobacterium*-mediated transformation. Figures 1B and C, and Supplemental Figure S1, show that apart from the *myosin XI-h* mutant, mutation of no other single myosin gene affected transformation. In particular, mutation of the *myosin XI-k* gene did not decrease transformation. Because of potential functional redundancy among the plant myosin VIII and XI family members (Prokhnevsky et al., 2008), we also tested several higher order myosin VIII and myosin XI mutants. Higher order *myosin XI-k* mutants (*XI-k/i*, *XI-k/1/2*, and *XI*-*k/1/2/i*) showed decreased transformation susceptibility (Figures 1B and 1C). Similarly, the higher order myosin VIII mutants *1/a*, *2/b*, *1/2/a*, and *1/2/a/b* showed reduced transformation susceptibility compared to wild-type Col-0 plants (Figures 1B and 1C).

We compared root and above-ground plant growth of each of the higher order myosin XI and myosin VIII mutants. Similar to what was previously observed (Ojangu et al., 2007; Peremyslov et al., 2008), the myosin *XI-k/1/2/i* quadruple mutant displayed a strong growth and developmental phenotype compared with Col-0 plants: the main roots had fewer lateral branches and shorter root hairs, and the plants were dwarf. However, the myosin *VIII 1/2/a/b* quadruple mutant appeared similar to Col-0 plants in growth, root development, and the ability to form calli from cut root segments (Supplemental Figures S2A and S2B). Roots and the crown of the plant are the natural target tissues for *Agrobacterium*-mediated transformation. Because we were conducting quantitative root transformation assays (Gelvin, 2006), we did not continue to investigate transformation susceptibility of the abnormal roots of the myosin *XI-k/1/2/i* quadruple mutant. We therefore turned our attention to investigating the importance of the various myosin VIII isoforms for transformation, and to the myosin XI-K protein.

### Expression of full-length myosin *VIII-2*, *VIII-A*, and *VIII-B* cDNAs, but not a myosin *VIII-1* cDNA, increases susceptibility of the myosin *VIII-1/2/a/b* quadruple mutant to *Agrobacterium*-mediated transformation

To determine the importance of each *myosin VIII* gene for *Agrobacterium*-mediated root transformation, we individually expressed cDNAs for each *myosin VIII* gene in the *myosin VIII-1/2/a/b* quadruple mutant. We placed each *myosin VIII* cDNA under the control of a strong Cauliflower Mosaic Virus (CaMV) double 35S promoter (hereafter reported in Supplemental Table 1 as the *Agrobacterium* strains used to make the transgenic lines; At2361-At2164; At2360 was used as an empty vector control) or a β-estradiol-inducible promoter (Zuo et al., 2000; At2389-2392) and generated multiple transgenic lines. Because of the importance of myosins in trafficking molecules and organelles through the plant cytoplasm (Avisar et al., 2008b), we hereafter limited our transformation assays to transient transformation which measures T-DNA and virulence effector protein movement through the cytoplasm, entry into the nucleus, and expression of non-integrated transgenes. Thus, these assays predominantly measure non-integrated transgenes.

Transient root transformation assays of five independent transgenic lines of each *myosin VIII* cDNA indicated that individual expression of the *myosin VIII-2*, *VIII-A*, and *VIII-B* cDNAs increased the transformation susceptibility of the *myosin VIII-1/2/a/b* quadruple mutant (Figure 2A and Supplemental Figures S3B-D), indicating a role for these myosins in *Agrobacterium*-mediated transformation. However, expression of the *myosin VIII-1* cDNA in the quadruple mutant background did not increase transformation susceptibility (Figure 2A and Supplemental Figure S3A). This latter result suggests either that myosin VIII-1 does not contribute to transformation or that expression of myosin VIII-1 is not sufficient to increase transformation in the absence of the other three myosin VIII isoforms. To distinguish between these two possibilities, we individually overexpressed each *myosin VIII* cDNA in wild-type Col-0 plants. Interestingly, overexpression of the *myosin VIII-1* cDNA, but not the *myosin VIII-2*, *VIII-A*, or *VIII-B* cDNAs, increased susceptibility of Col-0 plants to *Agrobacterium*-mediated transformation (Figure 2B and Supplemental Figures S3A-D). These results indicate that myosin VIII-1 can contribute to transformation, but that expression of myosin VIII-1 is not sufficient to alter transformation susceptibility in the absence of the other myosin VIII isoforms. Transgenic plants harboring inducible *myosin VIII* cDNAs yielded results similar to that of plants constitutively expressing these cDNAs (Supplemental Figures S4A-D).

**Figure 2.**
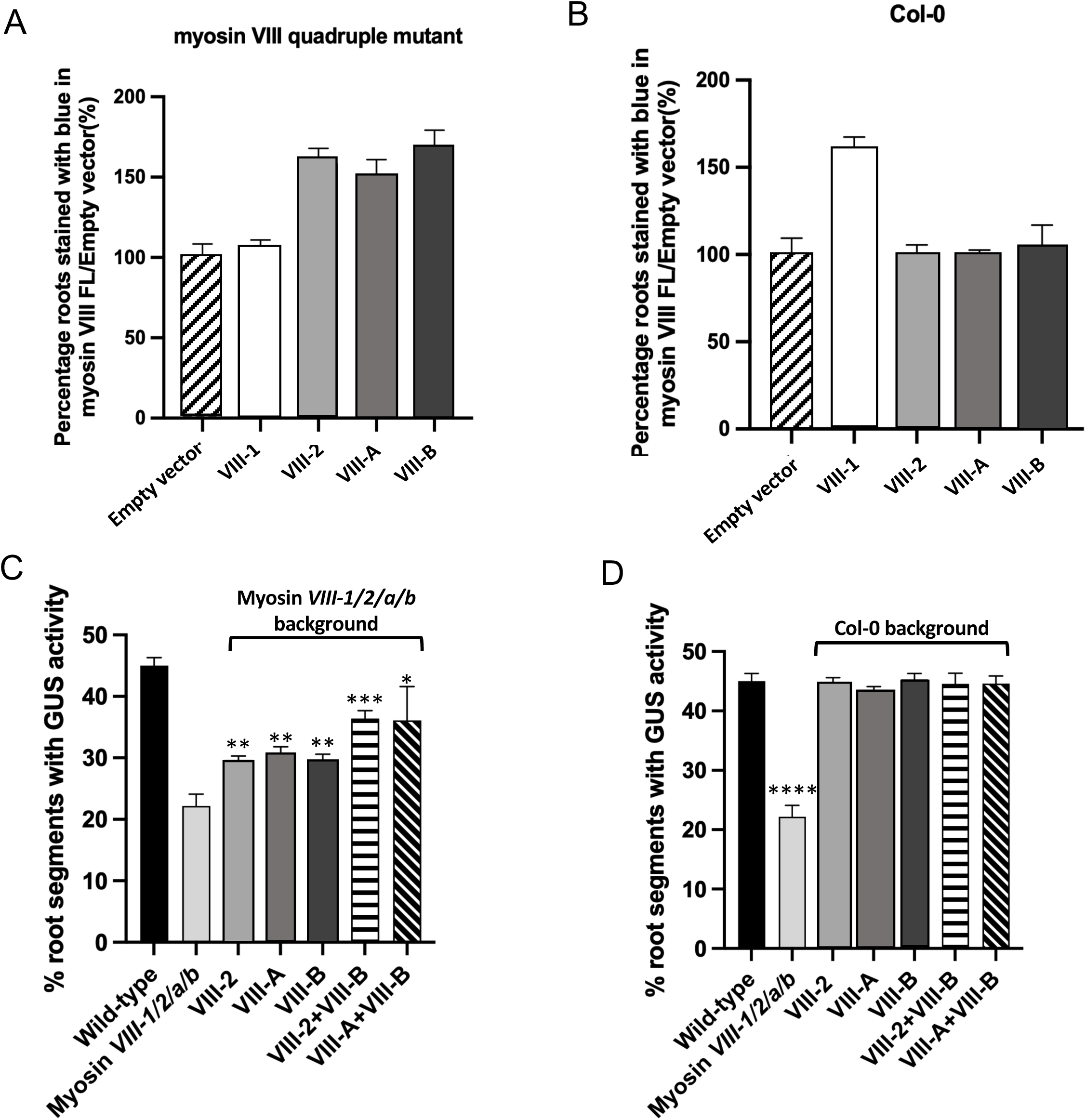
All Myosin VIII family members can contribute to *Agrobacterium*-mediated transformation. Root segments from transgenic *Arabidopsis* individually expressing either an empty vector or *myosin VIII* cDNAs in *myosin VIII-1/2/a/b* mutant plants **(A)**, or in wild-type plants **(B)**, were infected with *A. tumefaciens* At849. Transgenic *Arabidopsis* expressing cDNAs encoding *myosins VIII-2, VIII-A, VIII-B, VIII-2*+*VIII-B* or *VIII-A*+*VIII-B* in the *myosin VIII-1/2/a/b* mutant background **(C)** or in wild-type Col0 plants **(D)** were similarly infected with *A. tumefaciens* At849. Transient transformation efficiencies are indicated as the percentage of roots showing GUS activity six days after infection. A total of 10-15 plants and >100 segments per plant, from five independent transgenic lines, were tested for transient transformation. Values given are means ± SE Asterisks indicate significant differences compared to control plants that contain an empty vector (**A** and **B**), the *myosin VIII-1/2/a/b* quadruple mutant (**C**), or wild-type Col-0 plants (**D**). [*t-test*, \**P* < 0.05; \*\**P* < 0.01; \*\*\**P* < 0.001; \*\*\*\**P* < 0.0001].

Finally, we generated transgenic lines that expressed two *myosin VIII* cDNAs (*VIII-A* and *VIII-B* [At2375], or *VIII-2* and *VIII-B* [At2376]) in both the *myosin VIII-1/2/a/b* quadruple mutant (Figure 2C) and in wild-type Col-0 plants (Figure 2D). Overexpression of these two sets of myosin cDNAs did not increase transformation of otherwise wild-type plants. However, expression of these pairs of myosin cDNAs in the *myosin VIII-1/2/a/b* quadruple mutant enhanced transformation to a greater extent than did expression of these cDNAs individually (Figures 2C and Supplemental Figures S5A-D). These results indicate that *myosin VIII-A*, *VIII-B*, and *VIII-2* genes contribute synergistically to transformation. These results further suggest that the level of expression of the *myosin VIII-2*, *VIII-A*, and *VIII-B* genes in wild-type roots is sufficient for maximal transformation, and that increasing expression of these genes cannot further increase transformation.

### Inducible expression of *myosin VIII*, but not *myosin XI-K*, cargo binding domain transgenes affects *Agrobacterium*-mediated transient root transformation

Yang et al. (2017) demonstrated that expression of a myosin XI-K cargo binding (globular tail) domain inhibited VirE2 movement in tobacco leaf cells. To determine whether specific myosin VIII members are important for *Agrobacterium*-mediated transformation using a similar dominant-negative approach, we generated transgenic *Arabidopsis* lines individually expressing cDNAs encoding each of the four myosin VIII cargo binding domains (CBD; At2315-2318), or the myosin XI-K CBD (At2319), under the control of a β-estradiol inducible promoter. Because these CBD constructs lack the myosin motor domain, they are not able to transport their cargoes along actin filaments. We tested the susceptibility of 3-4 independent transgenic lines for each CBD construct to *Agrobacterium*-mediated transient transformation, either after 24 hr transgene induction or without induction. Induction of each of the four *myosin VIII* CBD transgenes decreased transformation two- to three-fold, whereas induction of the *myosin XI-K* CBD transgene had no significant effect on transformation (Figure 3 and Supplemental Figures S6A-E). These data further suggest that the four myosin VIII proteins contribute to *Agrobacterium*-mediated transformation, whereas myosin XI-K is not required for transformation. However, below we show that myosin XI-K is important for VirE2 movement through the cytoplasm.

**Figure 3.**
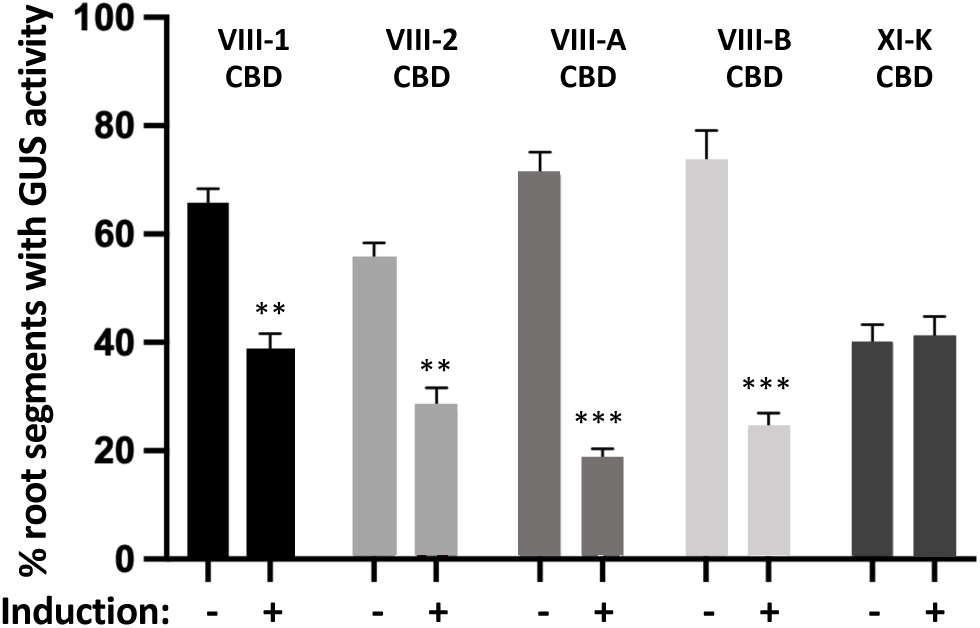
Inducible expression of myosin VIII, but not the myosin XI-K, cargo binding domains (CBD) inhibits transient transformation. Root segments from transgenic *Arabidopsis* expressing the cargo binding domain (CBD) of the indicated myosin protein, under the control of a β-estradiol inducible promoter, were treated with β-estradiol or control solutions for 24 hr before infection with *A. tumefaciens* At849. The percentage of roots showing GUS activity six days after infection indicates the transient transformation efficiency. A total of 10-15 plants and >100 segments per plant, from five independent transgenic lines, were tested for transient transformation. Values given are means ± SE. Asterisks indicate significant differences compared to uninduced plants. [*t-test*, \*\**P* < 0.01; \*\*\**P* < 0.001].

### Myosin VIII proteins are required for *Arabidopsis* roots expressing VirE2 to complement the virulence of a *virE2* mutant *Agrobacterium* strain, and to mitigate VirE2 aggregation

*Agrobacterium virE2* mutants are severely debilitated in virulence (Rossi et al., 1996). However, expression of VirE2 or VirE2-Venus transgenes in plants restores virulence, indicating that the major role of VirE2 in transformation occurs in the plant (Citovsky et al., 1992; Lapham et al., 2021). To determine the importance of myosin VIII proteins for such “*in planta*” complementation, we generated numerous *Arabidopsis* transgenic lines expressing a *VirE2-Venus* transgene, under the control of a β-estradiol-inducible promoter, in both wild-type and in *myosin VIII* quadruple mutant plants. Twenty four hr after incubation of roots of these lines in medium with or without inducer, we cut the roots into small segments and infected them with *A. tumefaciens* At1879 (lacking a *virE2* gene) containing the T-DNA binary vector pBISN2 with a *gusA*-intron gene in the T-DNA (Narasimhulu et al., 1996). Six days later we assessed transient transformation efficiency by conducting β-glucuronidase (GUS) activity assays using the chromogenic substrate 5-bromo-4-chloro-3-indolyl-β-D-glucuronide (X-gluc). As previously reported (Lapham et al., 2021), induction of the *VirE2-Venus* transgene in wild-type Col-0 plants complemented the loss of VirE2 function when a *virE2 Agrobacterium* mutant was used. However, induction of the *VirE2-Venus* transgene in *myosin VIII* quadruple mutant plants could not compensate the loss of VirE2 function when a *virE2 Agrobacterium* mutant was used (Figures 4A and 4B). Thus, *Myosin VIII* expression is required for *in planta* complementation of an *Agrobacterium virE2* mutant.

**Figure 4.**
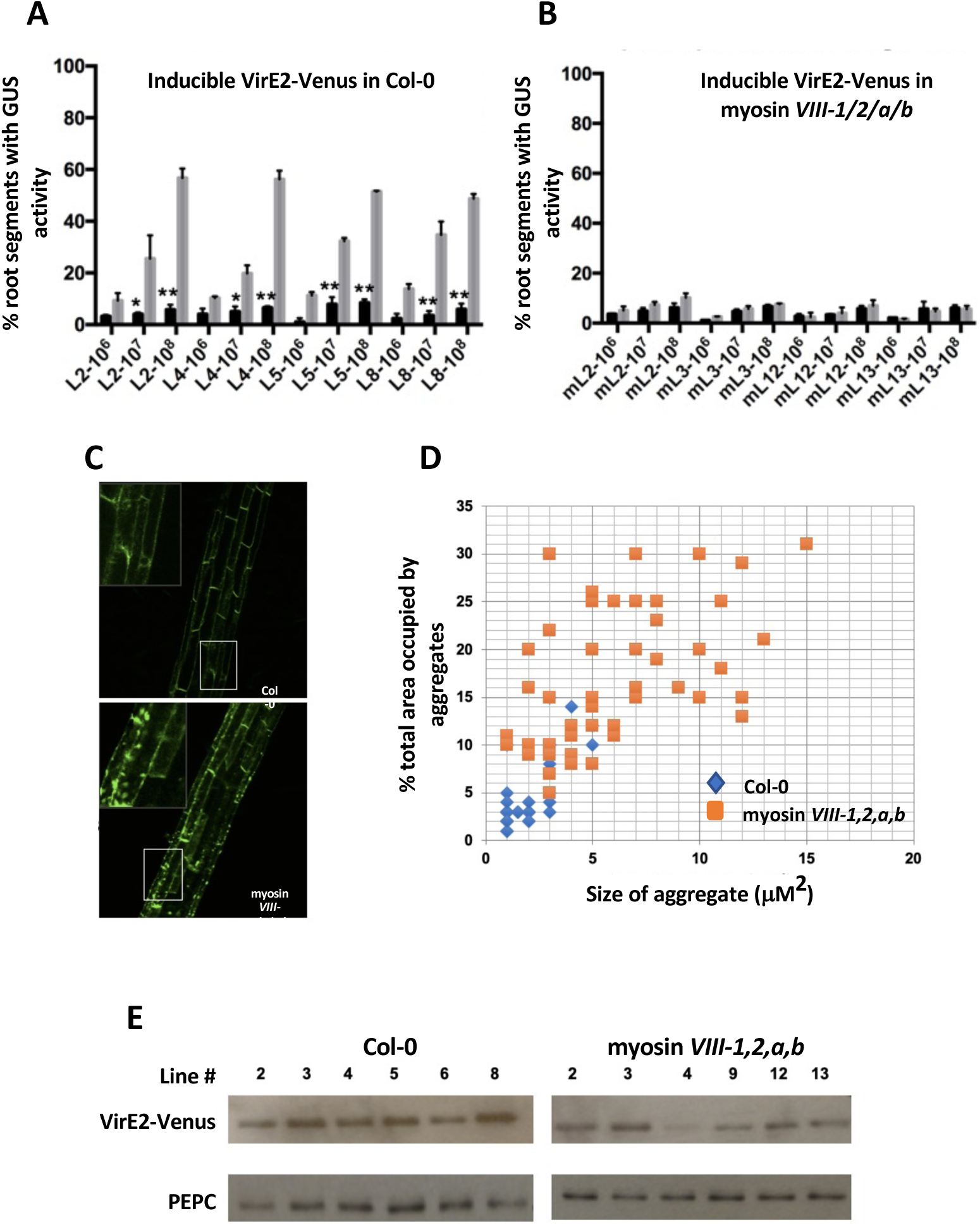
Expression of VirE2-Venus in the *myosin VIII-1/2/a/b* mutant cannot complement a *virE2* mutant *Agrobacterium* strain for transient transformation. **(A)** *Agrobacterium*-mediated transient transformation assays were conducted on roots of four independent transgenic lines of wild-type (Lines 2, 4, 5 and 8) or **(B)** the *myosin VIII-1/2/a/b* mutant (Lines 2, 3, 12, and 13). Root segments were inoculated with 10^6^, 10^7^, or 10^8^ cfu/ml of the *virE2* mutant strain *A. tumefaciens* At1879 containing the T-DNA binary vector pBISN2. Plants were treated with β-estradiol (gray bars) or control solution (black bars) for VirE2-Venus expression 24 hr before infection. The percentage of roots showing GUS activity was calculated as in Figure 1. A total of 10-15 plants from each line and >100 segments per plant were tested for transient transformation. Values given are means ± SE. Asterisks indicate significant differences compared to uninduced plants. [*t-test*, \**P* < 0.05; \*\**P* < 0.01]. **(C)** Confocal images showing aggregation of VirE2-Venus proteins in transgenic roots of either wild-type (top panel) or *myosin VIII 1/2/a/b* mutant backgrounds (bottom panel). **(D)** Quantitative analysis of the size of VirE2-Venus aggregates and the percentage of the cellular area occupied by the aggregates. Image J was used for analysis. The average VirE2-Venus aggregate size was 2.0±0.3 μm^2^ in wild-type roots, and 6.2±0.6 μm^2^ in *myosin VIII-1/2/a/b* quadruple mutant roots. **(E)** Western blot detection of VirE2-Venus proteins expressed in transgenic plants of either the Col-0 or the *myosin VIII-1/2/a/b* quadruple myosin mutant background. Mouse anti-GFP antibody was used to detect VirE2-Venus protein expressed after induction of individual transgenic lines. The house-keeping protein phosphoenolpyruvate carboxylase (PEPC) was detected using a rabbit anti-PEPC antibody and served as an internal control.

VirE2 usually localizes in the plant cytoplasm, although in the presence of the virulence effector protein VirD2 and T-strands a small amount may enter the nucleus (Gelvin, 2010; Li et al., 2020). To determine whether myosin VIII affects the subcellular distribution of VirE2, we used confocal microscopy to visualize β-estradiol induced VirE2-Venus in both the Col-0 and *myosin VIII* quadruple mutant backgrounds containing an inducible *VirE2-Venus* transgene. VirE2 is known to aggregate both *in vitro* and in plants (Bhattacharjee et al., 2008; Dym et al., 2008). Under our induction conditions VirE2-Venus showed a pattern of aggregation in the *myosin VIII* quadruple mutant distinct from that seen in the Col-0 background. In roots of the quadruple mutant, VirE2-Venus aggregated to a greater extent than it did in wild-type roots (Figure 4C). The sizes of VirE2 aggregates from two random areas of each of six images were measured and the average sizes were calculated. The average VirE2 aggregate size in root cells of the *myosin VIII* quadruple mutant was 6.2±0.6 μm^2^, compared to an average aggregate size of 2.0±0.3 μm^2^ in wild-type roots (Figure 4D). Furthermore, the total cellular area occupied by these VirE2 aggregates was greater in root cells of the *myosin VIII* mutant (17.0±1.1%) than in similar wild-type cells (4.7±0.9%) (Figure 4D). To determine whether this extensive VirE2 aggregation in myosin *VIII* mutant cells resulted from higher levels of VirE2-Venus expression in roots, we extracted total proteins from β-estradiol-induced roots of these plants and determined the relative amounts of VirE2-Venus expressed in roots using anti-GFP antibodies. As a loading control for the protein blots, we used an antibody targeting the house-keeping protein phosphoenolpyruvate carboxylase (PEPC). Strikingly, myosin *VIII* quadruple mutant roots express, on average, less VirE2-Venus than do wild-type roots (Figure 4E). Therefore, the more extensive aggregation of VirE2-Venus in *myosin VIII* mutant roots is unique and cannot be explained by greater accumulation of this protein.

To determine which myosin protein influences VirE2-Venus aggregation, we generated numerous transgenic *Arabidopsis* lines (using *Agrobacterium* strains At2315-At2319), in the *myosin VIII* quadruple mutant background, expressing both VirE2-Venus and individual *mCherry-myosin VIII CBD* cDNAs under the control of a β-estradiol-inducible promoter. Following induction, we measured the size of the VirE2-Venus aggregates in root cells and the percentage of root cell volume taken up by these aggregates. Figures 5A and 5B show that, compared to expression of VirE2-Venus in the absence of expression of a myosin CBD transgene, expression of myosin VIII-A, VIII-B, and VIII-2 CBDs reduced both the size of VirE2-Venus aggregates (from 7.5±1.9 μm^2^ down to 2.4±0.3 μm^2^, 3.7±0.5 μm^2^, and 3.1±0.3 μm^2^, respectively) and the percentage of the cell volume (from 17.7±2.0% down to 3.6±0.6%, 6.3±0.4%, and 6.1±0.5%, respectively) occupied by these aggregates, due to their dominant negative effects. However, the CBD of myosin VIII-1 did not substantially reduce aggregate size or the percentage of the cell volume occupied by the aggregates (Figure 5A): the average aggregates size was 6.7±0.7 μm^2^, and the percentage of the cell volume was 11.8±1.1%. These results suggest that myosin VIII-A, VIII-B, and VIII-2 CBDs interact with VirE2 in plant cells and that the myosin VIII-1 CBD may fundamentally differ from the CBDs of the other three myosin VIII members.

**Figure 5.**
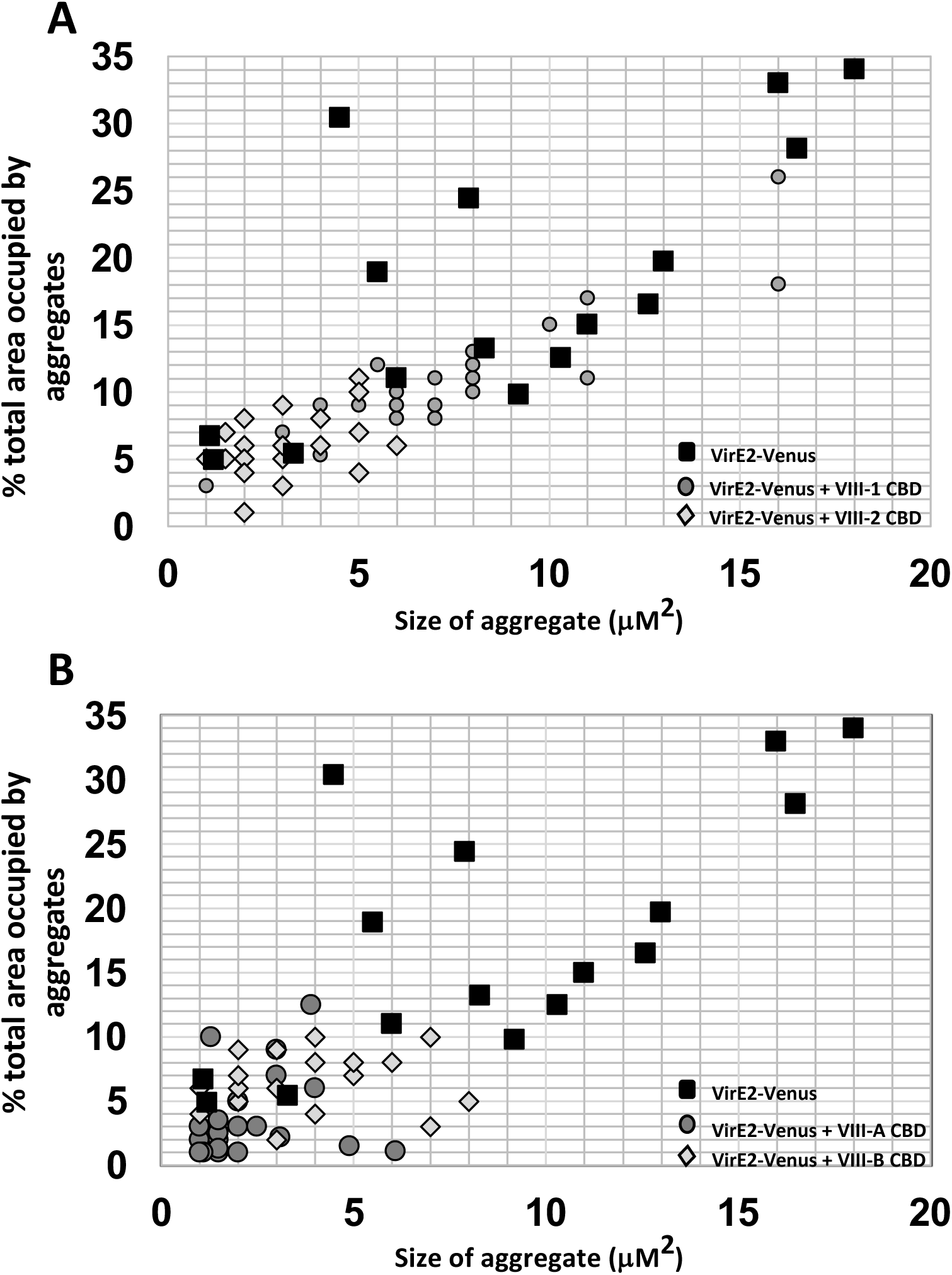
Analysis of VirE2-Venus aggregates in roots of transgenic *Arabidopsis* expressing the CBDs of various Myosin VIII proteins. Roots were treated with β-estradiol for eight hr. Random areas of confocal images from roots expressing **(A)** Myosin VIII-1 or VIII-2 CBDs, or **(B)** Myosin VIII-A or VIII-B CBDs were analyzed using Image J. Both the sizes and the percentage of the cellular area occupied by the VirE2-Venus aggregates are shown.

### VirE2 interacts directly with some, but not all, myosin VIII cargo binding domains and with the CBD of myosin XI-K

Because myosin VIII proteins are required for VirE2 to function in virulence, and because myosin VIII CBD expression affects the degree to which VirE2 aggregates *in planta*, we investigated whether myosin protein cargo binding domains interact with VirE2. We first tested the direct interaction of VirE2 with each of the myosin VIII and the myosin XI-K protein CBDs using an *in vitro* pull-down assay. We individually incubated recombinant VirE2-Venus with recombinant myc-tagged myosin CBDs. As a control, we substituted recombinant Venus protein for VirE2-Venus. As further controls, we incubated VirE2-Venus with myc-tagged VIP1, a protein known to interact with VirE2 (Tzfira et al., 2001; Djamei et al., 2007), or myc-tagged Lamin C, a protein known not to interact with VirE2 (Ueki and Citovsky, 2005). We captured VirE2-Venus, or Venus, and any interacting proteins on anti-GFP antibody beads, subjected the bound proteins to SDS gel electrophoresis, then conducted Western blot analysis using anti-myc antibodies. Figures 6A-F show that the CBDs of myosins VIII-A, VIII-B, VIII-2, and XI-K directly interacted with VirE2-Venus, but not with Venus. However, we were not able to detect interaction of VirE2-Venus with the CBD of myosin VIII-1. As expected, VirE2-Venus interacted with myc-tagged VIP1 but not with myc-tagged Lamin C (Figure 6). These data were confirmed using a yeast two-hybrid system. VirE2 interacted in yeast with the CBDs of myosins VIII-A, VIII-B, VIII-2, and XI-K, as well as with VIP1. However, VirE2 did not interact in yeast with the CBD of myosin VIII-1 or with the negative control Lamin C (Figures 7A and 7B).

**Figure 6.**
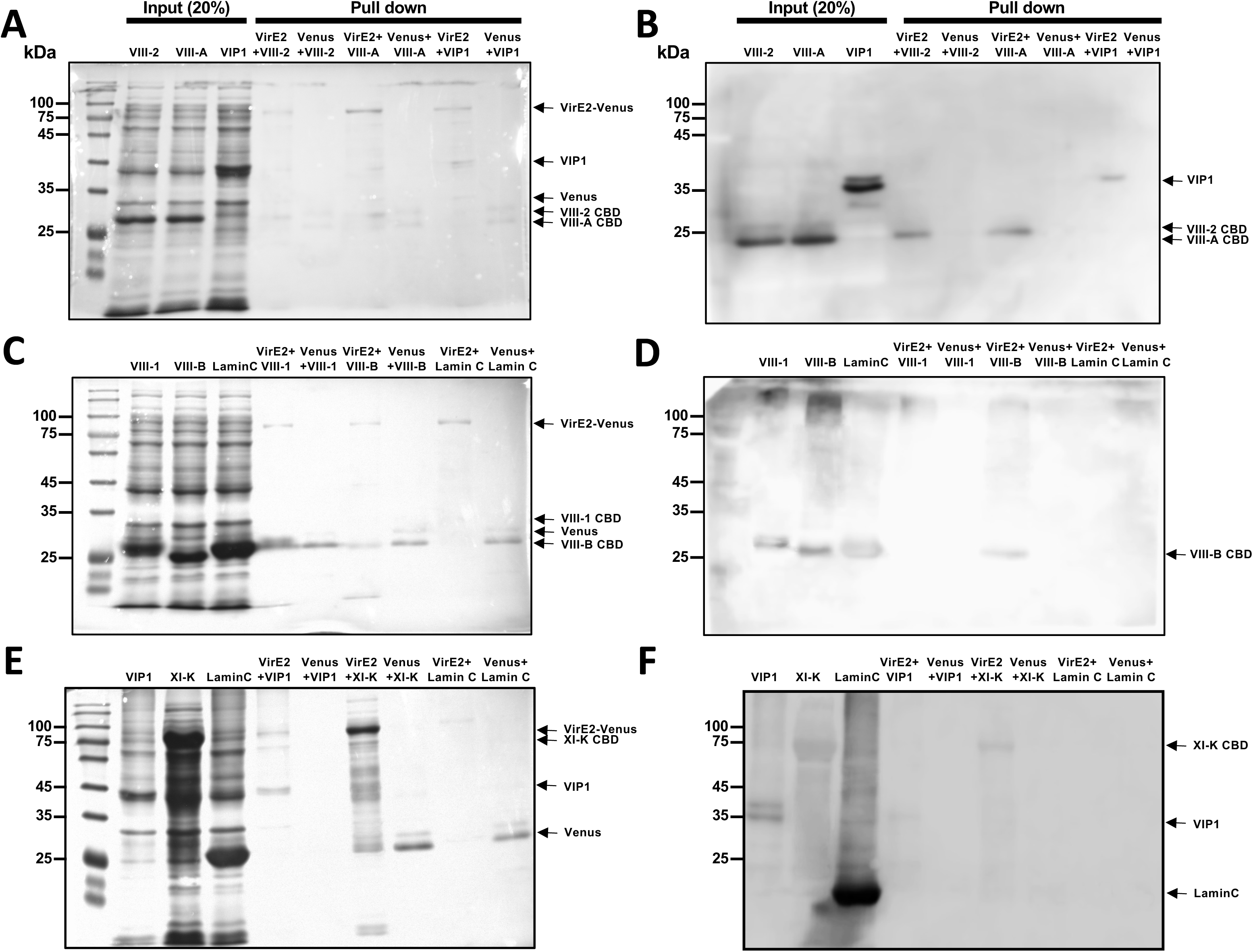
VirE2-Venus interacts with some myosin cargo binding domains (CBDs) *in vitro*. VirE2-Venus or Venus proteins were incubated with the indicated myc-tagged myosin CBD, VIP1, or Lamin C. Following binding to anti-GFP beads, proteins were eluted and subjected to Western blot analysis using anti-myc antibodies. Coomassie stained gels are shown in panels **A**, **C**, and **E**, and immunoblots in panels **B**, **D**, and **F**. VirE2-Venus interacts with the CBDs of myosins VIII-2, VIII-A, VIII-B, and XI-K, and with VIP1 (panels **B**, **D**, and **F**), but not with the CBD of myosin VIII-1 or with Lamin C (panels **D** and **F**). Venus, as a negative control, does not interact with any myc-tagged protein.

**Figure 7.**
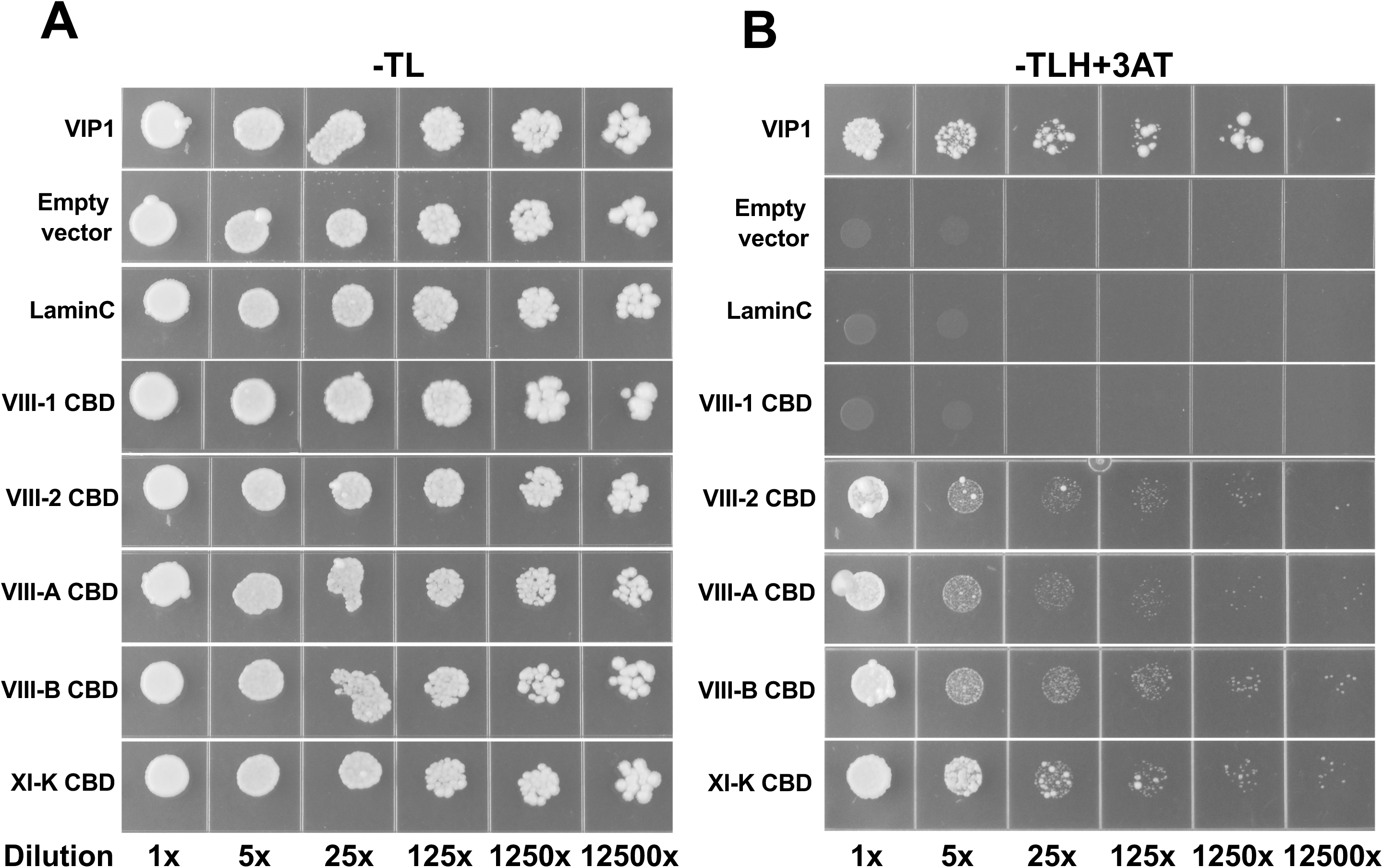
The cargo binding domains (CBD) of some myosins interact with VirE2 in yeast. Yeast two-hybrid analysis was conducted with VirE2 as the bait and individual myosin CBDs as the prey. Following co-transformation of the bait and prey plasmids, single colonies were picked and serial dilutions of yeast were grown on **(A)** SD-Trp-Leu (-TL) medium, or **(B)** SD-Trp-Leu-His medium supplemented with 3 mM 3-amino-1,2,4-triazole (-TLH+3AT). Colony growth on SD-TLH+3AT indicates interaction between VirE2 and the myosin CBD. VIP1 was used as a positive control for interaction with VirE2. Lamin C and the empty vector were used as negative controls. SD, synthetic dropout medium.

To assess the interactions of VirE2 with myosin VIII or XI-K CBD proteins *in vivo*, we conducted co-immunoprecipitation experiments using transgenic *Arabidopsis* roots expressing VirE2-Venus and, individually, each of the myc-tagged myosin VIII or the myosin XI-K CBDs. Using anti-GFP antibody beads to capture VirE2-Venus and associated proteins followed by immunoblotting using anti-myc tag antibody to detect CBDs, we observed robust co-immunoprecipitation between VirE2-Venus and the myosin VIII-2, VIII-A, VIII-B, and XI-K CBDs (∼4% of the input myc-myosin VIII-2, VIII-A, VIII-B, and XI-K CBDs were co-immunoprecipitated with anti-GFP antibody). However, we also observed some co-immunoprecipitation between VirE2 and the myc-tagged myosin VIII-1 CBD (∼1.6-2% of the input myosin VIII-1 CBD; Figures 8A and 8B). As expected, VirE2-Venus interacted with VIP1 (Figure 8A), and the myosin VIII-2 CBD did not interact with the control Venus protein (Figure 8B). The *in vivo* co-immunoprecipitation of VirE2-Venus with the myosin VIII-2, VIII-A, VIII-B, and XI-K CBDs recapitulated the *in vitro* interaction and yeast two-hybrid results. However, *in vivo* co-immunoprecipitation of the myosin VIII-1 CBD with VirE2-Venus, in the absence of interaction *in vitro* and in yeast, suggests that this *in vivo* interaction may be indirect and mediated by other factors within *Arabidopsis* roots.

**Figure 8.**
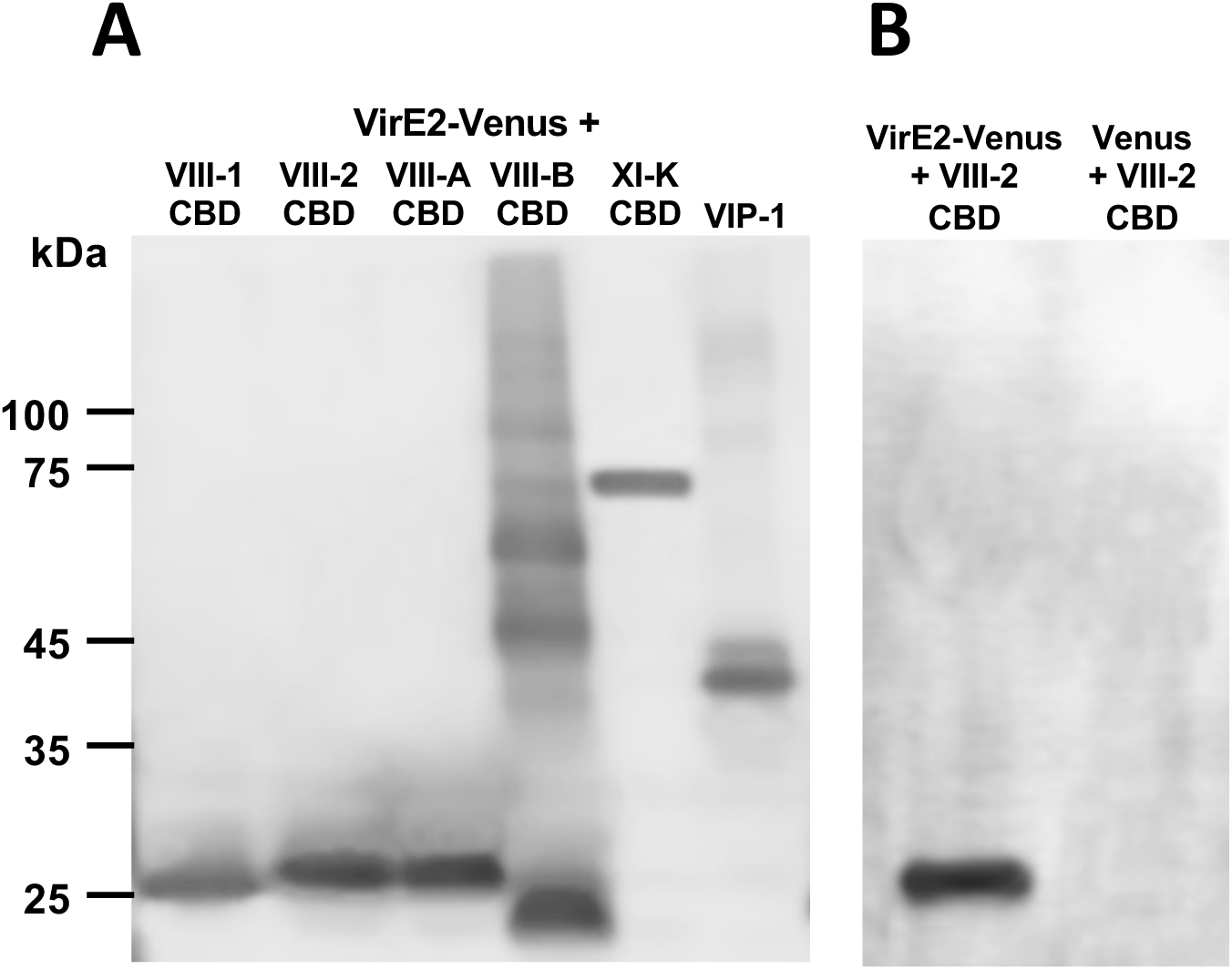
Myosin cargo binding domains can form complexes with VirE2 when expressed in *Arabidopsis.* **(A)** Protein extracts from roots of transgenic *Arabidopsis* plants expressing VirE2-Venus, or Venus, and the indicated myc-tagged myosin CBD transgenes were incubated with anti-GFP beads, the bound proteins eluted, and subjected to Western blot analysis using anti-myc antibodies. Myc-tagged VIP1 was used as a positive control. **(B)** Root extracts expressing VirE2-Venus or Venus and myc-tagged myosin VIII-2 CBD were incubated with anti-GFP beads, the bound proteins eluted, and subjected to Western blot analysis using anti-myc antibodies.

### VirE2 and Myosin VIII proteins co-localize at the cell periphery

The vast majority of VirE2 protein localizes within the cellular cytoplasm, and in wild-type *Arabidopsis* roots is often found in the peripheral regions of the cell (Shi et al., 2014). VirE2 has been proposed to form membrane channels through which VirD2/T-strands translocate from *Agrobacterium* into the plant cell (Dumas et al., 2001; Duckely et al., 2005). Myosin VIII proteins also localize to the plasma membrane (Golomb et al., 2008). To determine whether VirE2 and full-length myosin proteins co-localize at the cell periphery, we generated transgenic *Arabidopsis* plants expressing VirE2-Venus and various mCherry-tagged full-length myosin protein cDNAs under the control of a β-estradiol inducible promoter (using *Agrobacterium* strains At2447-At2450) in the Col-0 background. After β-estradiol induction, we imaged roots using confocal microscopy (Figure 9). Myosins VIII-1, VIII-A, and VIII-B localized mostly to the cellular periphery, whereas myosin VIII-2 localized both to the cellular periphery and to the nucleus. All myosin VIII and XI-K proteins also localized throughout the cytoplasm.

**Figure 9.**
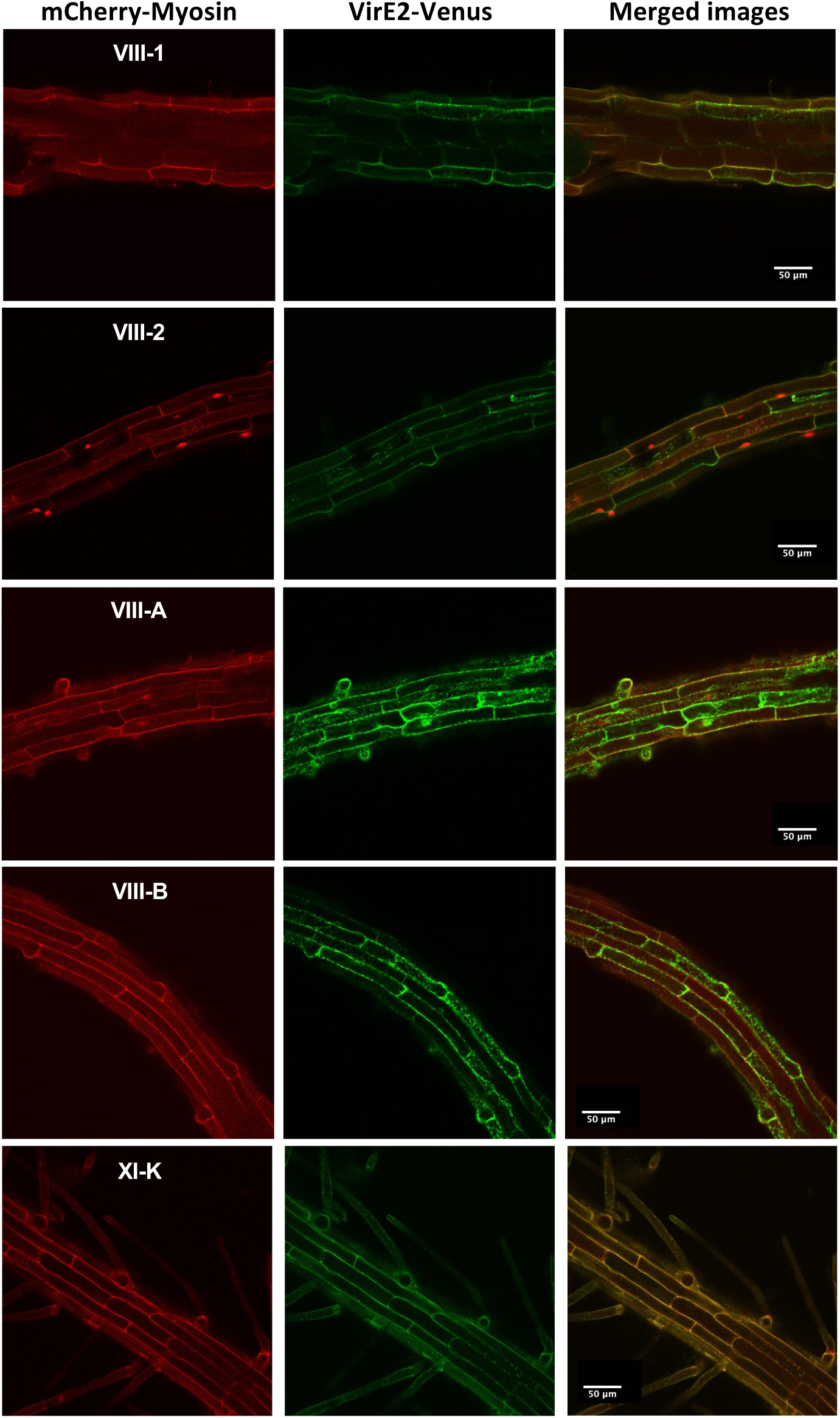
Co-localization of VirE2-Venus and mCherry-Myosins in *Arabidopsis* roots. Roots of transgenic *Arabidopsis* plants containing VirE2-Venus and mCherry-myosin genes, each under the control of a β-estradiol-inducible promoter, were treated with β-estradiol for 8 hr and examined by confocal microscopy. Bars indicate 50 μm.

We also determined whether VirE2 and myosin cargo binding domains co-localize at the cell periphery. We similarly generated transgenic *Arabidopsis* plants expressing VirE2-Venus and various mCherry-tagged myosin CBD cDNAs under the control of a β-estradiol inducible promoter (using *Agrobacterium* strains At2315-At2319). After β-estradiol induction, we imaged roots using confocal microscopy (Figures S10A and S10B). The CBDs of myosins VIII-A and VIII-2 localized almost exclusively to the cellular periphery, whereas the CBD of myosin VIII-B localized both to the cellular periphery and to the nucleus (Figure S10A). The myosin VIII-1 and XI-K CBDs localized not only at the cellular periphery but also throughout the cytoplasm (Figure S10A).

When co-expressed in the Col-0 background, VirE2 and the myosin VIII-2, VIII-A, VIII-B, and XI-K CBDs partially co-localized at the cell periphery (Pearson’s correlation coefficient = 0.56±0.04, 0.68±0.05, 0.63±0.04, and 0.69±0.03, respectively; overlap coefficient = 0.65±0.03, 0.72±0.06, 0.70±0.03, and 0.77±0.04, respectively; Figure S10B). However, co-localization of VirE2 with the myosin VIII-1 CBD at the cell periphery was less extensive (Pearson’s correlation coefficient = 0.44±0.03; overlap coefficient = 0.48±0.01; Figure S10B). Staining VirE2-Venus expressing *Arabidopsis* roots with the plasma membrane dye FM4-64 revealed that VirE2 localized near to but not in the plasma membrane (Pearson’s correlation coefficient = 0.42±0.07; overlap coefficient = 0.45±0.03; Supplemental Figures S7A and B). However, in the myosin VIII quadruple mutant background, VirE2 localized less extensively at the cellular periphery and more within the general cytoplasm (Supplemental Figures S8A and B). Co-expression of mCherry-myosin VIII-1, VIII-A, or VIII-2 CBDs with the plasma membrane marker PIP2A-Venus (using *Agrobacterium* strains At2370, At2371, and At2374, respectively) revealed that these myosin VIII CBDs co-localize with PIP2A (Supplemental Figures S9A-C), similar to what has previously been reported (Golomb et al., 2008).

Taken together, these results suggest that the myosin VIII-2, VIII-A, and VIII-B full-length proteins and their respective CBDs localize to the plasma membrane where they may temporarily tether VirE2.

### VirE2 and several myosin VIII full-length proteins interact at the cell periphery

In order to test whether VirE2 may be a cargo of myosin VIII proteins, we conducted bimolecular fluorescence complementation (BiFC) assays to visualize *in planta* protein-protein interactions. We generated transgenic *Arabidopsis* plants expressing VirE2-cYFP and nVenus-tagged full-length myosin protein cDNAs under the control of a β-estradiol inducible promoter (using *Agrobacterium* strains At2443-At2446). The plants also constitutively expressed mCherry-actin binding domain 2 (ABD2) as an actin marker. After inducing the plants with β-estradiol, we detected YFP fluorescence mainly at the cellular periphery of root cells when VirE2-cYFP was co-expressed with the nVenus-tagged myosins VIII-2, VIII-A, or VIII-B, but no fluorescence with myosin VIII-1 (Figure 10), indicating a lack of direct interaction with the myosin VIII-1 protein. These results again suggest that the CBDs of myosin VIII-2, VIII-A, and VIII-B may temporarily tether VirE2 to the plasma membrane.

**Figure 10.**
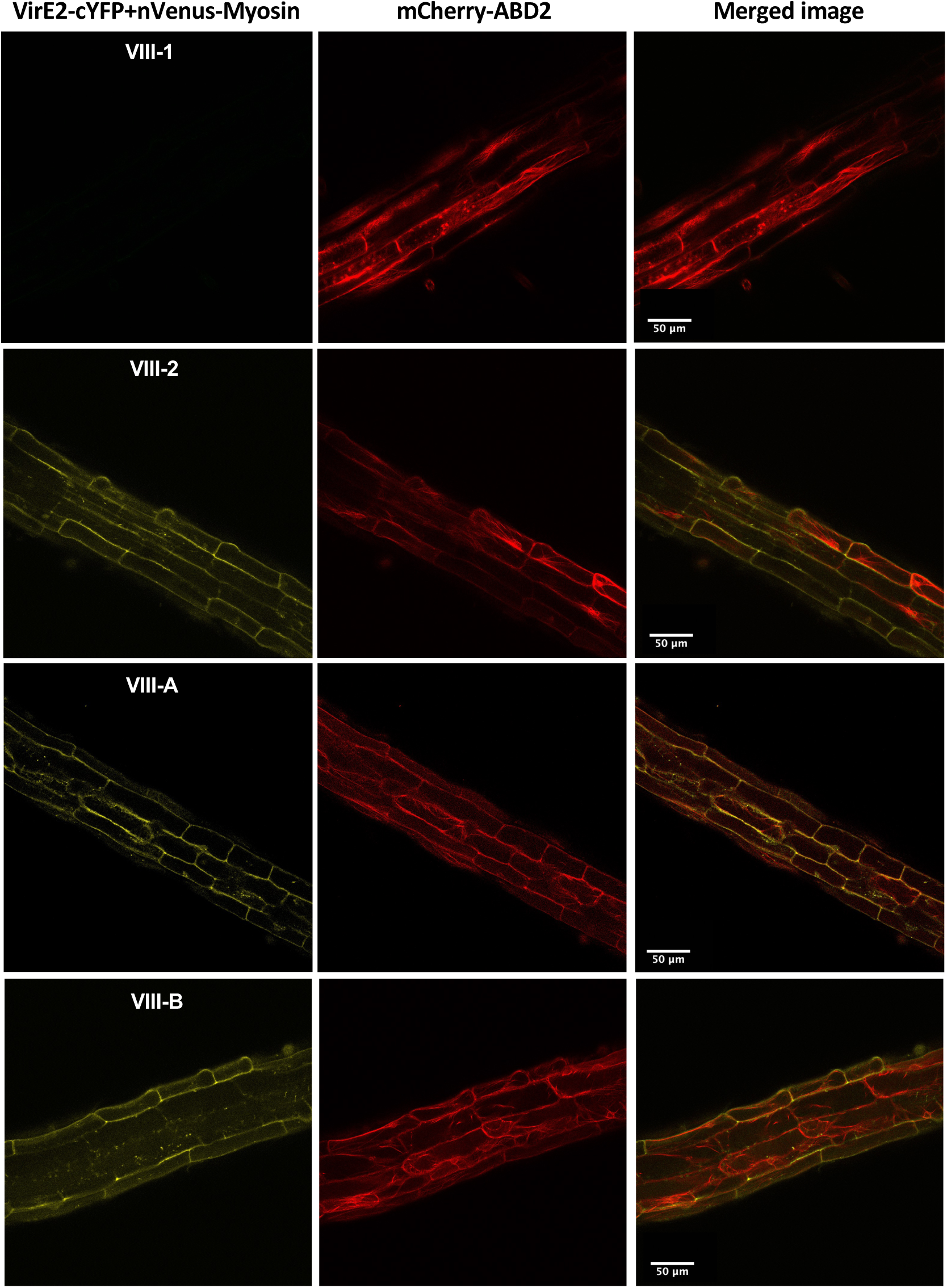
Bimolecular fluorescence complementation (BiFC) of VirE2 with various myosin full-length proteins in *Arabidopsis* roots. Transgenic *Arabidopsis* plants containing VirE2-cEYFP and the indicated myosin-nVenus constructs, each under the control of a β-estradiol-inducible promoter, as well as mCherry-ABD2 were treated with β-estradiol for 48 hr and the fluorescence signal detected by confocal microscopy. VirE2 interacts with the CBDs of myosins VIII-2, VIII-A, and VIII-B. VirE2 does not interact with the myosin VIII-1. Yellow fluorescence indicates interaction of VirE2-Venus with the myosin CBD. Actin filaments are labeled by mCherry-ABD2. Bars indicate 50 μm.

We similarly conducted bimolecular fluorescence complementation (BiFC) assays to visualize *in planta* protein-protein interactions between VirE2 and myosin cargo binding domains. We generated transgenic *Arabidopsis* plants expressing VirE2-cYFP and nVenus-tagged myosin CBD cDNAs under the control of a β-estradiol inducible promoter (using *Agrobacterium* strains At2320-At2324). The plants also constitutively expressed mCherry-actin binding domain 2 (ABD2) as an actin marker. After inducing the plants with β-estradiol, we detected YFP fluorescence mainly at the cellular periphery of root cells when VirE2-cYFP was co-expressed with the nVenus-tagged CBDs of myosins VIII-A, VIII-B, VIII-2, or myosin XI-K, but no fluorescence with the myosin VIII-1 CBD (Figure S11), indicating a lack of direct interaction. VirE2-cYFP additionally interacted with the CBD of myosin XI-K in the cytoplasm. VirE2-cYFP did not interact with the negative control protein Lamin C. These results again suggest that the CBDs of myosin VIII-2, VIII-A, VIII-B, and XI-K may temporarily tether VirE2 to the plasma membrane, but that only myosin XI-K may transport VirE2 along actin filaments through the cytoplasm.

### Some VirE2 molecules re-localize to the perinuclear region during infection by a virulent *Agrobacterium strain*

Some VirE2-Venus fluorescence, when constitutively expressed in *Arabidopsis* roots, localizes at the cellular periphery, especially at the cell poles (Shi et al., 2014; Li et al., 2020). Li et al. (2020) showed that during agroinfiltraion, a small amount of VirE2 can enter the nucleus of tobacco leaf cells, and this entry requires the presence of VirD2 and T-strands. To determine whether transgenically-expressed VirE2 also relocalizes to the nucleus in the presence of VirD2 and T-strands, we incubated *Arabidopsis* roots containing an inducible *VirE2-Venus* transgene with β-estradiol. After 24 hr, we infected root segments with a *virE2*-deletion *Agrobacterium* strain either lacking T-DNA (At2403; no binary vector) or containing T-DNA on a binary vector (At2404). Twenty-four hours after inoculation, we observed VirE2-Venus subcellular localization by confocal microscopy. Root segments infected by *A. tumefaciens* At2403 did not show altered VirE2-Venus subcellular localization: fluorescence was distributed throughout the cytoplasm, including at the cellular periphery (Supplemental Figure S12A). However, infection by *A. tumefaciens* At2404 resulted in a partial redistribution of VirE2-Venus within the cytoplasm: fluorescence increased in the perinuclear area (Supplemental Figure S12B). Quantitative analysis of fluorescence intensity showed that perinuclear VirE2 increased ∼5-fold in the presence of T-DNA (Figure 11A). However, we were unable to detect VirE2-Venus within the nucleus. These data suggest that the transfer of T-DNA to plant cells results in a partial redistribution of VirE2 to the area surrounding the nucleus.

**Figure 11.**
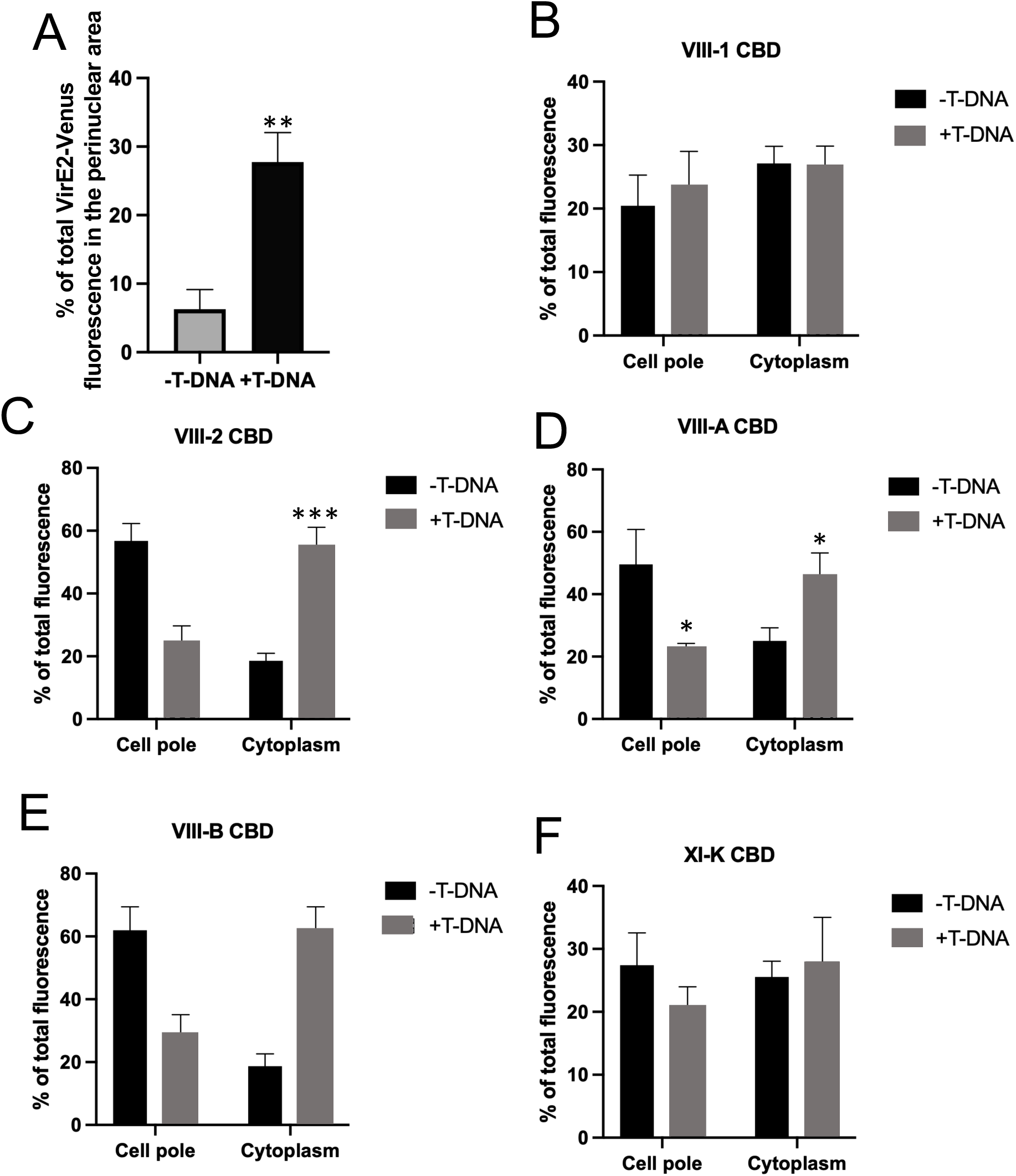
Incubation of *Arabidopsis* roots with *Agrobacterium* that can transfer T-DNA results in increased VirE2-Venus in the perinuclear region. **A)** Transgenic *Arabidopsis* plants containing an inducible *VirE2-Venus* transgene were treated with 5 μm β-estradiol for 24 hr, followed by inoculation with 10^8^ cfu/ml of the *virE2* mutant *A. tumefaciens* At1872 without a binary vector (At2403, left panel) or with T-DNA on a binary vector (At2404, right panel). The percentage of total VirE2-Venus fluorescence in the perinuclear area was calculated. Images of 10-15 transgenic lines and >50 cells were analyzed. The data indicate means ± SE. Asterisks indicate significant differences between roots inoculated with the two *Agrobacterium* strains. **(B-F)** the relative distribution of VirE2-Venus at the cell poles or in the cytoplasm following infection with *A. tumefaciens* At2403 or At2404 of roots co-expressing CBDs of myosin VIiI-1. myosin VIII-2, myosin VIII-A, myosin VIII-B, or myosin XI-K (B, C, D, E, F, respectively). Images of 10-15 transgenic lines and >50 cells were analyzed. The data indicate means ± SE. Asterisks indicate significant differences between roots inoculated with the two *Agrobacterium* strains. [*t-test*, \**P* < 0.05; \*\**P* < 0.01; \*\*\**P* < 0.001].

### T-strands delivered by *Agrobacterium* can relocalize VirE2 from the cellular periphery into the cytosol

Because much VirE2 produced *in planta* is already in the general cell cytosol, the data presented in Figure 11A and the movies in Supplemental Figure S13 were not able to show redistribution of VirE2-Venus specifically from the cell periphery into the perinuclear area. We therefore conducted the VirE2-Venus re-localization experiments, following *Agrobacterium* infection, on root segments overexpressing the various myosin VIII or myosin XI-K CBDs. We reasoned that when most VirE2-Venus is tethered to the plasma membrane by myosin VIII-A, VIII-B, VIII-2, and (to a lesser extent) XI-K, as seen in Supplemental Figure 10A, we may be able to detect re-distribution to VirE2-Venus more easily when T-strands are introduced into the cells by *Agrobacterium*.

We generated transgenic plants expressing VirE2-Venus and individual myc-tagged myosin CBD cDNAs (using *Agrobacterium* strains At2423-At2427), both under the control of a β-estradiol-inducible promoter. After inducing the transgenes for 24 hr, we infected root segments with *virE2* mutant *Agrobacterium* strains lacking or containing a T-DNA binary vector. The bacteria (At2403 and At2404, respectively) additionally contained a plasmid expressing mCherry to mark the location of the bacteria during infection. We observed fluorescence by confocal microscopy eight hours after infection and calculated the percentage of fluorescence shift between the cell periphery and the cytosol. When At2404, capable of transferring T-DNA, was used, some of the VirE2-Venus fluorescence signal re-localized from the cell periphery into the cytoplasm when plants expressed the myosin VIII-A, VIII-B, or VIII-2 CBDs (Figures 11C, 11D, and 11E, respectively, and movies in Supplemental Figures S15-17). No such redistribution of the VirE2-Venus signal was observed when we infected roots with a strain, At2403, lacking T-DNA. The distribution of VirE2-Venus signal was not altered in roots infected with *A. tumefaciens* At2404 and expressing the CBDs of myosins VIII-1 or XI-K (Figures 11 B and F; movies in Supplemental Figures S14 and S18). These results further suggest that myosins VIII-A, VIII-B, and VIII-2 may temporarily tether VirE2 at the cell periphery. However, upon transfer of T-DNA into the cell, VirE2 is released into the cellular interior.

### Following T-DNA transfer, VirE2 moves from the cellular periphery to the interior along actin filaments

To address whether VirE2-Venus moves along actin filaments, we used *Arabidopsis* transgenic lines expressing inducible VirE2-Venus and myc-tagged myosin VIII CBD cDNAs, and expressing mCherry-ABD2 to mark actin filaments (using *Agrobacterium* strains At2365-At2369). Time-lapse imaging of control plants, which expressed VirE2-Venus and mCherry-ABD2 but without any myosin CBD, indicated that some VirE2-Venus moves along actin filaments at a velocity of 0.353±0.06 μm/sec (Figure 12). In the presence of any of the four myosin VIII CBDs, the velocity of VirE2-Venus along actin filaments in leaf or root cells remained unchanged (Table 1). Additionally, the velocity of VirE2-Venus movement along actin filaments was not altered in the *myosin VIII-1/2/a/b* quadruple mutant (Figure 12, Table 1). However, when the myosin XI-K CBD was expressed in the plant, no VirE2 movement was detected in either leaf or root cells. These results indicate that only myosin XI-K, but not myosin VIII proteins, are important for VirE2 trafficking through leaf and root cells.

**Figure 12.**
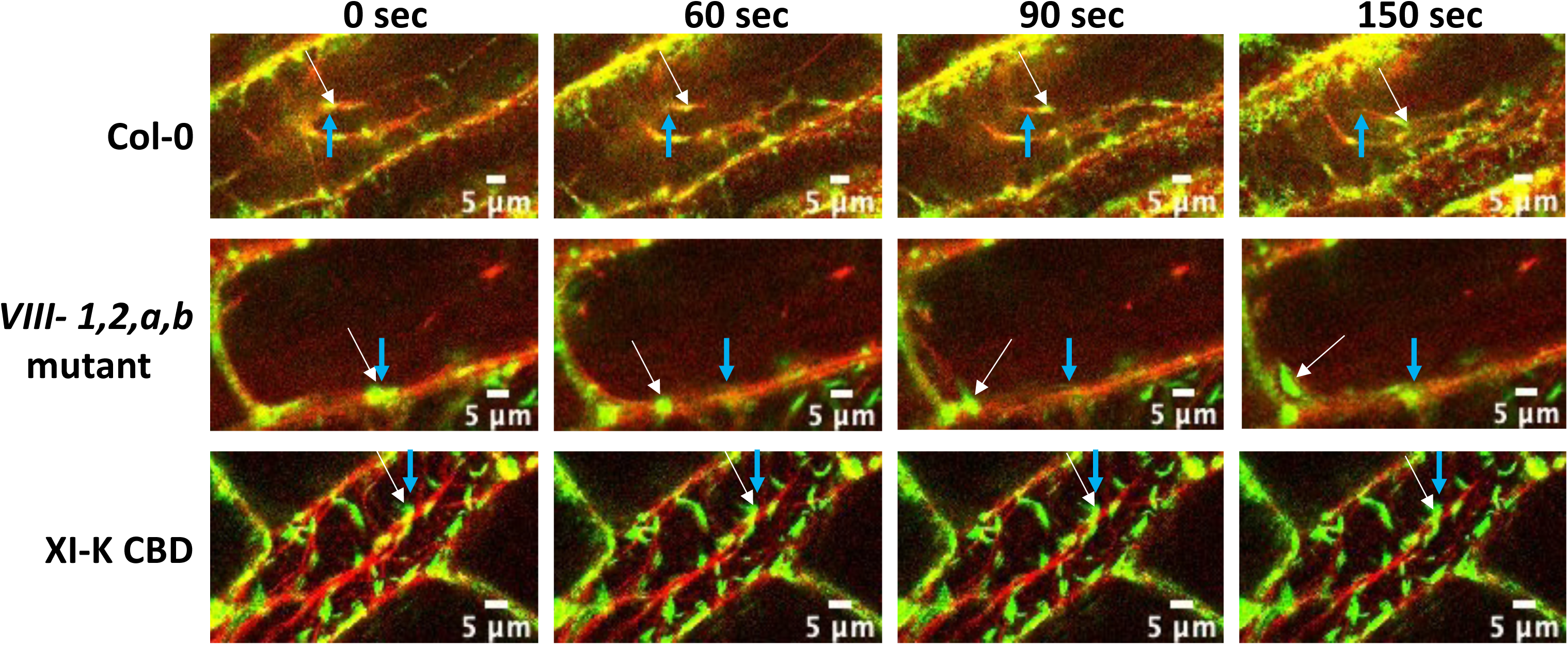
Loss of function of myosin XI-K, but not myosin VIII, disrupts movement of VirE2-Venus along actin filaments. Transgenic plants expressing mCherry-ABD2 constitutively and VirE2-Venus inducibly in Col-0, *myosin VIII-1/2/a/b*, and myosin XI-K CBD plants were used in this experiment. Roots of these plants were treated with 10 μm β-estradiol for 8 hr. Time lapse images were taken by confocal microscopy. Four images of each field are shown at the indicated times (0, 60, 90, and 120 seconds). Blue arrows indicate the initial position of a particular VirE2-Venus aggregate; white arrows indicate the position of the same aggregate at various times thereafter. Bars indicate 5 μm.

**Table 1.** Velocity of VirE2-Venus movement in leaf and root cells of various *Arabidopsis* transgenic lines.

| Transgene(s) in plant/<br>*Agrobacterium*<br>infection<sup>a</sup> | *Arabidopsis*<br>background | Velocity in leaves<br>($\mu\text{m}/\text{sec}$) | Velocity in roots<br>($\mu\text{m}/\text{sec}$) |
| --- | --- | --- | --- |
| VirE2-Venus | Col-0 | 0.353± 0.06 | 0.312± 0.07 |
| VirE2-Venus | myosin<br>*VIII-1/2/a/b* | 0.378± 0.02 | 0.318± 0.02 |
| VirE2-Venus +<br>Myosin VIII-1 CBD | Col-0 | 0.407 ± 0.06 | 0.341 ± 0.04 |
| VirE2-Venus +<br>Myosin VIII-2 CBD | Col-0 | 0.398 ± 0.07 | 0.321 ± 0.02 |
| VirE2-Venus +<br>Myosin VIII-A CBD | Col-0 | 0.387 ± 0.07 | 0.304 ± 0.01 |
| VirE2-Venus +<br>Myosin VIII-B CBD | Col-0 | 0.401 ± 0.05 | 0.328 ± 0.04 |
| VirE2-Venus +<br>Myosin XI-K CBD | Col-0 | Not detectable | Not detectable |
| VirE2-Venus +<br>Myosin XI-K CBD +<br>*Agrobacterium* (- T-DNA) | Col-0 | Not detectable | Not detectable |
| VirE2-Venus +<br>Myosin XI-K CBD +<br>*Agrobacterium* (+ T-DNA) | Col-0 | Not detectable | Not detectable |
| *Agrobacterium* VirE2-iGFP11<sup>b</sup> | Col-0 GFP1-10 | 0.323 ± 0.08 | ----- |

<sup>a</sup>All transgenes are under the control of a β-estradiol-inducible promoter; CBD, cargo binding domain. >50 cells of 10 independent transgenic lines were examined for each genotype of transgenic plant.
<sup>b</sup>Measurements were conducted on 19 VirE2 aggregates within 10 cells.

To test if the blockage of VirE2-Venus movement along actin filaments by the myosin XI-K CBD could be reversed by the presence of VirD2/T-strands, we infected roots with a *virE2* mutant *Agrobacterium* strain harboring a T-DNA with a mCherry-intron-NLS expression cassette (At2405). Time-lapse confocal microscopy still indicated no movement of VirE2-Venus in the mCherry expressing cells. (Table 1 and Supplemental Figure S19).

Taken together, our results indicate that myosin XI-K but not myosin VIII proteins are important for VirE2 intracellular trafficking. We suggest that myosin VIII proteins help tether VirE2 to the plasma membrane/peripheral regions of the cell. Upon introduction of VirD2/T-strands, VirE2 is released and is trafficked through the cell by myosin XI-K.

### VirE2 delivered from *Agrobacterium* into plant cells moves along filaments

The experiments conducted above measured VirE2 movement when the protein was expressed *in planta*. To determine if VirE2 movement also occurred when the protein is delivered from *Agrobacterium*, we adapted the split GFP system of Li and Pan (2017). We internally tagged VirE2 with GFP11 and used an *Agrobacterium* strain harboring this construct to infiltrate leaves of *Arabidopsis* expressing GFP1-10. Such an internally tagged VirE2 protein is active in transformation (Li and Pan, 2017). Forty eight hr post infiltration, we imaged the leaves using confocal microscopy. As a control, we imaged the leaves of uninfected GFP1-10 *Arabidopsis* plants. Similar to what we saw when VirE2-Venus was made in *Arabidopsis* root cells, VirE2 introduced into plant cells from *Agrobacterium* localized in the cytoplasm predominantly at the cellular periphery, but also along filaments traversing the cell (Figure 13). Time-lapse confocal microscopy of 19 VirE2 aggregates indicated that they moved along these filaments with a velocity of 0.323±0.08 μm/sec, similar to that of VirE2-Venus made *in planta* (Table 1).

**Figure 13.**
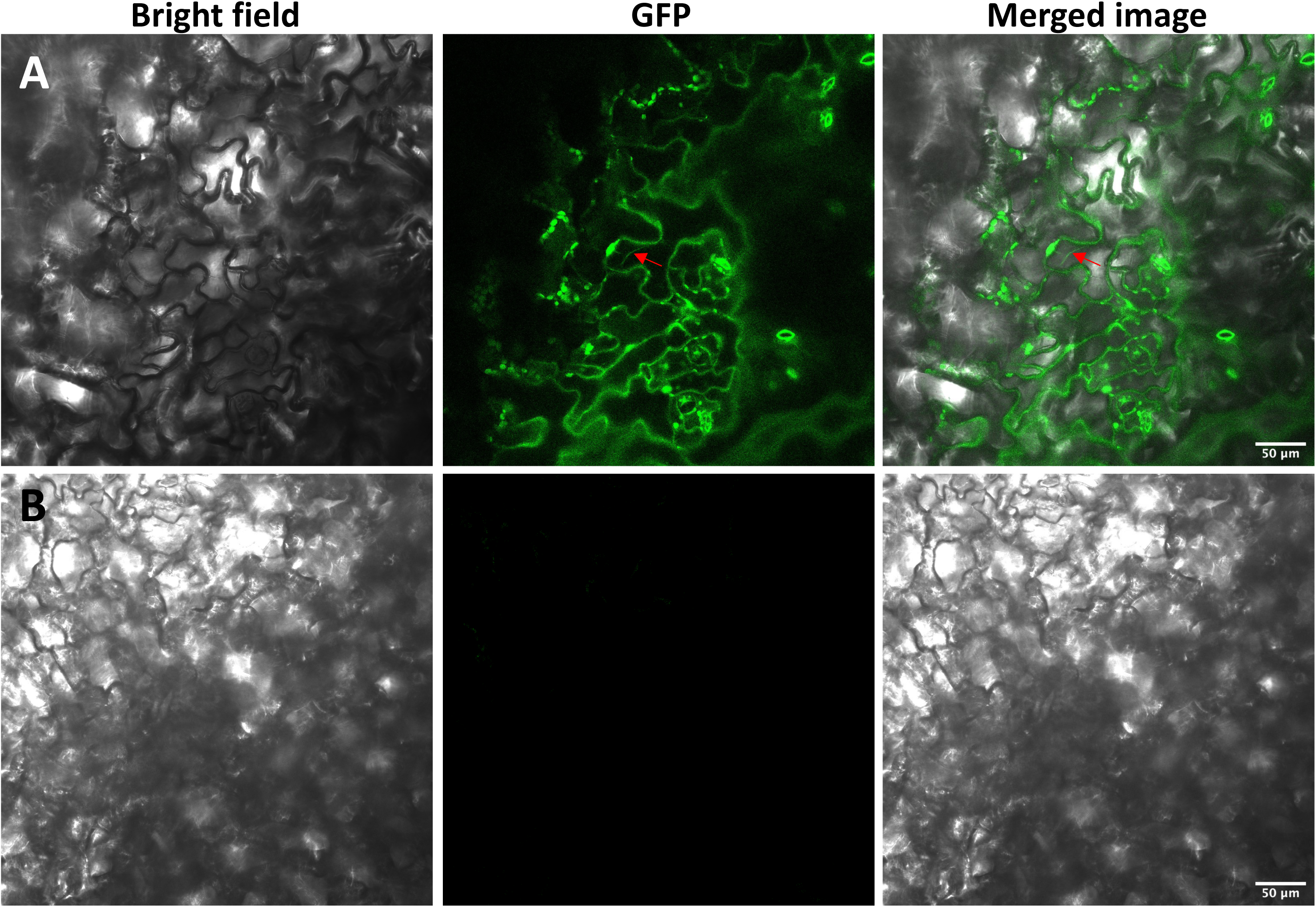
VirE2-iGFP11 delivered from *Agrobacterium* localizes to the periphery of *Arabidopsis* cells and along filaments. **(A)** *Arabidopsis* leaves expressing GFP1-10 were infiltrated with an *Agrobacterium* strain expressing VirE2-iGFP11. The leaves were imaged one day later by confocal microscopy. **(B)** Non-infiltrated leaves of *Arabidopsis* expressing GFP1-10. Red arrows indicate VirE2 along cytoplasmic filaments. Bars indicate 50 μm.

## DISCUSSION

### Importance of VirE2 in *Agrobacterium*-mediated plant transformation

*Agrobacterium tumefaciens* can transfer virulence (Vir) proteins and T-DNA into plant, fungal, and yeast cells (Chilton et al., 1977; Bundock et al., 1995; Piers et al., 1996; de Groot et al., 1998). During the transfer process, VirE2, the most abundant virulence protein (Engstrom et al., 1987), and four other Vir proteins exit the bacterium via a VirB/VirD4 type IV secretion system (T4SS) and facilitate transformation (Schrammeijer et al., 2003; Cascales and Christie, 2004; Vergunst et al., 2005). VirE2 is a single-strand (ss) DNA binding protein that binds ssDNA *in vitro* (Zupan et al., 1996). A popular theory proposes that VirE2 binds T-strands that enter the plant cell following infection by *Agrobacterium*, and that the resulting hypothetical T-complexes, composed of VirD2 covalently linked to T-strands and coated by VirE2 molecules, are the form of T-DNA that traffics within the cell (Li et al., 2020). However, T-complexes have not yet been identified in *Agrobacterium*-infected plants.

*virE2* mutant *Agrobacterium* strains are highly attenuated in virulence (Stachel and Nester, 1986). Fewer T-strands can be isolated from plant cells infected by an *Agrobacterium virE2* mutant than from cells infected by a wild-type *Agrobacterium* strain (Yusibov et al., 1994), and the few transformants produced by a *virE2* mutant *Agrobacterium* strain generally harbor large T-DNA deletions, suggesting a protective function of VirE2 on T-strands (Rossi et al., 1996). However, a *virE2* mutant *Agrobacterium* strain can efficiently transform plants expressing VirE2 (*in planta* complementation), indicating both the importance of VirE2 in transformation and that the function of VirE2 in transformation occurs within the plant (Citovsky et al., 1992; Lapham et al., 2021). The efficient rescue of an *Agrobacterium virE2* mutant by *in planta* complementation suggests that VirE2 trafficking within a plant cell can be studied using plant lines that produce VirE2. This is corroborated by our finding that active VirE2-iGFP11, when transferred from *Agrobacterium*, localizes predominantly to the leaf cell periphery (Figure 13), similar to what we observe with *in planta*-synthesized VirE2. In addition, movement of *Agrobacterium*-delivered VirE2 along cytoplasmic filaments occurs with the same velocity as does *in planta*-made VirE2-Venus (Table 1). Taken together, these findings validate our cytological observations of VirE2 localization and interaction with myosin proteins using *in planta*-made VirE2.

As well as its proposed protective function, VirE2 may facilitate T-DNA entry into the plant by forming single-strand DNA-specific pores in the plant plasma membrane, as it can in artificial membranes (Dumas et al., 2001). Following transfer to the plant, VirE2 initially accumulates on the cytoplasmic side of host plasma membranes at points where bacteria contact the plant cell (Li and Pan, 2017). Accumulation of VirE2 on the membrane is facilitated by its interaction with the *Agrobacterium* effector protein VirE3, which also accumulates in the host plasma membrane at contact sites with the bacteria (Lacroix et al., 2005; Li et al., 2018; Li et al., 2020). Interaction of VirE2 with importin alpha may help guide VirE2 towards the nucleus (Bhattacharjee et al., 2008), although this interaction is weak (Chang et al., 2014). Therefore, VirE2 may be involved in the trafficking of the T-complex inside the plant cell. However, our knowledge of the precise trafficking process of VirE2 inside the host cells, and the proteins that coordinate this trafficking to facilitate transformation, remains sparse.

### Importance of the actin cytoskeleton for transformation and intracellular VirE2 movement

The actin cytoskeleton is essential for numerous biological processes in plants, including cell morphogenesis, tip growth, organelle movement, vesicle trafficking, and maintenance of cell polarity ((Hussey et al., 2006; Staiger et al., 2009; Duan and Tominaga, 2018). Actin genes are categorized according to their expression patterns. In *Arabidopsis*, the vegetative actins (*ACT2*, *ACT7*, and *ACT8*) are mainly expressed in roots (Meagher et al., 1999), whereas *ACT4* and *ACT12* are primarily expressed during pollen development (Huang et al., 1996). We conducted transient and stable root transformation assays of *act2* and *act7* mutants (Figure 1A). These assays indicated that mutation of these genes greatly inhibited root transformation. Restoration of transformation-susceptibility by introducing an *ACT7* cDNA into *act7* mutant plants confirmed that the transformation deficiency of the *act7* mutant was caused by mutation of this gene. Mutation of the pollen-expressed *ACT12* gene had no effect on root transformation, as expected. In addition, we measured the velocity of VirE2-Venus movement along actin filaments, which were marked with mCherry-ABD2, in *Arabidopsis* cells. VirE2 aggregates moved with an average velocity of 0.353 ± 0.06 μm/sec in leaf cells and 0.312 ± 0.07 μm/sec in root cells (Table 1), similar to the velocity (0.434 μm/sec) recorded by Yang et al. (2017) within tobacco leaf cells.

Salman et al. (2005) showed that VirE2 binds to microtubules *in vitro*, and Sakalis et al., (2014) showed colocalization of YFP-VirE2 with fluorescently tagged microtubules in yeast and plant cells. However, they did not assay transformation in the presence of microtubule disrupting agents. We observed no difference in transformation between wild-type and *bot-1* mutant *Arabidopsis* roots, which have a disorganized cortical microtubule network, indicating that correctly organized microtubules are not required for efficient transformation of plants.

### Importance of *Arabidopsis* myosin isoforms for transformation

The movement of molecules, vesicles, and organelles along actin filaments relies on the highly conserved myosin motors (Avisar et al., 2008b; Madison and Nebenfuhr, 2013; Ueda et al., 2015; Citovsky and Liu, 2017). Myosins have actin-binding, ATPase, and cargo binding domains, and these activities contribute to a wide range of physiological processes, including root hair growth, organelle motility, tip growth of pollen tubes, and gamete nuclear migration (Prokhnevsky et al., 2008; Peremyslov et al., 2010; Kawashima and Tachikawa, 2014; Madison et al., 2015). *Arabidopsis* contains two classes of myosins, the plant-specific myosin VIII class comprising four members (myosins VIII-1, 2, A, and B) and the myosin XI class comprising 13 members (myosins XI-A through K, and myosins XI-1 and XI-2). Most of the myosin XI proteins are localized in the cytosol and are responsible for intracellular movement of their cargos, including vesicles and organelles (Peremyslov et al., 2008; Peremyslov et al., 2010). Myosin XI-K in particular is important for vesicular trafficking (Yang et al., 2014). Myosin VIII proteins localize predominantly to the cellular periphery, including the plasma membrane, plasmodesmata, and cortical microtubules (Golomb et al., 2008; Haraguchi et al., 2014; Bar-Sinai et al., 2022) although we have found that myosin VIII-2 also localizes to the nucleus (Figure 9). Myosin VIII proteins may be important for intercellular movement of viral proteins and for helping guide plant cell division (Avisar et al., 2008a; Amari et al., 2014; Wu and Bezanilla, 2014).

Upon entry into a plant cell, VirE2 may interact with VirE3 and associate with the plasma membrane where myosin VIII proteins localize, then move towards the nucleus after associating with T-DNA (Tu et al., 2018). We therefore tested the importance of each myosin XI and myosin VIII protein for transformation by conducting transient and stable root transformation assays on individual myosin gene mutants (Figures 1B and 1C). Interestingly, only disruption of the myosin *XI-h* gene substantially (greater than two-fold) inhibited transformation (Supplemental Figure S1A). Myosin XI-H is moderately strongly expressed in all root tissues and is phylogenetically most closely related to myosins XI-B and XI-G, that are not well-expressed in roots, and to myosin XI-2 which does show root expression (Peremyslov et al., 2011). No specific cellular function had previously been assigned to myosin XI-H (reviewed by Duan and Tominaga, 2018). We were somewhat surprised not to see an effect of a myosin *xi-k* mutation on transformation because Yang et al. (2017) showed that RNAi inhibition of *N. benthamiana* myosin XI-K decreased *Agrobacterium*-mediated transformation by ∼50%. However, *Arabidopsis* may encode more myosin XI isoforms than does *N. benthamiana* (Avisar et al., 2008b), permitting functional redundancy of the *Arabidopsis* myosin XI isoforms to conceal a role for *Arabidopsis* myosin XI-K in transformation.

Because the vast majority of single myosin XI or myosin VIII mutants did not affect transformation, we tested the effect of higher order myosin mutants on this process. Roots from plants mutant in two, three, or four myosin XI genes showed increasing deficiency in transformation susceptibility (Figures 1B and 1C). As previously noted, these higher order myosin XI mutants showed increasing root developmental defects as more myosin XI genes were mutated (Supplemental Figure S1B; Ojangu et al., 2007; Peremyslov et al., 2008). Because our *Arabidopsis* transformation assays were conducted on roots, the natural target for *Agrobacterium*-mediated transformation, we were concerned that these root developmental problems may complicate the interpretation of our transformation data. We therefore turned our attention to higher order *Arabidopsis* myosin VIII mutants whose roots appeared normal (Supplemental Figure S1B). These higher order myosin VIII mutants also showed decreased transformation susceptibility (Figures 1B and 1C).

To test the importance of each myosin *VIII* gene in transformation, we individually expressed cDNAs for each of the myosin VIII proteins in the myosin *VIII-1/2/a/b* quadruple mutant. Three of these genes, encoding myosins VIII-2, A, and B, could partially restore transformation susceptibility to the quadruple mutant (Figure 2A), indicating that they play an important role in transformation. Simultaneous expression of pairs of these cDNAs in the quadruple mutant further increased transformation (Figure 2C), indicating that these myosin VIII proteins are not completely redundant with regard to transformation. However, expression of the cDNA encoding myosin VIII-1 did not affect transformation (Figure 2A), suggesting a fundamental difference among these class VIII myosins for their roles in *Agrobacterium*-mediated transformation. As shown in Figures 6, 7, 8, 10 and discussed below, myosins VIII-2, VIII-A, and VIII-B can directly interact with *Agrobacterium* VirE2, an important effector protein for transformation, whereas myosin VIII-1 does not directly interact with VirE2. However, it may indirectly interact with VirE2 within a protein complex. Ectopically expressed myosin VIII-A, VIII-B, VIII-1, and VIII-2 localize strongly to the peripheral regions of root cells (myosin VIII-2 also localizes in the nucleus). Furthermore, expression of the CBDs of myosins VIII-2 and VIII-A decreased the size of VirE2 aggregates that form in the myosin *VIII-1/2/a/b* quadruple mutant, whereas expression of the myosin VIII-1 CBD did not. These observations indicate a fundamental difference among these myosin VIII isoforms in their interaction with VirE2, their subcellular localization, and their effects on transformation. Interestingly, overexpression of a myosin *VIII-1* cDNA in wild-type *Arabidopsis* increased transformation, indicating both that myosin VIII-1 does play some role in transformation, and that this protein is limiting in wild-type roots (Figure 2B). Overexpression of myosin *VIII-A*, *VIII-B*, or *VIII-2* cDNAs in wild-type roots, either individually or in combination, did not alter transformation susceptibility, indicating that these proteins are naturally found in amounts maximal for transformation.

### Some myosin VIII isoforms may tether VirE2 to the plasma membrane until T-strands enter the plant cell

Several studies have indicated that myosin VIII proteins, or their globular tail domains, localize at the plasma membrane, especially at plasmodesmata (Radford and White, 1998; Reichelt et al., 1999; Avisar et al., 2008a; Golomb et al., 2008; Amari et al., 2014). Our results showed that VirE2-Venus substantially co-localizes with each of the four transgenically expressed myosin VIII proteins and their CBDs at the plasma membrane of root cells, although co-localization with myosin VIII-1 is less extensive (Figure 9 and Supplemental Figures S9 and S10). When roots of these transgenic plants were incubated with *Agrobacterium* lacking T-DNA, the association of VirE2 with the peripheral regions of the cell remained largely unchanged. However, incubation of these roots with *Agrobacterium* containing T-DNA relocalized much of the VirE2-Venus signal to the cytosol. In addition, incubation of roots expressing VirE2-Venus (but not any myosin VIII CBD) with *Agrobacterium* containing T-DNA relocalized a portion of the VirE2-Venus signal to the perinuclear area, but not into the nucleus. These results are consistent with previous observations that VirE2 localizes mainly within the cytosol and, especially, perinuclear regions upon infection of cells with an *Agrobacterium* strain that can transfer VirD2/T-strands to plant cells, and that only a very small quantity may enter the nucleus (Li et al., 2020).

To examine how various myosin isoforms may faciltate VirE2 movement from the plasma membrane region to the cytosol, we measured the velocity of VirE2-Venus movement along mCherry-ABD2-labeled actin filaments with and without expression of various myosin VIII or myosin XI-K CBDs. For these experiments, it was assumed that the CBDs could act as domiant negative molecules to bind VirE2, thereby inhibiting VirE2 movement along these actin filaments. VirE2 introduced from *Agrobacterium* into *Arabidopsis* leaf cells moved along filaments with a velocity of 0.323 μm/sec (Table 1). In both the absence and presence of myosin VIII CBD expression, VirE2 trafficked along *Arabidopsis* actin filaments with a velocity of 0.31-0.40 μm/sec (Table 1), a velocity similar (∼0.502 μm/sec) to that calculated by Yang et al. (2017) in tobacco leaves. These data indicate that myosin VIII proteins are not important for intracellular VirE2 trafficking. However, co-expression of the myosin XI-K CBD with VirE2-Venus completely blocked VirE2 movement along actin filaments, suggesting that this myosin isoform is at least partially responsible for intracellular VirE2 trafficking, as suggested by Yang et al. (2017).

### A model for the role of myosin proteins in VirE2 plasma membrane tethering and movement through plant cells

Research from several laboratories, and this study, has informed our revised model for how *Agrobacterium* virulence effector proteins, such as VirE2, and T-DNA enter a plant cell, form complexes, and traffic T-DNA to the nucleus (Figure 14). VirE2 has the ability to generate pores in artificial lipid membranes through which single-strand DNA molecules, such as T-strands, may penetrate (Dumas et al., 2001; Duckely et al., 2005; Grange et al., 2008). VirE2 delivered from *Agrobacterium* localizes to the cytoplasmic side of the plant plasma membrane, held there by interaction with another virulence effector protein, VirE3 (Li et al., 2018; Li et al., 2020). Our data similarly show that both *Agrobacterium-*expressed VirE2 and plant-expressed VirE2-Venus, which can effect transformation by a *virE2* mutant *Agrobacterium* cell, partially localize to the cytoplasmic side of the plasma membrane. We propose that VirE2 is tethered to the plasma membrane by various myosin VIII isoforms. Upon entry of VirD2/T-strands into the plant, VirE2 is released from the plasma membrane and associates with T-strands. This hypothetical T-complex is trafficked along an actin/endoplasmic reticulum network (Yang et al., 2017) by myosin XI-K, or a functionally similar myosin XI protein, towards the nucleus. Consistent with this mechanism of trafficking, we have previously shown that a Rab GTPase, which modulates vesicle trafficking in biosynthetic and endocytic pathways, is important for transformation (Hwang and Gelvin, 2004), and that a VAP33-like SNARE protein, that localizes to ER-plasma membrane contact sites (Wang et al., 2014; Wang et al., 2015), interacts with VirE2 (Lee et al., 2008).

**Figure 14.**
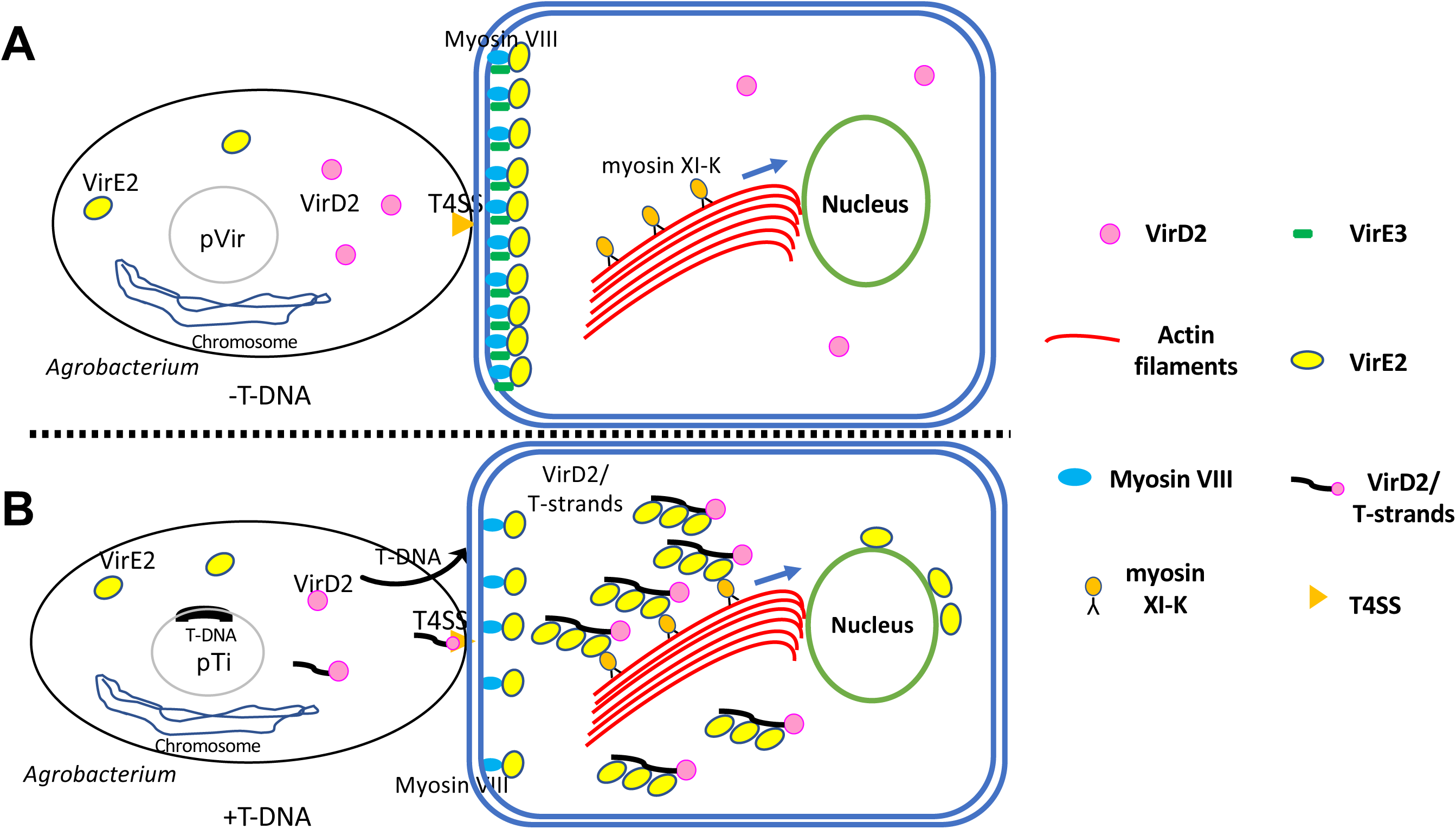
Model of myosin VIII and myosin XI-K involvement in VirE2 trafficking to the perinuclear region. **(A)** *Agrobacterium* virulence effector proteins, including VirE2 and VirE3, are secreted into host cells through a Type IV secretion system. VirE2 and VirE3 interact at the plant plasma membrane. Myosin VIII proteins localize at the plasma membrane and help tether VirE2 to the plasma membrane when no T-DNA is present. **(B)** When T-strands (as a VirD2/T-strand complex) enter the host cell, VirE2 molecules are released and some may bind T-DNA, following which VirE2 interacts with myosin XI-K and moves along actin filaments to the perinuclear region. Not shown are the *Agrobacterium* virulence effector proteins VirD5 and VirF which also are transferred to the plant.

We are aware that one aspect of our analysis does not easily fit this proposed model: Expression of the myosin XI-K CBD completely inhibited detectable VirE2 movement through the cytosol but had no effect on *Agrobacterium*-mediated transformation. Inhibition of VirE2 movement by the myosin XI-K CBD occurred regardless of whether *Agrobacterium* was present to transfer T-DNA into the cells. We note that the requirement for VirE2 in *Agrobacterium*-mediated transformation is not absolute: 1. *Agrobacterium-*mediated yeast transformation does not require VirE2, although mutation of *virE2* results in a ∼10-fold decrease in yeast transformation (Bundock et al., 1995), and 2. Some *A. rhizogenes* strains do not encode VirE2, although they are highly virulent. These strains encode two proteins, GALLS-FL and GALLS-CT, that can functionally complement a *virE2*-mutant *A. tumefaciens* strain, although the amino acid sequence and protein domains of VirE2 and GALLS proteins are highly dissimilar (Hodges et al., 2004, 2006; Ream, 2009). With regard to the results of experiments conducted in this study, we suggest either that a small, undetectable population of VirE2 molecules associates with incoming T-strands and traffics towards the nucleus to effect transformation, or (a somewhat heretical model) that VirE2 does not interact with T-strands in plant cells and that tracking VirE2 movement through the cell does not correlate with intracellular T-DNA movement. Future experiments will attempt to resolve this conundrum.

## MATERIALS AND METHODS

### Plant and bacterial materials and growth conditions

Seeds of wild-type (ecotype Col-0), mutant, and transgenic *Arabidopsis thaliana* were surface-sterilized with 50% bleach and 0.01% SDS, washed five times with sterile water, then placed on Gamborg’s B5 medium (Gibco) with the appropriate antibiotics (when used, hygromycin 10 µg/mL). Seedlings were grown in 16 hr light/8 hr dark conditions at 25°C. *E. coli* strains were grown at 37°C in LB medium (per liter: Tryptone 10 g, NaCl 10 g, Yeast extract 5 g). *A. tumefaciens* strains were grown at 28°C in YEP medium (per liter: Bacto peptone 10 g, Yeast extract 10 g, NaCl 5 g). When used, antibiotic concentrations were: For *E. coli*, ampicillin, 100 μg/ml; kanamycin, 50 μg/ml; spectinomycin, 100 μg/ml; for *A. tumefaciens*, rifampicin, 10 μg/ml; spectinomycin, 100 μg/ml. All bacterial strains and plasmid constructs used in this study are listed in Supplemental Tables S1 and S2.

### Construction of plasmids and binary vectors and generation of transgenic plants

A description of all plasmids used in this study, including details of their construction, is given in Supplemental Table S1. Primers used for these constructions are given in Supplemental Table S3. Final *E. coli* and *A. tumefaciens* strains are given in Supplemental Table S2. Recombinant constructions were transformed into *E. coli* TOP10 or DH10B before characterization and transfer to the appropriate *Agrobacterium* strains by electroporation. For protein expression, plasmids were transferred to *E. coli* BL21DE3. T-DNA binary vectors in *Agrobacterium* were used to transform plants of the Col-0 or the *myosin VIII-1/2/a/b* quadruple mutant background utilizing a floral dip transformation protocol (Clough and Bent, 1998).

### *Agrobacterium*-mediated transient and stable transformation assays

Overnight *Agrobacterium* cultures were diluted 1:10 into 20 ml of YEP medium containing the appropriate antibiotics and grown at 28°C until they reached an A600 = 0.83 (∼10^9^ cells/ml). The cells were collected by centrifugation, washed with sterile 0.9% NaCl, and serially diluted in 0.9% NaCl for transient and stable root transformation assays. For transient transformation assays, infection of *Arabidopsis* root segments was performed with *Agrobacterium* strains containing the T-DNA binary vector pBISN1 (Narasimhulu et al., 1996) as previously described (Tenea et al., 2009). Transformation efficiency was reported as the percentage of root segments stained blue with 5-bromo-4-chloro-3-indolyl-β-D-glucuronic acid (X-gluc) after two days co-cultivation with bacteria on MS medium plus an additional four days on CIM (Callus-Inducing Medium) containing 100 µg/ml timentin. For stable transformation, after two days of cocultivation, root segments were separated onto CIM containing 100 µg/ml timentin and the appropriate selection agent. The plates were incubated at 23°C for four weeks, after which they were scored for callus development on the ends of the roots. Stable transformation efficiency was calculated as the percentage of total segments showing calli. Transient and stable transformation assays were conducted in triplicate, with >100 root segments per replicate.

### Bimolecular fluorescence complementation (BiFC)

A VirE2 coding sequence was fused with cEYFP (C-terminal 64 amino acid residues of an *EYFP* gene) and under the control of a β-estradiol-inducer promoter in the vector pSAT1 to make pSAT1-Pi-VirE2-cEYFP. cDNAs encoding myosin VIII CBDs were amplified using the appropriate primers (Supplemental Table S3), fused with nVenus (N-terminal 174 amino acids of a *Venus* gene) and cloned, under the control of a β-estradiol-inducer promoter in pSAT6, to make pSAT6-Pi-nVenus-myosin VIII CBD. The Pi-VirE2-cEYFP expression cassette and the Pi-nVenus-myosin VIII CBD expression cassettes were digested with *Asc*I and PI-*Psp*I, respectively, and cloned into the same sites in the binary vector pE4437. pE4437 also contains a *hptII* plant selection marker, an expression cassette for a nuclear marker (P35S-Cerulean-NLS), and an actin cytoskeleton marker (P35S-mCherry-actin binding domain2, ABD2). These final constructs were introduced into *A. tumefaciens* GV3101::pMP90 (Koncz and Schell, 1986) by electroporation to make *A. tumefaciens* At2320-At2324, respectively, and used to generate transgenic Col-0 *Arabidopsis* plants for *in vivo* BiFC assays.

### Yeast two hybrid assay

An octopine-type VirE2 gene from pTiA6 was cloned into the bait vector pGBKT7, and the myosin VIII and myosin XI CBD cDNAs were individually cloned into the prey vector pGADT7 (Clontech). As negative controls, empty bait, empty prey, and Lamin C prey plasmids were used (Krendel et al., 2007). A cDNA encoding the VirE2-interacting protein 1 (VIP1) was used as a positive control (Tzfira et al., 2001). Yeast transformations were performed using a lithium acetate method (Sato et al., 1994). Cells were plated onto synthetic dropout (SD)-agar plates lacking tryptophan and leucine. Three days after transformation, yeast cells from the resulting colonies were plated onto SD medium lacking tryptophan, leucine, and histidine (-Trp-Leu-His) supplemented with 3-amino-1,2,4-triazole (3AT) to test for protein interactions.

### *In vitro* protein pull-down assay

The plasmid pET28a (Novagen, San Diego, CA) was used to generate recombinant proteins fused in-frame with a Venus tag or a myc tag. A full-length cDNA encoding VirE2 fused with Venus was cloned into pET28a with *Nco*I+*Not*I. cDNAs encoding the CBDs (amino acids 953-1,166 for myosin VIII-1; amino acids 1,003-1,220 for myosin VIII-2; amino acids 948-1,153 for myosin VIII-A; amino acids 954-1,136 for myosin VIII-B; amino acids 808-1,531 for myosin XI-K) of myosin VIII and XI-K were amplified and cloned into pET28a with *Nco*I+*Bam*HI (myosin VIII CBDs) or *Nco*I+*Not*I (myosin XI-K CBD) to generate myc fusions. The full-length cDNA of VIP1 was cloned into pET28a with *Nco*I+*Bam*HI to use as a myc-tagged positive control, and for negative controls Lamin C fused with a myc tag, or Venus only, were cloned into pET28a with *Sma*I+*Bam*HI. The constructs were transformed into *E. coli* BL21 (DE3). Bacteria were cultured at 16°C for 18 hours in the presence of 0.5 mM isopropylthio-β-galactoside (IPTG). The cells were centrifuged and the pellet was sonicated in cell lysis buffers (20 mM HEPES-KOH pH 7.2, 50 mM potassium acetate, 50 mM KCl, 1 mM EDTA, 1 mM EGTA, 1 mM DTT, 5% glycerol, 0.5% Triton X-100, 1 mM phenylmethylsulfonylfluoride [PMSF]). VirE2-Venus and Venus cell lysates were incubated with GFP-Trap magnetic agarose beads (Chromotek, USA) at 4°C for 2 hr with gentle shaking. The beads were washed three times with cell lysate buffer (0.2% Triton X-100) and then incubated with bacterial lysates from *E. coli* cells expressing myc-tagged VIP1, myc-tagged Lamin C, or myc-tagged myosin VIII/XI-K CBDs proteins that were prepared as described above. The binding reactions were incubated at 4°C for 1 hr with gentle shaking. The beads were washed three times with cell lysate buffer (0.2% Triton X-100) followed by boiling in sodium dodecyl sulfate polyacrylamide gel electrophoresis (SDS-PAGE) buffer for 10 min. Protein samples were loaded onto 10% SDS polyacrylamide gels, subjected to electrophoresis (60 Volts 1.5 hrs for the stacking gel and 100 Volts 4.5 hrs for the separating gel), then either stained with Coomassie blue or subjected to western blot analysis using a 1:1000 dilution of mouse anti-GFP or mouse anti-Myc antibodies (Cell Signaling Technology, USA). After addition of a 1:5000 dilution of horseradish peroxidase-conjugated horse anti-mouse secondary antibodies (Catalog #7076; Cell Signaling Technology, USA), proteins were detected using an ECL kit (LI-COR Biosciences, Lincoln, NE, USA).

### *In vivo* co-immunoprecipitation (co-IP) assay

Transgenic *Arabidopsis* plants (generated using *Agrobacterium* strains At2423-At2427) harboring an inducible expression cassette of either VirE2-Venus, or Venus, and of each myc-tagged myosin VIII or XI-K CBDs were used for *in vivo* protein-protein interaction assays. Transgenic lines expressing, upon induction, myc-tagged CBDs and Venus were used as negative controls (At2428). Four-week old plants were treated with 5 µM β-estradiol for 24 hr, and roots were harvested and ground in liquid nitrogen using a mortar and pestle. A total of 0.25 g of the ground tissue was transferred into a 2 ml tube and 500 µl of CoIP buffer (50 mM Tris-HCl, pH 7.5, 150 mM NaCl, 0.1% Triton X-100, 0.2% Nonidet P-40, 0.6 mM PMSF, and 20 µM MG132 in a Roche protease inhibitor cocktail) was added. The mixture was mixed thoroughly by repeated cycles of 30 sec vortexing followed by chilling on ice. The samples were centrifuged at 4°C, 13000g for 5 min. A total of 20 µl of anti-GFP beads was pre-washed with wash buffer (50 mM Tris-HCl, pH 7.5, 150 mM NaCl, 0.6 mM PMSF, and 20 µM MG132 with Roche protease inhibitor cocktail) three times, the beads were added to the extracted supernatant solution, and the reactions were incubated at 4°C for 1 hr with gentle shaking. The beads were washed five times with wash buffer, proteins were eluted by incubation with 4x LDS sample loading buffer (Invitrogen, CA, USA) at 95℃ for 3 min. The proteins were fractionated by SDS-PAGE and analyzed using western blots, with a 1:1000 dilution of anti-GFP and anti-Myc antibodies, respectively. The proteins were detected using an ECL kit (LI-COR Biosciencces, Lincoln, NE, USA).

### Localization of VirE2-Venus after infection of *Arabidopsis* roots with *Agrobacterium* strains harboring or lacking T-DNA

Transgenic *A. thaliana* lines expressing a *VirE2-Venus* gene under the control of a β-estradiol inducible promoter were treated with 5 µM β-estradiol for 24 hr. Roots were infected with a *virE2* mutant strain *A. tumefaciens* strain (*A. tumefaciens* EHA105 containing a non-polar deletion of *virE2*; At1872) containing the plasmid pBBR1-MCS2::mCherry to track red fluorescent bacteria (At2405). The *Agrobacterium* cells either contained or lacked the T-DNA binary vector pE4672 (*A. tumefaciens* At2404 and At2403, respectively). Overnight bacterial cultures were transferred to fresh AB-glucose medium and grown until the cell density reached an A_600_=0.83. The bacteria were centrifuged and resuspended in induction medium (1x AB salts, 2% glucose, 30 mM MES, 2 mM Na2HPO4, 2 mM NaH2PO4) containing 100 µM acetosyringone and cultured for 16-20 hr. Six hr after cocultivation with bacteria at 10^8^ cfu/ml, root segments were observed by confocal microscopy (Zesis 880 Upright Confocal microscope with a Plan-Apo 20×/0.8 objective). Images from 100 cells of each infection were analyzed using Image J.

### Localization of VirE2-Venus after infiltration of *Arabidopsis* GFP1-10 leaves with an *Agrobacterium* strain expressing VirE2-iGFP11

The delivery of *Agrobacterium*-derived VirE2 to *Arabidopsis* was performed by a split-GFP approach as described previously (Li et al., 2014). Briefly, the large fragment GFP1-10 was expressed in transgenic *Arabidopsis* plants. A plasmid encoding the small GFP11 fragment fused internally with VirE2 was introduced into *A. tumefaciens* At1872 (lacking *virE2*) and used to infiltrate *Arabidopsis* leaves expressing GFP1-10. GFP fluorescence signals were visualized by confocal microscopy. For Agro-infiltration, *Agrobacteria* expressing VirE2-GFP11 was cultured in YEP medium supplied with spectinomycin at 28°C overnight until they reached an A_600_= 0.8. The bacteria were collected by centrifugation and the pellet was washed in infiltration solution (10 mM MES, pH 5.6, 10 mM MgCl2, 200 μM acetosyringone). The bacterial cells were suspended in infiltration solution to an A_600_= 0.8-1. The bacteria were maintained at room temperature for 2-4 hours, then infiltrated into *Arabidopsis* leaves using a 1 mL syringe. Infiltrated plants were maintained in the dark overnight, then transferred to an illuminated growth chamber until imaging.

### Confocal microscopy imaging

For VirE2-Venus trafficking, transgenic plantlets harboring an inducible *Venus-VirE2* transgene (generated using *A. tumefaciens* At2365-At2369) were placed on B5 agar plates containing 100 µg/ml timentin and 20 µg/ml hygromycin and grown vertically for one week. The plants were treated with 10 µM β-estradiol for 9 hr, washed with sterile water, and time lapse images of the roots were taken using a Zeiss 880 confocal microscope equipped with a 20 X water objective. For VirE2 relocalization assays, similarly grown plants were treated with 5 μM β-estradiol for 24 hr, washed with sterile water, and infected with 10^8^ cfu/ml of a *virE2* mutant *Agrobacterium* strain containing or lacking a T-DNA binary vector. Infections were conducted for 8 hr as described above, and images were taken using a Zeiss 880 confocal microscope equipped with a Plan-Apo 20×/0.8 objective.

## SUPPLEMENTAL DATA

**Supplemental Figure S1.** Most *myosin VIII* and *myosin XI* single mutants are not deficient in stable transformation.

**Supplemental Figure S2.** Growth and morphology of wild-type and higher order *myosin VIII* and *XI* mutant plants.

**Supplemental Figure S3.** Effect on transformation of expressing full-length myosin cDNAs in wild-type and *myosin VIII-1/2/a/b* mutant plants.

**Supplemental Figure S4.** Effect on transformation of expressing full-length inducible myosin cDNAs in wild-type and *myosin VIII-1/2/a/b* mutant plants.

**Supplemental Figure S5.** Effect on transformation of expressing pairs of full-length myosin cDNAs in wild-type and *myosin VIII-1/2/a/b* mutant plants.

**Supplemental Figure S6.** Inducible expression of myosin VIII, but not myosin XI-K, CBDs inhibits transformation.

**Supplemental Figure S7.** Co-localization of VirE2-Venus and FM4-64.

**Supplemental Figure S8.** Subcellular localization of VirE2-Venus in transgenic plants. **Supplemental Figure S9.** Myosin VIII CBDs colocalize with the plasma membrane marker PIP2A.

**Supplemental Figure S10.** VirE2 re-localizes to the perinuclear area after infection by an *Agrobacterium* strain that can transfer T-DNA.

**Supplemental Figure S11.** Infection of root cells with an *Agrobacterium* strain capable of transferring T-DNA re-localizes VirE2-Venus from the cellular periphery into the cytoplasm.

**Supplemental Figure S12.** Infection of root cells with an *Agrobacterium* strain capable of transferring T-DNA does not re-localizes VirE2-Venus from the cellular periphery into the cytoplasm when the myosin VIII-1 CBD is expressed.

**Supplemental Figure S13.** Infection of root cells with an *Agrobacterium* strain capable of transferring T-DNA re-localizes VirE2-Venus from the cellular periphery into the cytoplasm.

**Supplemental Figure S14.** Infection of root cells with an *Agrobacterium* strain capable of transferring T-DNA re-localizes VirE2-Venus from the cellular periphery into the cytoplasm.

**Supplemental Figure S15.** Infection of root cells with an *Agrobacterium* strain capable of transferring T-DNA re-localizes VirE2-Venus from the cellular periphery into the cytoplasm.

**Supplemental Figure S16.** Expression of the myosin XI-K CBD does not affect VirE2-Venus localization upon infection by *Agrobacterium*.

**Supplemental Figure S17.** Expression of the myosin XI-K CBD inhibits movement of VirE2-Venus.

## ACKNOWLEDGEMENTS

The authors thank Dr. Valerian V. Dolja, Oregon State University, for providing the *Arabidopsis* mutant lines. We thank the Purdue University Imaging Facility for use of their confocal microscope. We also thank Dr. Kiran Mysore for critical reading of the manuscript. Dr. Praveen Rao contributed data for Figure 1A. Ms. Fang-Yu Hsu contributed data for Figures 4D and 4E.

