## Supplemental Tables 1-3 for "Myosin VIII and XI isoforms interact with *Agrobacterium* VirE2 protein and help direct transport from the plasma membrane to the perinuclear region during plant transformation"

**Supplemental Table 1. Plasmids used in this study.**

| Plasmid | Important features/proteins | Plasmid description | Antibiotic resistance | Reference |
|---|---|---|---|---|
| pE886 | Cloning vector | pBlusescript KS+ | Amp | Kozaki et al., 2004 |
| pE2648 | Cloning vector | pRK310 | Tet | Schmidhauser & Helinski, 1985 |
| pE3086 | EYFP | pSAT6-P35S-cEYFP-N1 | Amp | Citovsky et al., 2006 |
| pE3129 | PIP2A | pSAT6-PIP2A-mRFP | Amp | Gelvin lab strain |
| pE3183 | myc | pSAT6-P35S-myc-MCS | Amp | Gelvin lab strain |
| pE3230 | nVenus | pSAT6-P35S-nVenus-C | Amp | Lee and Gelvin 2014 |
| pE3275 | mCherry | pSAT6-P35S-mCherry-C1-B | Amp | Lee and Gelvin 2014 |
| pE3341 | mCherry | pBBR1-MCS2::mCherry | Kan | From Clay Fuqua |
| pE3447 | Lamin C | pBSKS-Lamin C | Amp | Gelvin lab strain |
| pE3519 | Binary vector | pPZP-*bar*-RCS2 | Spec | Gelvin lab strain |
| pE3522 | Yeast bait vector | pGBKT7 | Kan | Lu et al., 2010 |
| pE3534 | Cerulean | pSAT4A-Cerulean-N | Amp | Gelvin lab strain |
| pE3542 | Venus | pSAT1-Venus-C | Amp | Lee and Gelvin 2014 |
| pE3745 | Yeast prey vector | pGADT7 | Amp | Chien et al., 1991 |
| pE3802 | For N-terminal nVenus tagging of myosin XI-K CBD | pSAT6-P35S-nVenus-myosin XI-K CBD | Amp | Gelvin lab strain |

| | | | | |
|---|---|---|---|---|
| pE3982 | VirE2-cEYFP | pSAT1-$P_{35S}$-VirE2-cEYFP | Amp | Gelvin lab strain |
| pE4014 | VirE2-cCFP | *Nco*I-*Bam*HI fragment from pSAT4-VirE2-cCFP cloned into the same sites in pGBKT7 | Kan | Gelvin lab strain |
| pE4224 | β-estradiol-inducible promoter | pSAT1-β-estradiol-inducible promoter | Amp | Gelvin lab strain |
| pE4282 | VirE2-Venus | pSAT1-Pi-VirE2-Venus | Amp | Gelvin lab strain |
| pE4292 | VirE2-Venus, mCherry-ABD2 | pPZP-Pi-VirE2-Venus-$P_{nos}$-mCherry-ABD2 | Spec | Gelvin lab strain |
| pE4317 | Binary vector; VirE2-Venus, hptII | pPZP-Pi-VirE2-Venus-*hptII* | Spec | Gelvin lab strain |
| pE4381 | mCherry-intron-NLS | pPZP-$P_{35S}$-mCherry-intron-NLS | Spec | Gelvin lab strain |
| pE4437 | Binary vector; Cerulean-NLS, mCherry-ABD2 | pPZP -$P_{nos}$-Cerulean-NLS-$P_{35S}$-mCherry-ABD2 | Spec | Gelvin lab strain |
| pE4438 | VirE2-Venus, Cerulean-NLS | pPZP-Pi-VirE2-Venus-$P_{nos}$-Cerulean-NLS | Spec | Gelvin lab strain |
| pE4515 | pSAT1-$P_{35S}$ | pSAT1-MCS | Kan | Gelvin lab strain |
| pE4668 | Binary vector; *bar* | pPZP-RCS-$P_{nos}$-bar-$T_{nos}$ | Spec | Gelvin lab strain |
| pE4672 | Binary vector; *hptII* | pPZP-$P_{nos}$-*hptII* | Spec | Lee and Gelvin, 2014 |

| | | | | |
|---|---|---|---|---|
| pE4712 | Myosin VIII-1 CBD | PCR fragment of myosin VIII-1 CBD with *KpnI*/*Bam*HI sites, cloned into the *Eco*RV site of pBluescript KS+ to make pBS-myosin VIII-1 CBD | Amp | This study |
| pE4713 | Myosin VIII-2 CBD | PCR fragment of myosin VIII-2 CBD with *Kpn*I/*Bam*HI sites, cloned into the *Eco*RV site of pBluescript KS+ to make pBS-myosin VIII-2 CBD | Amp | This study |
| pE4714 | Myosin VIII-A CBD | PCR fragment of myosin VIII-A CBD with *Kpn*I/*Bam*HI sites, cloned into the *Eco*RV site of pBluescript KS+ to make pBS-myosin VIII-A CBD | Amp | This study |
| pE4715 | Myosin VIII-B CBD | PCR fragment of myosin VIII-B CBD with *Kpn*I/*Bam*HI sites, cloned into the *Eco*RV site of pBluescript KS+ to make pBS-myosin VIII-B CBD | Amp | This study |
| pE4716 | β-estradiol-inducible promoter | PCR fragment of inducible promoter from pE4224 with *Age*I/*Eco*RV sites, cloned into the *Eco*RV site of pBluescript KS+ to make pBS-Pi promoter | Amp | This study |
| pE4720 | pSAT6-Pi-mCherry-C | *Age*I+*Eco*RV digested pE4716, cloned into *Age*I+*Eco*RV digested pE3275, to make pSAT6-Pi-mCherry | Amp | This study |
| pE4721 | mCherry-Myosin VIII-1 CBD | *Kpn*I+*Bam*HI digested pE4712 cloned into *Kpn*I+*Bam*HI digested pE4720 to make pSAT6-Pi-mCherry-myosin VIII-1 CBD | Amp | This study |
| pE4722 | mCherry-Myosin VIII-2 CBD | *Kpn*I+*Bam*HI digested pE4713 cloned into *Kpn*I+*Bam*HI digested pE4720 to make pSAT6-Pi-mCherry-myosin VIII-2 CBD | Amp | This study |
| pE4723 | mCherry-Myosin VIII-A CBD | *Kpn*I+*Bam*HI digested pE4714 cloned into *Kpn*I+*Bam*HI digested pE4720 to make pSAT6-Pi-mCherry-myosin VIII-A CBD | Amp | This study |
| pE4724 | mCherry-Myosin VIII-B CBD | *Kpn*I+*Bam*HI digested pE4715 cloned into *Kpn*I+*Bam*HI digested pE4720 to make pSAT6-Pi-mCherry-myosin VIII-B CBD | Amp | This study |

| | | | | |
|---|---|---|---|---|
| pE4725 | mCherry-Myosin XI-K CBD | *Kpn*I+*Pst*I (Blunt) digested pE3802 cloned into *Kpn*I+*Bam*HI (Blunt) digested pE4720 to make pSAT6-Pi-mCherry-myosin XI-K CBD | Amp | This study |
| pE4726 | mCherry-Myosin VIII-1 CBD, VirE2-Venus, Cerulean-NLS | PI-*Psp*I digested pE4721 cloned into PI-*Psp*I digested pE4438 to make pPZP-Pi-VirE2-Venus-$P_{nos}$-Cerulean-NLS-Pi-mCherry-myosin VIII-1 CBD | Spec | This study |
| pE4727 | mCherry-Myosin VIII-2 CBD, VirE2-Venus, Cerulean-NLS | PI-*Psp*I digested pE4722 cloned into PI-*Psp*I digested pE4438 to make pPZP-Pi-VirE2-Venus-$P_{nos}$-Cerulean-NLS-Pi-mCherry-myosin VIII-2 CBD | Spec | This study |
| pE4728 | mCherry-Myosin VIII-A CBD, VirE2-Venus, Cerulean-NLS | PI-*Psp*I digested pE4723 cloned into PI- *Psp*I digested pE4438 to make pPZP-Pi-VirE2-Venus-$P_{nos}$-Cerulean-NLS-Pi-mCherry-myosin VIII-A CBD | Spec | This study |
| pE4729 | mCherry-Myosin VIII-B CBD, VirE2-Venus, Cerulean-NLS | PI-*Psp*I digested pE4724 cloned into PI-*Psp*I digested pE4438 to make pPZP-Pi-VirE2-Venus-$P_{nos}$-Cerulean-NLS-Pi-mCherry-myosin VIII-B CBD | Spec | This study |
| pE4730 | mCherry-Myosin XI-K CBD, VirE2-Venus, Cerulean-NLS | PI-*Psp*I digested pE4725 cloned into PI-*Psp*I digested pE4438 to make pPZP-Pi-VirE2-Venus-$P_{nos}$-Cerulean-NLS-Pi-mCherry-myosin XI-K CBD | Spec | This study |
| pE4731 | pSAT6-Pi-nVenus-C | *Age*I+*Eco*RV digested pE4716 cloned into *Age*I+*Eco*RV digested pE3230 to make pSAT6-Pi-nVenus | Amp | This study |
| pE4732 | VirE2-cEYFP | *Age*I+*Swa*I digested pE4731 cloned into *Age*I+*Swa*I digested pE3982 to make pSAT1-Pi-VirE2-cEYFP | Amp | This study |
| pE4733 | nVenus-Myosin VIII-1 CBD | *Kpn*I+*Bam*HI digested pE4712 cloned into *Kpn*I+*Bam*HI digested pE4731 to make pSAT6-Pi-nVenus-myosin VIII-1 CBD | Amp | This study |
| pE4734 | nVenus-Myosin VIII-2 CBD | *Kpn*I+*Bam*HI digested pE4713 cloned into *Kpn*I+*Bam*HI digested pE4731 to make pSAT6-Pi-nVenus-myosin VIII-2 CBD | Amp | This study |
| pE4735 | nVenus-Myosin VIII-A CBD | *Kpn*I+*Bam*HI digested pE4714 cloned into *Kpn*I+*Bam*HI digested pE4731 to make pSAT6-Pi-nVenus-myosin VIII-A CBD | Amp | This study |

| | | | | |
|---|---|---|---|---|
| pE4736 | nVenus-Myosin VIII-B CBD | *Kpn*I+*Bam*HI digested pE4715 cloned into *Kpn*I+*Bam*HI digested pE4731 to make pSAT6-Pi-nVenus-myosin VIII-B CBD | Amp | This study |
| pE4737 | nVenus-Myosin XI-K CBD | *Kpn*I+*Not*I digested pE4725 cloned into *Kpn*I+*Not*I digested pE4731 to make pSAT6-Pi-nVenus-myosin XI-K CBD | Amp | This study |
| pE4738 | VirE2-cYFP, Cerulean-NLS, mCherry-ABD2 | *Asc*I digested pE4732 cloned into *Asc*I digested pE4437, to make pPZP-Pi-VirE2-cYFP-$P_{nos}$-Cerulean-NLS-$P_{35S}$-mCherry-ABD2 | Spec | This study |
| pE4739 | pSAT1-Pi-mCherry | *Xho*I (Blunt)/*Not*I digested pE4424 cloned into *Nco*I (Blunt)/*Not*I digested pE3275 to make pSAT1-inducible promoter-mCherry | Amp | This study |
| pE4755 | nVenus-Myosin VIII-A CBD, VirE2-cYFP, Cerulean-NLS, mCherry-ABD2 | PI-*Psp*I digested pE4735 cloned into PI-*Psp*I digested pE4738 to make pPZP-Pi-VirE2-cYFP-$P_{nos}$-Cerulean-NLS-$P_{35S}$-mCherry-ABD2-Pi-nVenus-myosin VIII-A CBD | Spec | This study |
| pE4756 | nVenus-Myosin VIII-B CBD, VirE2-cYFP, Cerulean-NLS, mCherry-ABD2 | PI-*Psp*I digested pE4736 cloned into PI-*Psp*I digested pE4738 to make pPZP-Pi-VirE2-cYFP-$P_{nos}$-Cerulean-NLS-$P_{35S}$-mCherry-ABD2-Pi-nVenus-myosin VIII-B CBD | Spec | This study |
| pE4757 | nVenus-Myosin XI-K CBD, VirE2-cYFP, Cerulean-NLS, mCherry-ABD2 | PI-*Psp*I digested pE4737 cloned into PI-*Psp*I digested pE4738 to make pPZP-Pi-VirE2-cYFP-$P_{nos}$-Cerulean-NLS-$P_{35S}$-mCherry-ABD2-Pi-nVenus-myosin XI-K CBD | Spec | This study |
| pE4758 | nVenus-Myosin VIII-1 CBD, VirE2-cYFP, Cerulean-NLS, mCherry-ABD2 | PI-*Psp*I digested pE4733 cloned into PI-*Psp*I digested pE4738 to make pPZP-Pi-VirE2-cYFP-$P_{nos}$-Cerulean-NLS-$P_{35S}$-mCherry-ABD2-Pi-nVenus-myosin VIII-1 CBD | Spec | This study |
| pE4759 | nVenus-Myosin VIII-2 CBD, VirE2-cYFP, Cerulean-NLS, mCherryABD2 | PI-*Psp*I digested pE4734 cloned into PI-*Psp*I digested pE4738 to make pPZP-Pi-VirE2-cYFP-$P_{nos}$-Cerulean-NLS-$P_{35S}$-mCherry-ABD2-Pi-nVenus-myosin VIII-2 CBD | Spec | This study |

| | | | | |
|---|---|---|---|---|
| pE4774 | myc | PCR fragment of myc tag (pE3183 as template) with *NcoI* and *Not*I (35S terminator), cloned into *Eco*RV site of pBlusescript KS+ (pE886) to make pBS-myc | Amp | This study |
| pE4775 | myc | *Nco*I+*Not*I digested pBS-myc cloned into *Nco*I+*Not*I digested pE4720, to make Pi-pSAT6-myc | Amp | This study |
| pE4776 | myc-Myosin VIII-1 CBD | *Kpn*I+*Bam*HI digested pE4712 cloned into *Kpn*I+*Bam*HI digested pE4775 to make pSAT6-Pi-myc-myosin VIII-1 CBD | Amp | This study |
| pE4777 | myc-Myosin VIII-2 CBD | *Kpn*I+*Bam*HI digested pE4713 cloned into *Kpn*I+*Bam*HI digested pE4775 to make pSAT6-Pi-myc-myosin VIII-2 CBD | Amp | This study |
| pE4778 | myc-Myosin VIII-A CBD | *Kpn*I+*Bam*HI digested pE4714 cloned into *Kpn*I+*Bam*HI digested pE4775 to make pSAT6-Pi-myc-myosin VIII-A CBD | Amp | This study |
| pE4779 | myc-Myosin VIII-B CBD | *Kpn*I+*Bam*HI digested pE4715 cloned into *Kpn*I+*Bam*HI digested pE4775 to make pSAT6-Pi-myc-myosin VIII-B CBD | Amp | This study |
| pE4780 | myc-Myosin XI-K CBD | *Kpn*I+*Not*I digested pE4725 cloned into *Kpn*I+*Not*I digested pE4775 to make pSAT6-Pi-myc-myosin XI-K CBD | Amp | This study |
| pE4781 | mCherry-Myosin VIII-1 CBD, Cerulean-NLS | *Asc*I digested pE4726 self-ligated to make pPZP-$P_{nos}$-Cerulean-NLS-Pi-mCherry-myosin VIII-1 CBD | Spec | This study |
| pE4782 | mCherry-Myosin VIII-2 CBD, Cerulean-NLS | *Asc*I digested pE4727 self-ligated to make pPZP-$P_{nos}$-Cerulean-NLS-Pi-mCherry-myosin VIII-2 CBD | Spec | This study |
| pE4783 | mCherry-Myosin VIII-A CBD, Cerulean-NLS | *Asc*I digested pE4728 self-ligated to make pPZP-$P_{nos}$-Cerulean-NLS-Pi-mCherry-myosin VIII-A CBD | Spec | This study |
| pE4784 | mCherry-Myosin VIII-B CBD, Cerulean-NLS | *Asc*I digested pE472 self-ligated to make pPZP-$P_{nos}$-Cerulean-NLS-Pi-mCherry-myosin VIII-B CBD | Spec | This study |
| pE4785 | mCherry-Myosin XI-K CBD, Cerulean-NLS | *Asc*I digested pE4730 self-ligated to make pPZP-$P_{nos}$-Cerulean-NLS-Pi-mCherry-myosin XI-K CBD | Spec | This study |

| | | | | |
|---|---|---|---|---|
| pE4786 | myc-Myosin VIII-1 CBD, VirE2-Venus | PI-*Psp*I digested pE4776 cloned into PI-*Psp*I digested pE4317 to make pPZP-Pi-VirE2-Venus-Pi-myc-myosin VIII-1 CBD | Spec | This study |
| pE4787 | myc-Myosin VIII-2 CBD, VirE2-Venus | PI-*Psp*I digested pE4777 cloned into PI-*Psp*I digested pE4317 to make pPZP-Pi-VirE2-Venus-Pi-myc-myosin VIII-2 CBD | Spec | This study |
| pE4788 | myc-Myosin VIII-A CBD, VirE2-Venus | PI-*Psp*I digested pE4778 cloned into PI-*Psp*I digested pE4317 to make pPZP-Pi-VirE2-Venus-Pi-myc-myosin VIII-A CBD | Spec | This study |
| pE4789 | myc-Myosin VIII-B CBD, VirE2-Venus | PI-*Psp*I digested pE4779 cloned into PI-*Psp*I digested pE4317 to make pPZP-Pi-VirE2-Venus-Pi-myc-myosin VIII-B CBD | Spec | This study |
| pE4790 | myc-Myosin XI-K CBD, VirE2-Venus | PI-*Psp*I digested pE4780 cloned into PI-*Psp*I digested pE4317 to make pPZP-Pi-VirE2-Venus-Pi-myc-myosin XI-K CBD | Spec | This study |
| pE4791 | New inducible promoter | *Age*I+*Swa*I digested pE4754 cloned into *Age*I+*Swa*I digested pE4731 to make pSAT6-Pi(New)-nVenus | Amp | This study |
| pE4792 | nVenus-Lamin C | *Sma*I+*Bam*HI digested pE3447 cloned into *Sma*I+*Bam*HI digested pE4791 to make pSAT6-Pi(new)-nVenus-Lamin C | Amp | This study |
| pE4814 | Venus | *Nco*I+*Not*I digested pE3542 cloned into *Nco*I+*Not*I digested pET28a, to make pET28a-Venus | Kan | This study |
| pE4815 | VirE2-Venus | *Nco*I+*Not*I digested pE4282 cloned into *Nco*I/+*Not*I digested pET28a, to make pET28a-VirE2-Venus | Kan | This study |
| pE4816 | myc-Myosin VIII-1 CBD | *Nco*I+*Bam*HI digested pE4776 cloned into *Nco*I+*Bam*HI digested pET28a to make pET28a-myc-myosin VIII-1 CBD | Kan | This study |
| pE4817 | myc-Myosin VIII-2 CBD | *Nco*I+*Bam*HI digested pE4777 cloned into *Nco*I+*Bam*HI digested pET28a to make pET28a-myc-myosin VIII-2 CBD | Kan | This study |
| pE4818 | myc-Myosin VIII-A CBD | *Nco*I+*Bam*HI digested pE4778 cloned into *Nco*I+*Bam*HI digested pET28a to make pET28a-myc-myosin VIII-A CBD | Kan | This study |

| | | | | |
|---|---|---|---|---|
| pE4819 | myc-Myosin VIII-B CBD | *Nco*I+*Bam*HI digested pE4779 cloned into *Nco*I+*Bam*HI digested pET28a to make pET28a-myc-myosin VIII-B CBD | Kan | This study |
| pE4820 | myc-Myosin XI-K CBD | *Nco*I+*Not*I digested pE4780 cloned into *Nco*I+*Not*I digested pET28a to make pET28a-myc-myosin XI-K CBD | Kan | This study |
| pE4821 | myc-VIP1 | *Xho*I+*Bam*HI digested pE3395 cloned into *Xho*I+*Bam*HI digested pE4775 to make Pi-pSAT6-myc-VIP1 | Amp | This study |
| pE4822 | myc-VIP1 | *Nco*I+*Bam*HI digested pE4821 cloned into *Nco*I+*Bam*HI digested pET28a to make pET28a-myc-VIP1 | Kan | This study |
| pE4831 | Gal4BD-Myosin VIII-1 CBD | *Nco*I+*Bam*HI digested pE4776 cloned into *Nco*I+*Bam*HI digested pE3745 to make pGADT7-myosin VIII-1 CBD | Amp | This study |
| pE4832 | Gal4BD-Myosin VIII-2 CBD | *Nco*I+*Bam*HI digested pE4777 cloned into *Nco*I+*Bam*HI digested pE3745 to make pGADT7-myosin VIII-2 CBD | Amp | This study |
| pE4833 | Gal4BD-Myosin VIII-A CBD | *Nco*I+*Bam*HI digested pE4778 cloned into *Nco*I+*Bam*HI digested pE3745 to make pGADT7-myosin VIII-A CBD | Amp | This study |
| pE4834 | Gal4BD-Myosin VIII-B CBD | *Nco*I+*Bam*HI digested pE4779 cloned into *Nco*I+*Bam*HI digested pE3745 to make pGADT7-myosin VIII-B CBD | Amp | This study |
| pE4835 | Gal4BD-Myosin XI-K CBD | *Pst*I-Blunt+*Bam*HI digested pE3082 cloned into *Eco*R53KI+*Bam*HI digested pE3745 to make pGADT7-myosin XI-K CBD | Amp | This study |
| pE4836 | Gal4BD-VIP1 | *Nco*I+*Bam*HI digested pE4821 cloned into *Nco*I+*Bam*HI digested pE3745 to make pGADT7-VIP1 | Amp | This study |
| pE4837 | myc-Lamin C | *Sma*I+*Bam*HI digested pE4822 cloned into *Sma*I+*Bam*HI digested pE4792 to make pET28a-myc-Lamin C | Kan | This study |
| pE4843 | Myosin VIII-1 | PCR fragment of myosin VIII-1 full length with *Kpn*I/*Bam*HI sites, cloned into *Eco*RV site of pBlusescript KS+ to make pBS-Myosin VIII-1 full length | Amp | This study |

| | | | | |
|---|---|---|---|---|
| pE4844 | Myosin VIII-2 | PCR fragment of myosin VIII-2 full length with *Kpn*I/*Sma*I sites, cloned into *Eco*RV site of pBlusescript KS+ to make pBS-Myosin VIII-2 full length | Amp | This study |
| pE4845 | Myosin VIII-A | PCR fragment of myosin VIII-A full length with *Kpn*I/*Bam*HI sites, cloned into *Eco*RV site of pBlusescript KS+ to make pBS-Myosin VIII-A full length | Amp | This study |
| pE4846 | Myosin VIII-B | PCR fragment of myosin VIII-B full length with *Kpn*I/*Sma*I sites, cloned into *Eco*RV site of pBlusescript KS+ to make pBS-myosin VIII-B full length | Amp | This study |
| pE4847 | Myosin VIII-1 | *Kpn*I+*Bam*HI digested pE4843 cloned into *Kpn*I+*Bam*HI digested pE3086 to make pSAT6-myosin VIII-1 full length | Amp | This study |
| pE4848 | Myosin VIII-B | *Kpn*I+*Sma*I digested pE4846 cloned into *Kpn*I+*Sma*I digested pE3086 to make pSAT6-myosin VIII-B full length | Amp | This study |
| pE4849 | myc-Myosin VIII-1 CBD, Venus | *Asc*I digested pE3542 cloned into *Asc*I digested pE4786 to make pPZP-35S-Venus-Pi-myc-myosin VIII-1 CBD | Spec | This study |
| pE4850 | myc-Myosin VIII-2 CBD, Venus | *Asc*I digested pE3542 cloned into *Asc*I digested pE4787 to make pPZP-35S-Venus-Pi-myc-myosin VIII-2 CBD | Spec | This study |
| pE4851 | myc-Myosin VIII-A CBD, Venus | *Asc*I digested pE3542 cloned into *Asc*I digested pE4788 to make pPZP-35S-Venus-Pi-myc-myosin VIII-A CBD | Spec | This study |
| pE4852 | myc-Myosin VIII-2 CBD, VirE2-Venus, mCherry-ABD2 | PI-*Psp*I digested pE4777 cloned into PI-*Psp*I digested pE4292 to make pPZP-Pi-VirE2-Venus-$P_{nos}$-mCherry-ABD2-Pi-myc-myosin VIII-2 CBD | Spec | This study |
| pE4853 | myc-Myosin VIII-A CBD, VirE2-Venus, mCherry-ABD2 | PI-*Psp*I digested pE4778 cloned into PI-*Psp*I digested pE4292 to make pPZP-Pi-VirE2-Venus-$P_{nos}$-mCherry-ABD2-Pi-myc-myosin VIII-A CBD | Spec | This study |

| | | | | |
|---|---|---|---|---|
| pE4854 | Myosin VIII-1, hptII | PI-*Psp*I digested pE4847 cloned into PI-*Psp*I digested pE4672 to make pPZP-hpt-XVE-$P_{35S}$-myosin VIII-1 full length | Spec | This study |
| pE4855 | myc-Myosin VIII-1 CBD, VirE2-Venus, mCherry-ABD2 | PI-*Psp*I digested pE4776 cloned into PI-*Psp*I digested pE4292 to make pPZP-Pi-VirE2-Venus-$P_{nos}$-mCherry-ABD2-Pi-myc-myosin VIII-1 CBD | Spec | This study |
| pE4856 | myc-Myosin VIII-B CBD, VirE2-Venus, mCherry-ABD2 | PI-*Psp*I digested pE4779 cloned into PI-*Psp*I digested pE4292 to make pPZP-Pi-VirE2-Venus-$P_{nos}$-mCherry-ABD2-Pi-myc-myosin VIII-B CBD | Spec | This study |
| pE4857 | Myosin VIII-B, hptII | PI-*Psp*I digested pE4848 cloned into PI-*Psp*I digested pE4672 to make pPZP-*hptII*-XVE-$P_{35S}$-myosin VIII-B full length | Spec | This study |
| pE4858 | Myosin VIII-A | *Kpn*I+*Bam*HI digested pE4845 cloned into *Kpn*I+*Bam*HI digested pE4515 to make pSAT1-myosin VIII-A full length | Kan | This study |
| pE4859 | PIP2A-Venus | *Xho*I+*Bam*HI digested pE3129 (PIP2A) cloned into *Xho*I+*Bam*HI digested pE3758 to make pSAT1-PIP2A-Venus | Amp | This study |
| pE4874 | Myosin VIII-A, *hptII* | *Asc*I digested pE4858 cloned into *Asc*I digested pE4672 to make pPZP-XVE-$P_{35S}$-myosin VIII-A full length | Spec | This study |
| pE4875 | Myosin VIII-2 | *Kpn*I+*Sma*I digested pE4844 cloned into *Kpn*I+*Sma*I digested pE4515 to make pSAT1-myosin VIII-2 full length | Kan | This study |
| pE4876 | Myosin VIII-2, *hptII* | *Asc*I digested pE4875 cloned into *Asc*I digested pE4672 to make pPZP-XVE-$P_{35S}$-myosin VIII-2 full length | Spec | This study |
| pE4877 | mCherry-Myosin VIII-1 CBD, Cerulean, PIP2A-Venus | *Asc*I digested pE4859 cloned into *Asc*I digested pE4781 to make pPZP-$P_{nos}$-Cerulean-$P_{35S}$-PIP2A-Venus-Pi-mCherry-myosin VIII-1 CBD | Spec | This study |
| pE4878 | mCherry-Myosin VIII-A CBD, Cerulean, PIP2A-Venus | *Asc*I digested pE4859 cloned into *Asc*I digested pE4783 to make pPZP-$P_{nos}$-Cerulean-$P_{35S}$-PIP2A-Venus-Pi-mCherry-myosin VIII-A CBD | Spec | This study |

| | | | | |
|---|---|---|---|---|
| pE4879 | mCherry-Myosin VIII-B CBD, Cerulean, PIP2A-Venus | *Asc*I digested pE4859 cloned into *Asc*I digested pE4784 to make pPZP-$P_{nos}$-Cerulean-$P_{35S}$-PIP2A-Venus-Pi-mCherry-myosin VIII-B CBD | Spec | This study |
| pE4880 | mCherry-Myosin XI-K CBD, Cerulean, PIP2A-Venus | *Asc*I digested pE4859 cloned into *Asc*I digested pE4785 to make pPZP-$P_{nos}$-Cerulean-$P_{35S}$-PIP2A-Venus-Pi-mCherry-myosin XI-K CBD | Spec | This study |
| pE4889 | myc-Myosin XI-K CBD, VirE2-Venus, mCherry-ABD2 | PI-*Psp*I digested pE4780 cloned into PI-*Psp*I digested pE4292 to make pPZP-Pi-VirE2-Venus-$P_{nos}$-mCherry-ABD2-Pi-myc-myosin XI-K CBD | Spec | This study |
| pE4890 | mCherry-Myosin VIII-2 CBD, Cerulean, PIP2A-Venus | *Asc*I digested pE4859 cloned into *Asc*I digested pE4782 to make pPZP-$P_{nos}$-Cerulean-$P_{35S}$-PIP2A-Venus-Pi-mCherry-myosin VIII-2 CBD | Spec | This study |
| pE4891 | Myosin VIII-A, Myosin VIII-B, hptII | *Asc*I digested pE4858 cloned into *Asc*I digested pE4857 to make pPZP-hpt-XVE- $P_{35S}$-myosin VIII-B full length-myosin VIII-A full length | Spec | This study |
| pE4892 | Myosin VIII-2, Myosin VIII-B, hptII | *Asc*I digested pE4875 cloned into *Asc*I digested pE4857 to make pPZP-hpt-XVE- $P_{35S}$-myosin VIII-B full length-myosin VIII-2 full length | Spec | This study |
| pE4908 | Myosin VIII-B | *Age*I+*Swa*I digested pE4848 cloned into *Age*I+*Swa*I digested pE4754 to make pSAT6-Pi-myosin VIII-B FL | Amp | This study |
| pE4909 | Myosin VIII-1 | *Swa*I+*Bam*HI digested pE4847 cloned into *Swa*I+*Bam*HI digested pE4754 to make pSAT1-Pi-myosin VIII-1 FL | Amp | This study |
| pE4910 | Myosin VIII-2 | *Kpn*I+*Sma*I digested pE4875 cloned into *Kpn*I+*Sma*I digested pE4908 to make pSAT6-Pi-myosin VIII-2 full length | Amp | This study |
| pE4911 | Myosin VIII-A | *Kpn*I+*Bam*HI digested pE4858 cloned into *Kpn*I+*Bam*HI digested pE4909 to make pSAT1-Pi-myosin VIII-A full length | Amp | This study |

| | | | | |
|---|---|---|---|---|
| pE4912 | Myosin VIII-1, *hptII* | *Asc*I digested pE4909 cloned into *Asc*I digested pE4672 to make pPZP-*hptII*-XVE- Pi-myosin VIII-1 full length | Spec | This study |
| pE4913 | Myosin VIII-A, *hptII* | *Asc*I digested pE4911 cloned into *Asc*I digested pE4672 to make pPZP-*hptII*-XVE- Pi-myosin VIII-A full length | Spec | This study |
| pE4914 | Myosin VIII-B, *hptII* | PI-*Psp*I digested pE4908 cloned into PI-*Psp*I digested pE4672 to make pPZP-*hptII*-XVE-Pi-myosin VIII-B full length | Spec | This study |
| pE4915 | Myosin VIII-2, *hptII* | PI-*Psp*I digested pE4910 cloned into PI-*Psp*I digested pE4672 to make pPZP-*hptII*-XVE-Pi-myosin VIII-2 full length | Spec | This study |
| pE4969 | PIP2A | *Xho*I+*Bam*HI digested pE3129 (PIP2A) cloned into *Xho*I+*Bam*HI digested pE3534 to make pSAT4A-$P_{35S}$-PIP2A-Cerulean | Amp | This study |
| pE4970 | Myosin VIII-1 | *Kpn*I+*Bam*HI digested pE4843 cloned into *Kpn*I+*Bam*HI digested pE4791 to make pSAT6-Pi-nVenus-myosin VIII-1 full length | Amp | This study |
| pE4971 | Myosin VIII-2 | *Kpn*I-*Sma*I fragment from pE4844 (myo-VIII-2) cloned into *Kpn*I-*Sma*I sites of pE4791 to make pSAT6-Pi-nVenus-myosin VIII-2 full length | Amp | This study |
| pE4972 | Myosin VIII-A | *Kpn*I-*Bam*HI fragment from pE4845 (myo-VIII-A) cloned into *Kpn*I-*Bam*HI sites of pE4791 to make pSAT6-Pi-nVenus-myosin VIII-A full length | Amp | This study |
| pE4973 | Myosin VIII-B | *Kpn*I-*Sma*I fragment from pE4846 (myo-VIII-B) cloned into *Kpn*I-*Sma*I sites of pE4791 to make pSAT6-Pi-nVenus-myosin VIII-B full length | Amp | This study |
| pE4974 | Myosin VIII-1 | *Kpn*I+*Bam*HI fragment (myo-VIII-I) from pE4843 cloned into *Kpn*I+*Bam*HI sites of pE4720 to make pSAT6-Pi-mCherry-myosin VIII-1 full length | Amp | This study |
| pE4975 | Myosin VIII-2 | *Kpn*I+*Sma*I fragment (myo-VIII-2) from pE4844 cloned into *Kpn*I+*Sma*I sites of pE4720 (pSAT6-Pi-mCherry) to make pSAT6-Pi-mCherry-myosin VIII-2 full length | Amp | This study |

| | | | | |
|---|---|---|---|---|
| pE4976 | Myosin VIII-A | *Kpn*I+*Bam*HI fragment (myo-VIII-A) from pE4845 cloned into the same sites of pE4720 to make pSAT6-Pi-mCherry-myosin VIII-A full length | Amp | This study |
| pE4977 | Myosin VIII-B | *Kpn*I+*Sma*I fragment (myo-VIII-B) from pE4846 cloned into *Kpn*I+*Sma*I sites of pE4720 to make pSAT6-Pi-mCherry-myosin VIII-B full length | Amp | This study |
| pE4978 | GFP1-10 | *Nco*I+*Bst*BI (Blunt) fragment (GFP1-10) from pE4923 cloned into *Nco*I+*Sma*I sites of pE3183 to make pSAT6-$P_{35S}$-GFP1-10 | Amp | This study |
| pE4979 | GFP1-10 | PI-*Psp*I fragment ($P_{35S}$-GFP1-10) from pE4978 cloned into PI-*Psp*I site of pE4668 to make GFP1-10 to make pPZP-$P_{35S}$-GFP1-10-$P_{nos}$-bar | Spec | This study |
| pE4980 | PIP2A-Cerulean | I-*Sce*I fragment ($P_{35S}$-PIP2A-Cerulean) from pE4969 cloned into I-*Sce*I site of pE4738 to make pPZP-Pi-VirE2-cYFP-$P_{35S}$-PIP2A-Cerulean-$P_{35S}$ -ABD2-mCherry | Spec | This study |
| pE4981 | PIP2A-Cerulean | I-*Sce*I fragment ($P_{35S}$-PIP2A-Cerulean) from pE4969 cloned into I-*Sce*I digested pE4438 backbone to make pPZP-Pi-VirE2-Venus-$P_{35S}$-PIP2A-Cerulean | Spec | This study |
| pE4993 | Myosin VIII-1 | PI-*Psp*I fragment (myosin VIII-1 full length) from pE4970 cloned into PI-*Psp*I site of pE4980 to make pPZP-Pi-VirE2-cYFP-$P_{35S}$-PIP2A-Cerulean $P_{35S}$-mCherry-ABD2-Pi-nVenus-myosin VIII-1 full length | Spec | This study |
| pE4994 | Myosin VIII-2 | PI-*Psp*I fragment (myosin VIII-2 full length) from pE4971 cloned into PI-*Psp*I site of pE4980 to make pPZP-Pi-VirE2-cYFP-$P_{35S}$-PIP2A-Cerulean-$P_{35S}$-mCherry-ABD2-Pi-nVenus-myosin VIII-2 full length | Spec | This study |

| | | | | |
|---|---|---|---|---|
| pE4995 | Myosin VIII-A | PI-*Psp*I fragment (myosin VIII-A full length) from pE4972 cloned into PI-*Psp*I site of pE4980 to make pPZP-Pi-VirE2-cYFP-$P_{35S}$-PIP2A-Cerulean-$P_{35S}$-mCherry-ABD2-Pi-nVenus-myosin VIII-A full length | Spec | This study |
| pE4996 | Myosin VIII-B | PI-*Psp*I fragment (myosin VIII-B full length) from pE4973 cloned into PI-*Psp*I site of pE4980 to make pPZP-Pi-VirE2-cYFP-$P_{35S}$-PIP2A-Cerulean-$P_{35S}$-mCherry-ABD2-Pi-nVenus-myosin VIII-B full length | Spec | This study |
| pE4997 | Myosin VIII-1 | PI-*Psp*I fragment (myosin VIII-1 full length) from pE4974 cloned into PI-*Psp*I site of pE4981 to make pPZP-Pi-VirE2-Venus- $P_{35S}$-PIP2A-Cerulean-Pi-mCherry-myosin VIII-1 full length | Spec | This study |
| pE4998 | Myosin VIII-2 | PI-*Psp*I fragment (myosin VIII-2 full length) from pE4975 cloned into PI-*Psp*I site of pE4981 to make pPZP-Pi-VirE2-Venus-$P_{35S}$- $P_{35S}$-PIP2A-Cerulean-Pi-mCherry-myosin VIII-2 full length | Spec | This study |
| pE4999 | Myosin VIII-A | PI-*Psp*I fragment (myosin VIII-A full length) from pE4976 cloned into PI-*Psp*I site of pE4981 to make pPZP-Pi-VirE2-Venus-$P_{35S}$-$P_{35S}$-PIP2A-Cerulean-Pi-mCherry-myosin VIII-A full length | Spec | This study |
| pE5000 | Myosin VIII-B | PI-*Psp*I fragment (myosin VIII-B full length) from pE4977 cloned into PI-*Psp*I site of pE4981 to make pPZP-Pi-VirE2-Venus-$P_{35S}$-PIP2A-Cerulean-Pi-mCherry-myosin VIII-B full length | Spec | This study |
| pE5002 | Myosin XI-K | myosin-XI-K (genomic clone full-length)-mCherry in pMDC32 | Kan | This study |
| pE5003 | Myosin XI-K | *Asc*I fragment (myosinXI-K full length genomic clone) of pE5002 cloned into *Asc*I digested pE4282 to make pMDC-myosin XI-K-mCherry-Pi-VirE2-Venus | Kan | This study |

| | | | | |
|---|---|---|---|---|
| pE5007 | VirE2::iGFP11 | VirE2 with GFP11 fragment cloned into the *Stu*I site of pE2469 | Amp | This study |
| pE5010 | VirE2-iGFP11 | *Bam*HI digested pE5007 ligated with *Bam*HI digested pE2648 to make pRK310-$P_{VirE}$-VirE1-VirE2-iGFP11 | Amp, Tet | This study |
| pE5027 | GFP1-10 | PI-*Psp*I digested pE4978 cloned into PI-*Psp*I digested pE4437 to make pPZP-$P_{nos}$-Cerulean-D2NLS-$P_{35S}$-mCherry-ABD2-$P_{35S}$-GFP1-10 | Spec | This study |

Amp, ampicillin; Gent, gentamicin; Kan, kanamycin; Rif, rifampicin; Spec, spectinomycin; hpt/hptII, hyromycin phosphotransferase II

$P_{35S}$, CaMV 35S promoter; $P_{nos}$, nopaline synthase promoter; Pi, inducible promoter
CBD, cargo binding domain

**Supplemental Table 2. Bacterial strains used in this study.**
***Agrobacterium* strains**

| Stock number | Bacterial host | Plasmid harbored | Important features | Antibiotic resistance | Reference |
|---|---|---|---|---|---|
| At1872 | EHA105ΔVirE2 | ----- | Non-polar *virE2* deletion of pTiEHA105 | Rif | Gelvin lab strain |
| At1879 | At1872 | pBISN2 | pBISN2 *gusA-intron* gene | Kan, Rif, Spec | Gelvin lab strain |
| At2315 | GV3101 | pE4726 | mCherry-myosin VIII-1 CBD<br>VirE2-Venus<br>Cerulean-NLS | Rif, Spec | This study |
| At2316 | GV3101 | pE4727 | mCherry-myosin VIII-2 CBD<br>VirE2-Venus<br>Cerulean-NLS | Rif, Spec | This study |
| At2317 | GV3101 | pE4728 | mCherry-myosin VIII-A CBD<br>VirE2-Venus<br>Cerulean-NLS | Rif, Spec | This study |
| At2318 | GV3101 | pE4729 | mCherry-myosin VIII-B CBD<br>VirE2-Venus<br>Cerulean-NLS | Rif, Spec | This study |
| At2319 | GV3101 | pE4730 | mCherry-myosin XI-K CBD<br>VirE2-Venus<br>Cerulean-NLS | Rif, Spec | This study |
| At2320 | GV3101 | pE4758 | nVenus-myosin VIII-1 CBD<br>VirE2-cYFP<br>Cerulean-NLS<br>mCherry-ABD2 | Rif, Spec | This study |
| At2321 | GV3101 | pE4759 | nVenus-myosin VIII-2 CBD<br>VirE2-cYFP<br>Cerulean-NLS<br>mCherryABD2 | Rif, Spec | This study |

| | | | | | |
|---|---|---|---|---|---|
| At2322 | GV3101 | pE4755 | nVenus-myosin VIII-A CBD<br>VirE2-cYFP<br>Cerulean-NLS<br>mCherry-ABD2 | Rif, Spec | This study |
| At2323 | GV3101 | pE4756 | nVenus-myosin VIII-B CBD<br>VirE2-cYFP<br>Cerulean-NLS<br>mCherry-ABD2 | Rif, Spec | This study |
| At2324 | GV3101 | pE4757 | nVenus-myosin XI-K CBD<br>VirE2-cYFP<br>Cerulean-NLS<br>mCherry-ABD2 | Rif, Spec | This study |
| At2360 | GV3101 | pE4672 | Binary vector; *hptII* | Rif, Spec | This study |
| At2361 | GV3101 | pE4854 | myosin VIII-1<br>*hptII* | Rif, Spec | This study |
| At2362 | GV3101 | pE4857 | myosin VIII-B<br>*hptII* | Rif, Spec | This study |
| At2363 | GV3101 | pE4874 | myosin VIII-A<br>*hptII* | Rif, Spec | This study |
| At2364 | GV3101 | pE4876 | myosin VIII-2<br>*hptII* | Rif, Spec | This study |
| At2365 | GV3101 | pE4852 | myc-myosin VIII-2 CBD<br>VirE2-Venus<br>mCherry-ABD2 | Rif, Spec | This study |
| At2366 | GV3101 | pE4853 | myc-myosin VIII-A CBD<br>VirE2-Venus<br>mCherry-ABD2 | Rif, Spec | This study |
| At2367 | GV3101 | pE4855 | myc-myosin VIII-1 CBD<br>VirE2-Venus<br>mCherry-ABD2 | Rif, Spec | This study |
| At2368 | GV3101 | pE4856 | myc-myosin VIII-B CBD<br>VirE2-Venus<br>mCherry-ABD2 | Rif, Spec | This study |
| At2369 | GV3101 | pE4889 | myc-myosin XI-K CBD<br>VirE2-Venus<br>mCherry-ABD2 | Rif, Spec | This study |
| At2370 | GV3101 | pE4877 | mCherry-myosin VIII-1 CBD<br>Cerulean<br>PIP2A-Venus | Rif, Spec | This study |

| | | | | | |
|---|---|---|---|---|---|
| At2371 | GV3101 | pE4878 | mCherry-myosin VIII-A CBD<br>Cerulean<br>PIP2A-Venus | Rif, Spec | This study |
| At2372 | GV3101 | pE4879 | mCherry-myosin VIII-B CBD<br>Cerulean<br>PIP2A-Venus | Rif, Spec | This study |
| At2373 | GV3101 | pE4880 | mCherry-myosin XI-K CBD<br>Cerulean<br>PIP2A-Venus | Rif, Spec | This study |
| At2374 | GV3101 | pE4890 | mCherry-myosin VIII-2 CBD<br>Cerulean<br>PIP2A-Venus | Rif, Spec | This study |
| At2375 | GV3101 | pE4891 | myosin VIII-A<br>myosin VIII-B<br>*hptII* | Rif, Spec | This study |
| At2376 | GV3101 | pE4892 | myosin VIII-2<br>myosin VIII-B<br>*hptII* | Rif, Spec | This study |
| At2389 | GV3101 | pE4912 | myosin VIII-1<br>*hptII* | Rif, Spec | This study |
| At2390 | GV3101 | pE4915 | myosin VIII-2<br>*hptII* | Rif, Spec | This study |
| At2391 | GV3101 | pE4913 | myosin VIII-A<br>*hptII* | Rif, Spec | This study |
| At2392 | GV3101 | pE4914 | myosin VIII-B<br>*hptII* | Rif, Spec | This study |
| At2403 | At1872 | pE3341 | pBBR1-MCS2::mCherry | Kan | This study |
| At2404 | At2403 | pE4672 | pPZP-RCS-$P_{nos}$-*hptII* | Kan, Spec | This study |
| At2405 | At1872 | pE4381 | pPZP-$P_{35S}$-mCherry-intron-NLS | Spec | This study |

| | | | | | |
|---|---|---|---|---|---|
| At2423 | GV3101 | pE4786 | myc-myosin VIII-1 CBD VirE2-Venus | Rif, Spec | This study |
| At2424 | GV3101 | pE4787 | myc-myosin VIII-2 CBD VirE2-Venus | Rif, Spec | This study |
| At2425 | GV3101 | pE4788 | myc-myosin VIII-A CBD VirE2-Venus | Rif, Spec | This study |
| At2426 | GV3101 | pE4789 | myc-myosin VIII-B CBD VirE2-Venus | Rif, Spec | This study |
| At2427 | GV3101 | pE4790 | myc-myosin XI-K CBD VirE2-Venus | Rif, Spec | This study |
| At2428 | GV3101 | pE4850 | myc-myosin VIII-2 CBD Venus | Rif, Spec | This study |

Kan, kanamycin; Rif, rifampicin; Spec, spectinomycin
CBD, cargo binding domain; FL, full-length

***E. coli* strains**

| **Stock number** | **Host** | **Description of strain** | **Antibiotic resistance** | **Reference** |
|---|---|---|---|---|
| E4823 | BL21(DE3) | pE4814 in BL21DE3 | Kan | This study |
| E4824 | BL21(DE3) | pE4815 in BL21DE3 | Kan | This study |
| E4825 | BL21(DE3) | pE4816 in BL21DE3 | Kan | This study |
| E4826 | BL21(DE3) | pE4817 in BL21DE3 | Kan | This study |
| E4827 | BL21(DE3) | pE4818 in BL21DE3 | Kan | This study |
| E4828 | BL21(DE3) | pE4819 in BL21DE3 | Kan | This study |
| E4829 | BL21(DE3) | pE4820 in BL21DE3 | Kan | This study |
| E4830 | BL21(DE3) | pE4822 in BL21DE3 | Kan | This study |
| Top10 | N/A | $F^-$ *mcr*A Δ(*mrr-hsd*RMS-*mcr*BC) φ80 *lac*ZΔM15Δ *lac*X74 *rec*A1 *ara*Δ139Δ(*ara-leu*)7697 *gal*U *gal*K *rps*L (StrR) *end*A1 *nup*G | Str | Invitrogen |
| BL21(DE3) | N/A | *E. coli* B $F^-$ *dcm ompT hsdS* ($rB^-mB^-$) *gal* λ(DE3) | None | Studier and Moffatt, 1986 |

Kan, kanamycin; Str, streptomycin; N/A, not applicable

**Supplemental Table 3. Primers used in this study.**

| Purpose | Forward primer | Reverse primer |
|---|---|---|
| **myosin cDNA amplification** | | |
| VIII-1 CBD | 5’-CCGGGGTACCAAAGCATCA GTACTTTCCGAGCT-3’ | 5’-CCGCGGATCCTCAATACC TGGTGCTATTTCTCC-3’ |
| VIII-2 CBD | 5’-CCGGGGTACCTCTATGT CTGATCTCCAGAAACG-3’ | 5’-CCGCGGATCCCTAGCC TCTTTTTCCCCACCA-3’ |
| VIII-A CBD | 5’-CCGGGGTACCTATCTTT CCGATCTTCAGCGT-3’ | 5’-CCGCGGATCCTTAATACC TAGTACTCCTCAACCTC-3’ |
| VIII-B CBD | 5’-CCGGGGTACCGTTCTTG CTGATCTCCAGAGCC-3’ | 5’-CCGCGGATCCTCAATAAC TTTTCTTGCACCACCAA-3’ |
| XI-K CBD | 5’-CCGGGGTACCGCAAGG AAAGCTATTGAAGAAGC-3’ | 5’-CCGCGGATCCTTACGAT GTACTGCCTTCTTTAC-3’ |
| VIII-1 FL | 5’-AATTGGTACCATGTCTC AGAAGGTTACTCCATT-3’ | 5’-TTAAGGATCCTCAATAC CTGGTGCTATTTCTCC-3’ |
| VIII-2 FL | 5’-AATTGGTACCATGATGT TATCGGCATCGCCGAAC-3’ | 5’-TTAACCCGGGCTAGCCT CTTTTTCCCCACCA-3’ |
| VIII-A FL | 5’-AATTGGTACCATGGCAC ACAAGGTTAAGGCATC-3’ | 5’-TTAAGGATCCTTAATAC CTAGTACTCCTCAACCTC-3’ |
| VIII-B FL | 5’-AATTGGTACCATGATGA AAAGTTCAGTGAAGG-3’ | 5’-TTAACCCGGGTCAATAA CTTTTCTTGCACCACC-3’ |

| Purpose | Forward primer | Reverse primer |
|---|---|---|
| **Plant genotyping** | | |
| Myosin VIII-1 | 5’-TTCGTGTGAACGTTGATTCTG-3’ | 5’-TCCAGCTTGAATAGATGACGG-3’ |
| Myosin VIII-2 | 5’-CTGAGTTCAGGAGTGTTTCCG-3’ | 5’-TGTACCTCCGAACTGGACAAG-3’ |
| Myosin VIII-A | 5’-AGGTTGTACAACACTGCTGGC-3’ | 5’-ACGAGAAATGGTCTTGTGCTG-3’ |
| Myosin VIII-B | 5’-AGTAAAGCAGGGCCAGTTTTG-3’ | 5’-CTACACTTTGCTTCAGCAGGG-3’ |
| Myosin XI-1 | 5’-TCAAAACGTTGAACTAACCGG-3’ | 5’- TTGTTTGGACGGGTATCTCAG-3’ |
| Myosin XI-2 | 5’-GTGCTCCCGAAGATCCTATT-3’ | 5’-CAGTGCAACCACATGAAGATG-3’ |
| Myosin XI-B | 5’-CAGAAGCAACCCACTTCAGTC-3’ | 5’-GCTACCGAAGGAAGGACTGTC-3’ |
| Myosin XI-C | 5’-TTATGATCACACCCGAGGAAG-3’ | 5’-AAAGACCCACAACAAAGGGAC-3’ |
| Myosin XI-H | 5’-AAATGCTTGAGCTGGAACATG -3’ | 5’-TGCCCTGATTTTTCATTTGTC-3’ |
| Myosin XI-J | 5’-CTCACCTTGCAAAGTGGAGTC-3’ | 5’-CTCAACTTGCAATAAGGCCTG-3’ |
| Myosin XI-E | 5’-TTGGGATGACCCAACTTGTAC-3’ | 5’-GCCTGGAATACACTGAAGCTG-3’ |
| Myosin XI-I | 5’-TTCTGCAATTTCAATTCAGGC-3’ | 5’-CAGCAGACTTCTCCTTCATGG-3’ |
| Myosin XI-K | 5’-GGGTAGCAAGATACTCCTCGG-3’ | 5’-GCAAGAGCAACTCAATTCTGG-3’ |
| Salk T-DNA left border | 5’-TGGTTCACGTAGTGGGCCATCG-3’ | |

CBD, cargo binding domain; FL, full-length
