## Supplemental Figures 1-19 for "Myosin VIII and XI isoforms interact with *Agrobacterium* VirE2 protein and help direct transport from the plasma membrane to the perinuclear region during plant transformation"

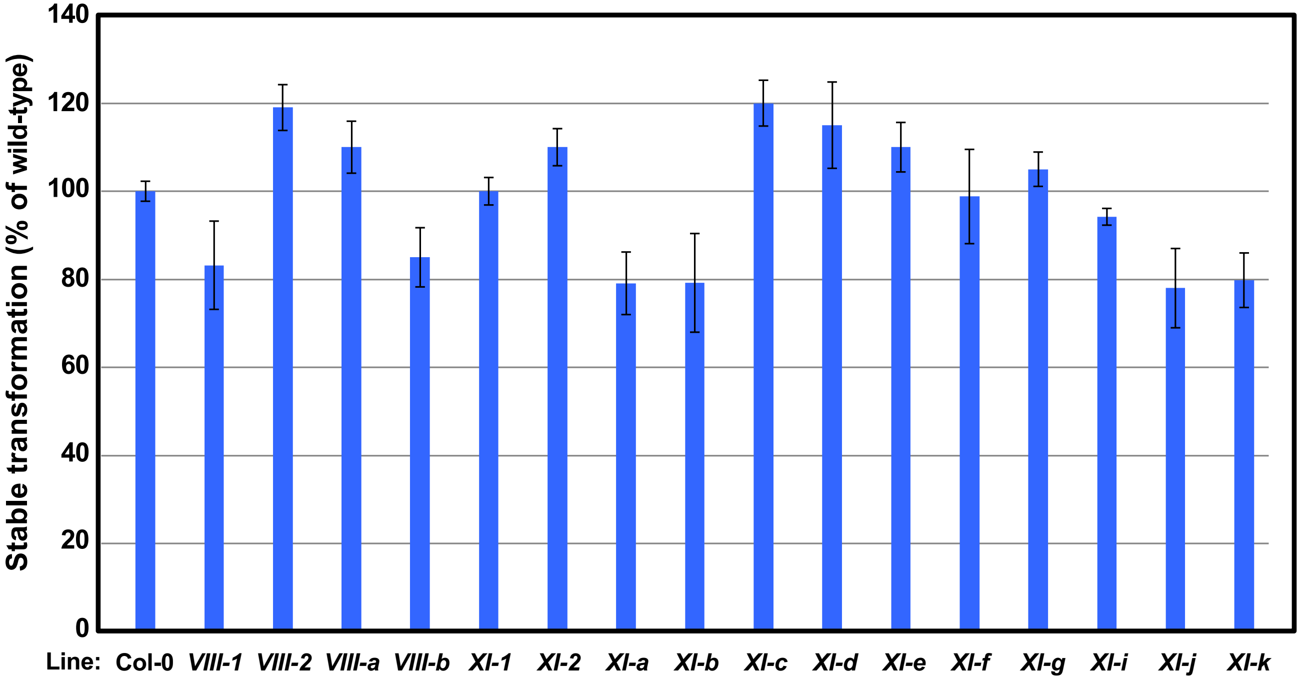


**Supplemental Figure S1. Most *myosin VIII* and *myosin XI* single mutants are not deficient in stable transformation.** Root segments from the indicated *Arabidopsis myosin VIII* or *myosin XI* single mutants, as well as control wild-type plants, were infected with the tumorigenic strain *A. tumefaciens* A208 at the concentration of $10^6$ cfu/ml and the tumors were scored after 30 days of infection. A total of 10-15 plants and >100 segments per plant were tested for stable transformation. Values given are means ± SE. No significant differences were observed compared to wild-type plants.

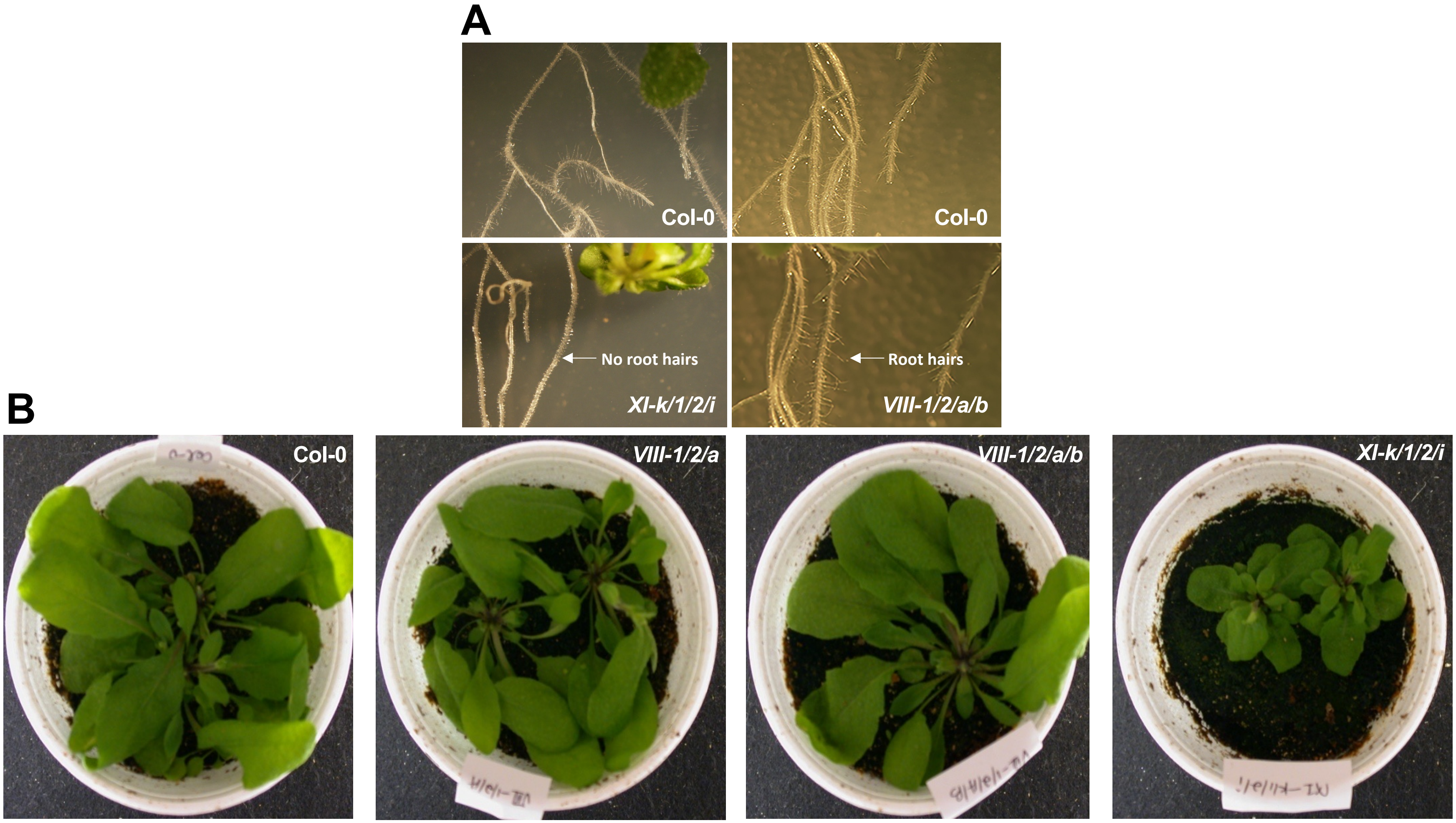


**Supplemental Figure S2. Growth and morphology of wild-type and higher order *myosin VIII* and *XI* mutant plants. (A)** Roots; **(B)** Above-ground tissues.

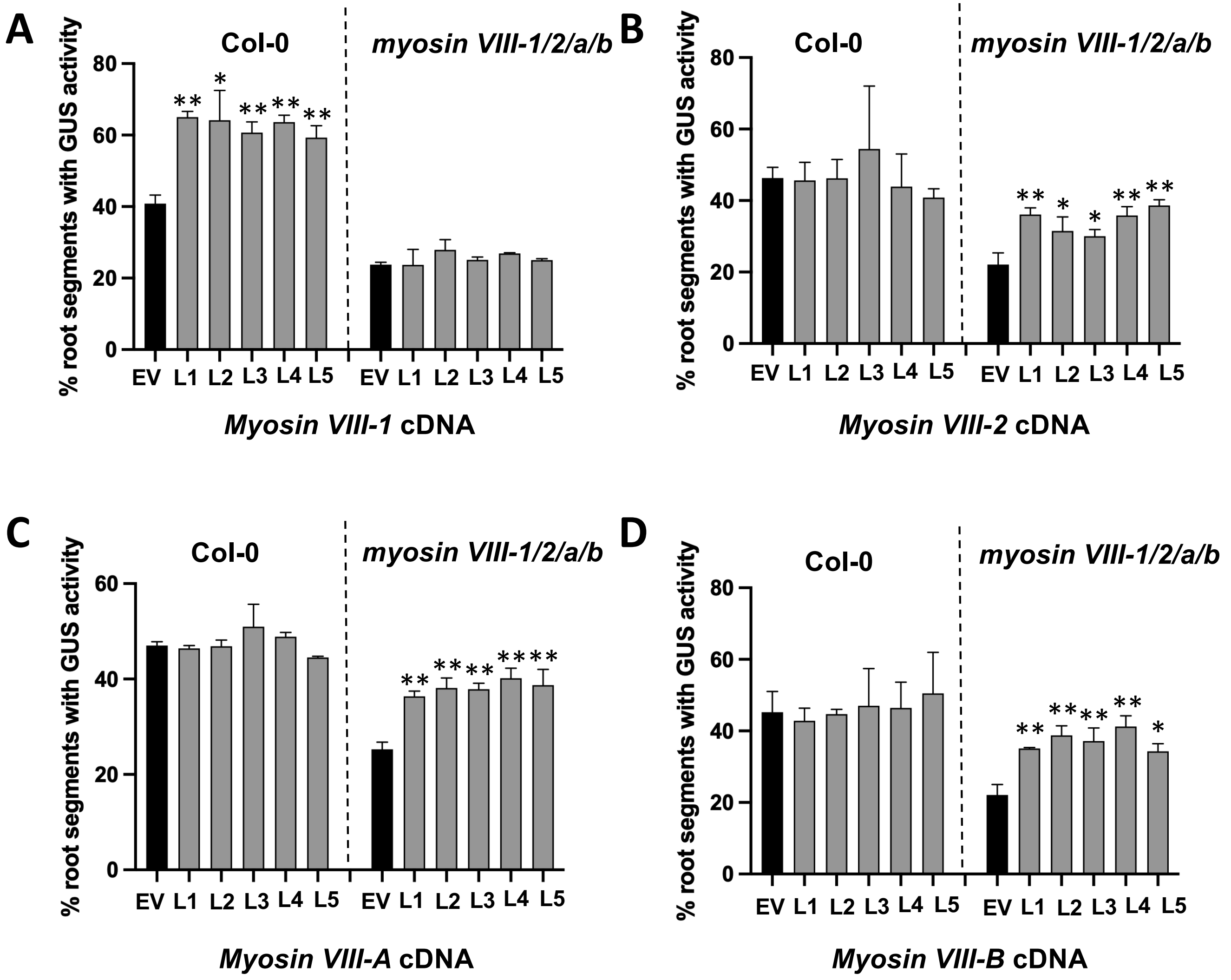


**Supplemental Figure S3. Effect on transformation of expressing full-length myosin cDNAs in wild-type and *myosin VIII-1/2/a/b* mutant plants.** Root segments of five transgenic plants of each independent line, expressing the indicated *myosin VIII* cDNAs in the indicated genetic backgrounds, were infected with *A. tumefaciens* At849 containing the binary vector pBISN1 with a *gusA*-intron gene in the T-DNA region, at $10^7$ cfu/ml. Six days after infection the root segments were stained with X-gluc and the percentage of roots showing GUS activity was calculated. (**A**) *myosin VIII-1* cDNA; (**B**) *myosin VIII-2* cDNA; (**C**) *myosin VIII-A* cDNA; (**D**) *myosin VIII-B* cDNA. Each bar indicates the mean of five plants and >100 root segments/plant ± SE for each transgenic line. Asterisks indicate significant differences of the indicated *myosin VIII* cDNA transgenic plant compared with plants harboring an empty vector (EV). L, Independent transgenic line.
[*t-test*, $*P < 0.05$; $**P < 0.01$].

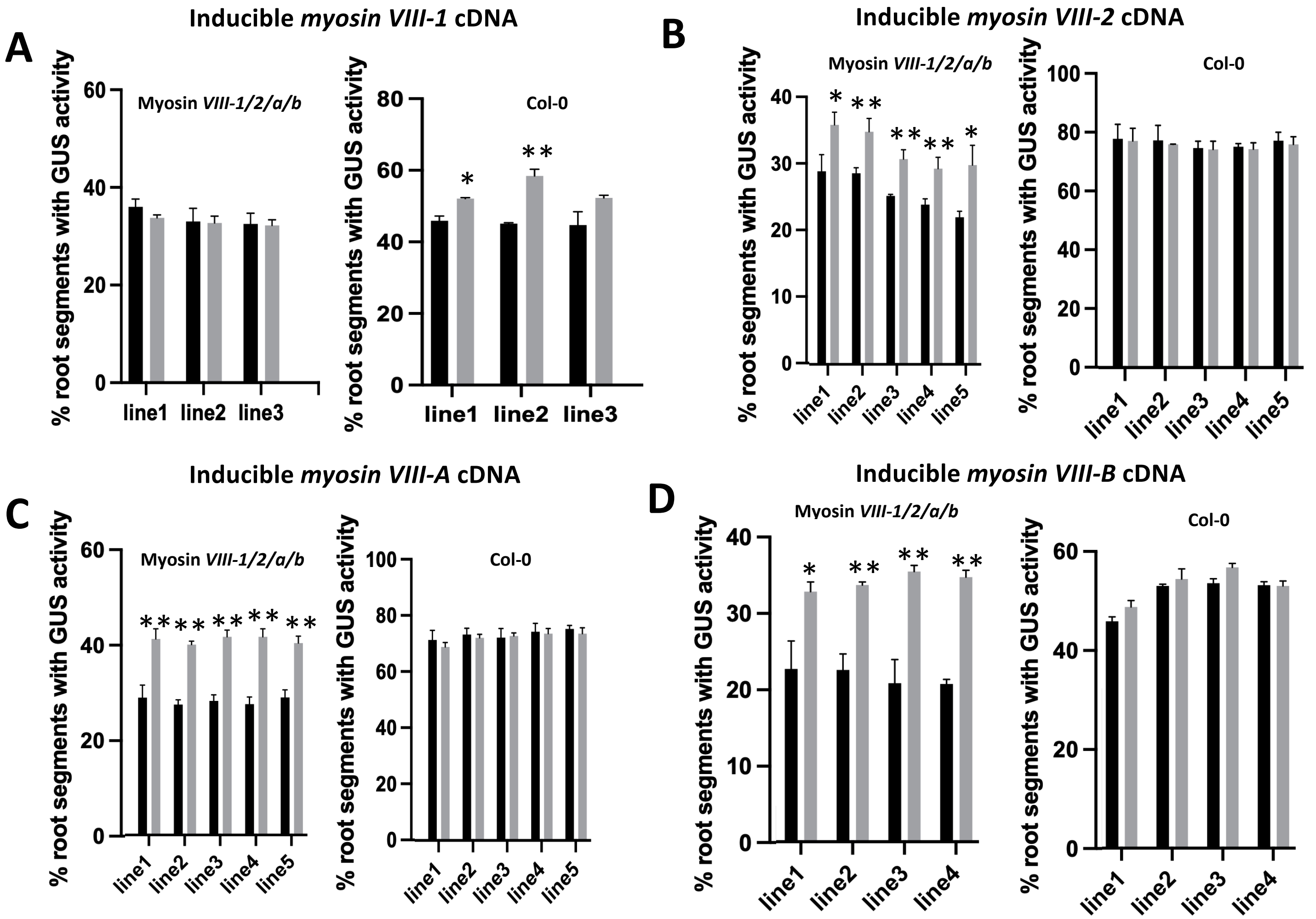


**Supplemental Figure S4. Effect on transformation of expressing full-length inducible myosin cDNAs in wild-type and *myosin VIII-1/2/a/b* mutant plants.** Root segments of five transgenic plants of each independent line, expressing the indicated inducible *myosin VIII* cDNAs in the indicated genetic backgrounds, treated with β-estradiol or control solution for 24 hr, then infected with *A. tumefaciens* At849 containing the binary vector pBISN1 with a *gusA*-intron gene in the T-DNA region, at $10^7$ cfu/ml. Six days after infection the root segments were stained with X-gluc and the percentage of roots showing GUS activity was calculated. (**A**) *myosin VIII-1* cDNA; (**B**) *myosin VIII-2* cDNA; (**C**) *myosin VIII-A* cDNA; (**D**) *myosin VIII-B* cDNA. Each bar indicates the mean of five plants and >100 root segments/plant ± SE for each transgenic line. Asterisks indicate significant differences of the indicated *myosin VIII* cDNA transgenic plant compared with plants treated with control solution. [*t-test*, $*P < 0.05$; $**P < 0.01$].

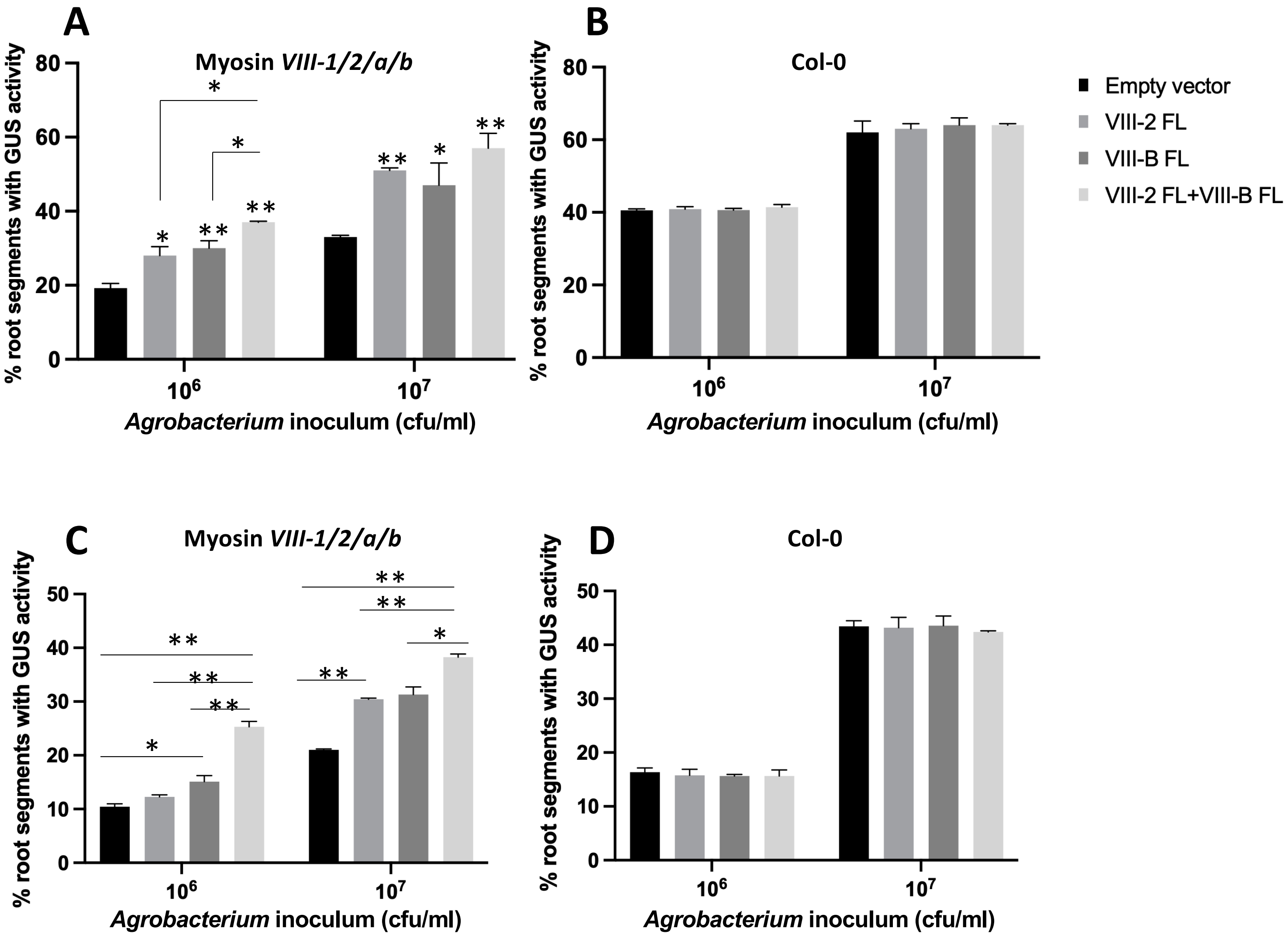


**Supplemental Figure S5. Effect on transformation of expressing pairs of full-length myosin cDNAs in wild-type and *myosin VIII-1/2/a/b* mutant plants.** Root segments of five transgenic plants of each independent line, expressing the indicated single and pairs of *myosin VIII* cDNAs in the indicated genetic backgrounds, were infected with *A. tumefaciens* At849 as mentioned above. The percentage of roots showing GUS activity was calculated. **(A)** *myosin VIII-2*, *myosin VIII-B*, and *myosin VIII-2+VIII-B* cDNAs in the *myosin VIII-1/2/a/b* mutant; **(B)** *myosin VIII-2, myosin VIII-B*, and *myosin VIII-2+VIII-B* cDNAs in Col-0; **(C)** *myosin VIII-A*, *myosin VIII-B*, and *myosin VIII-A+VIII-B* cDNAs in the *myosin VIII-1/2/a/b* mutant; **(D)** *myosin VIII-A*, *myosin VIII-B*, and *myosin VIII-A+VIII-B* cDNAs in Col-0. Each bar indicates the mean of five plants and >100 root segments/plant ± SE for each transgenic line. Except where otherwise indicated, asterisks indicate significant differences of the indicated *myosin VIII* cDNA transgenic plant compared with plants harboring an empty vector.
[*t-test*, **P* < 0.05; ***P* < 0.01].

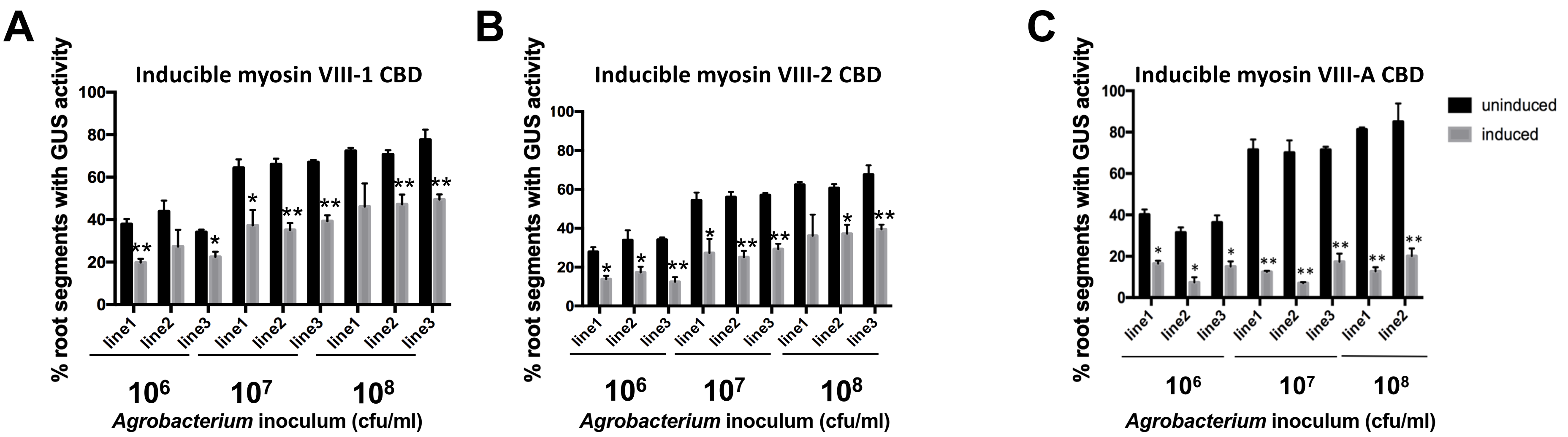


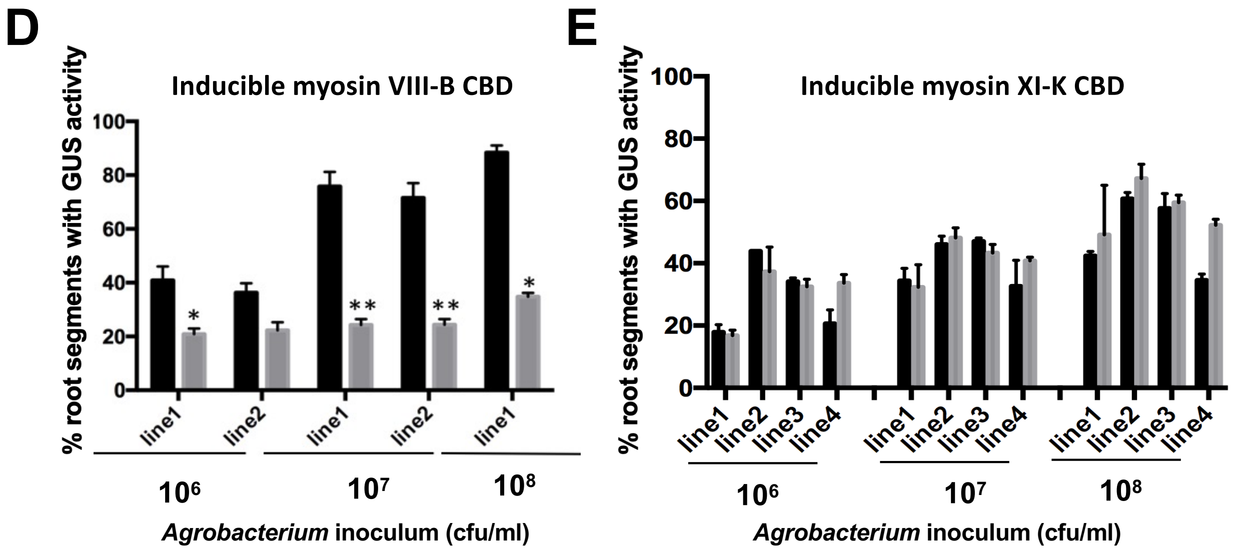


**Supplemental Figure S6. Inducible expression of myosin VIII, but not myosin XI-K, CBDs inhibits transformation.** Root segments of transgenic plants of each independent line, expressing the indicated inducible *myosin VIII* or *myosin XI-K* CBD cDNA, were treated with β-estradiol or control solution for 24 hr, then infected with *A. tumefaciens* At849 containing the binary vector pBISN1 with a *gusA*-intron gene in the T-DNA region, at the indicated $10^6$, $10^7$ and $10^8$ cfu/ml. Six days after infection the root segments were stained with X-gluc and the percentage of roots showing GUS activity was calculated. (**A**) myosin VIII-1 CBD; (**B**) myosin VIII-2 CBD; (**C**) myosin VIII-A CBD; (**D**) myosin VIII-B CBD; (**E**) myosin XI-K CBD. Each bar indicates the mean of five plants and >100 root segments/plant ± SE for each transgenic line. Asterisks indicate significant differences of the indicated *myosin VIII* cDNA transgenic plant compared with plants treated with control solution. [*t-test*, $*P < 0.05$; $**P < 0.01$].

A

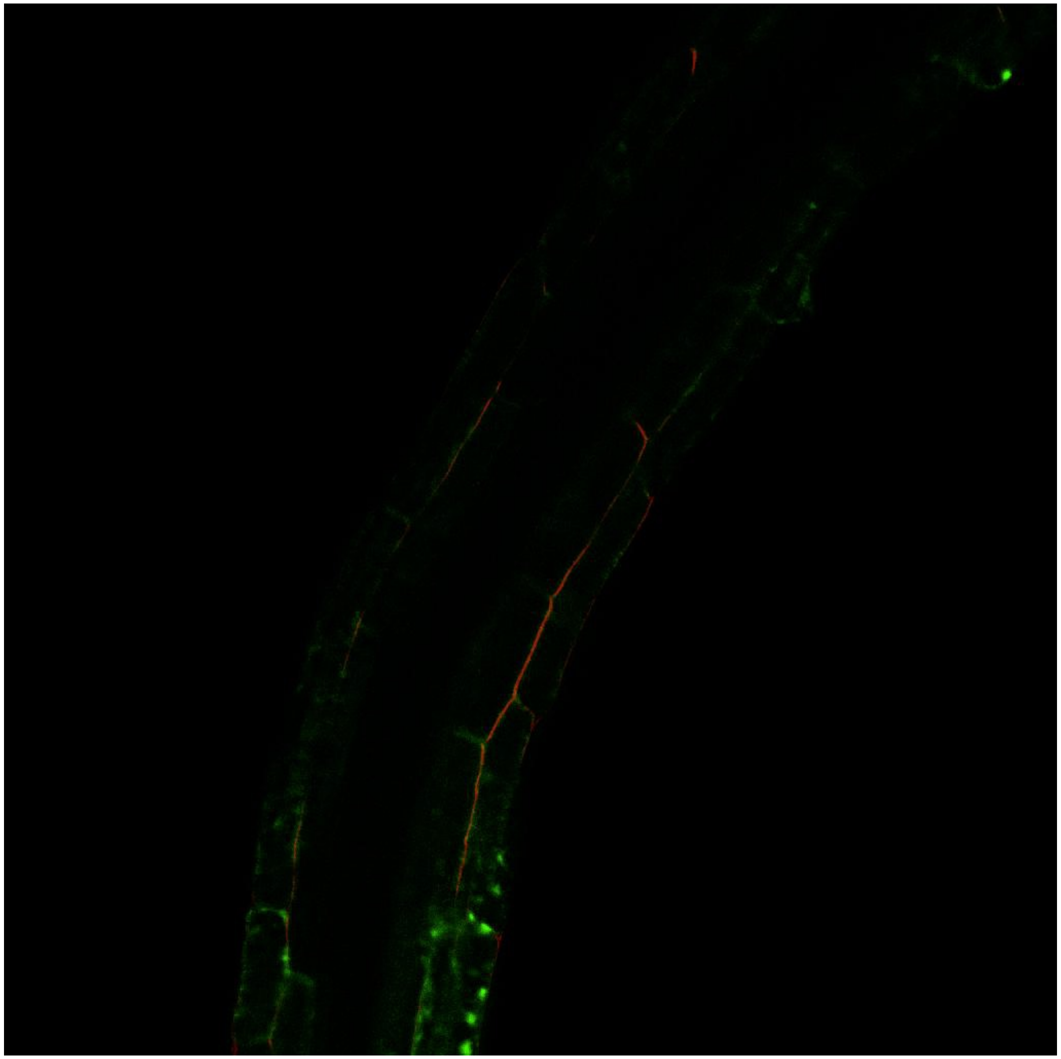

B

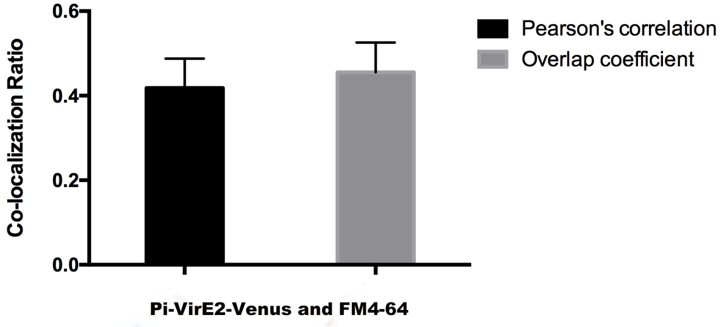


**Supplemental Figure S7. Lack of co-localization of VirE2-Venus and FM4-64. (A)** Inducible VirE2-Venus transgenic plants were treated with β-estradiol for 8 hr, then the root cells were stained with FM4-64 for 5 minutes before imaging. The Z-stack image was taken using a Zesis LSM 880 Upright Confocal Plan-Apo microscope with a 20×/0.8 objective. **(B)** Pearson's and overlap correlations of Venus and red FM4-64 fluorescence. A total of 10-15 transgenic lines and >50 cells were used for quantitative analysis using Image J. Values <0.5 indicate lack of co-localization of the two proteins.

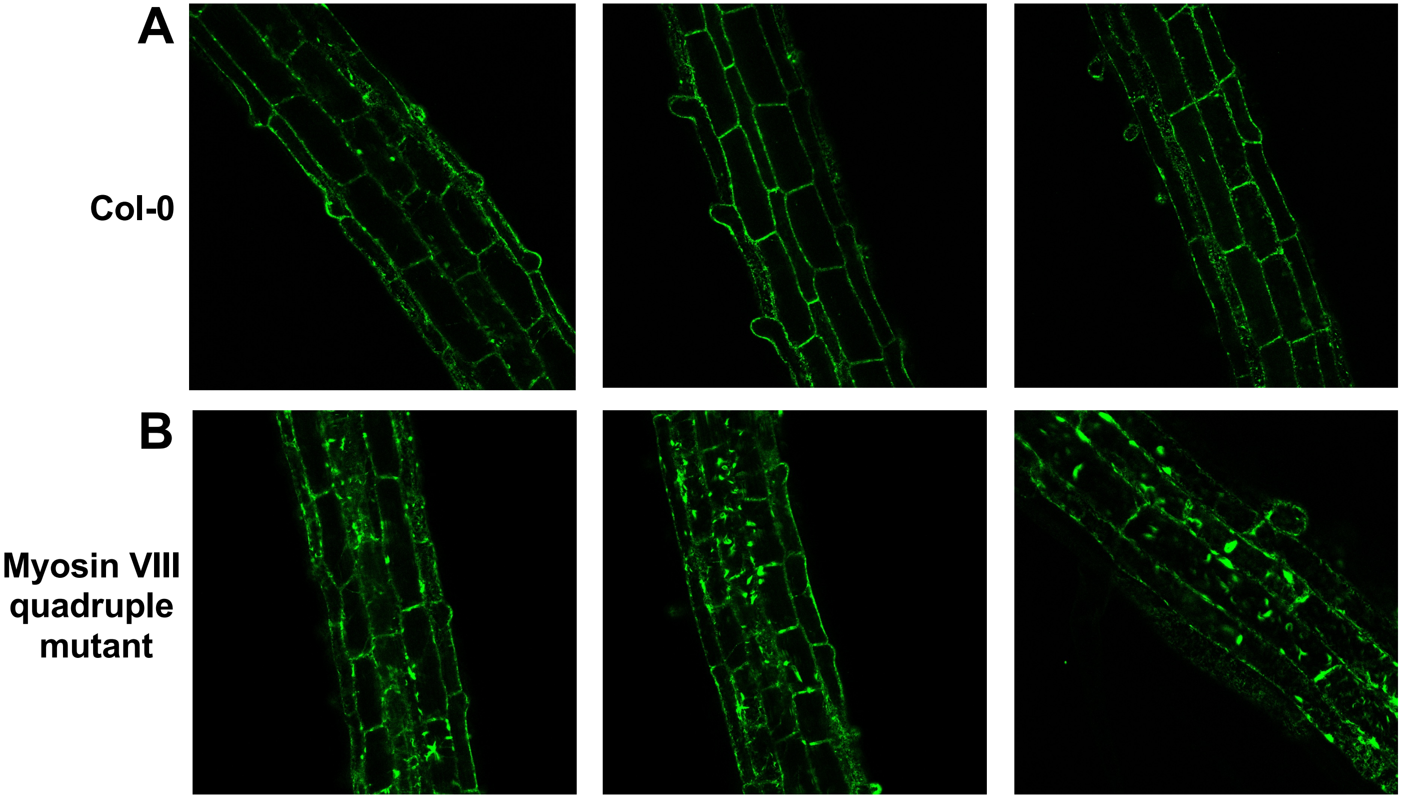


**Supplemental Figure S8. Subcellular localization of VirE2-Venus in transgenic plants. (A)** Inducible VirE2-Venus expressed in wild-type plants. **(B)** Inducible VirE2-Venus expressed in *myosin VIII-1/2/a/b* quadruple mutant plants. Roots from three independent transgenic plants were treated with β-estradiol for 8 hr, and the images were taken using a Zesis LSM 880 Upright Confocal Plan-Apo microscope with a 20×/0.8 objective.

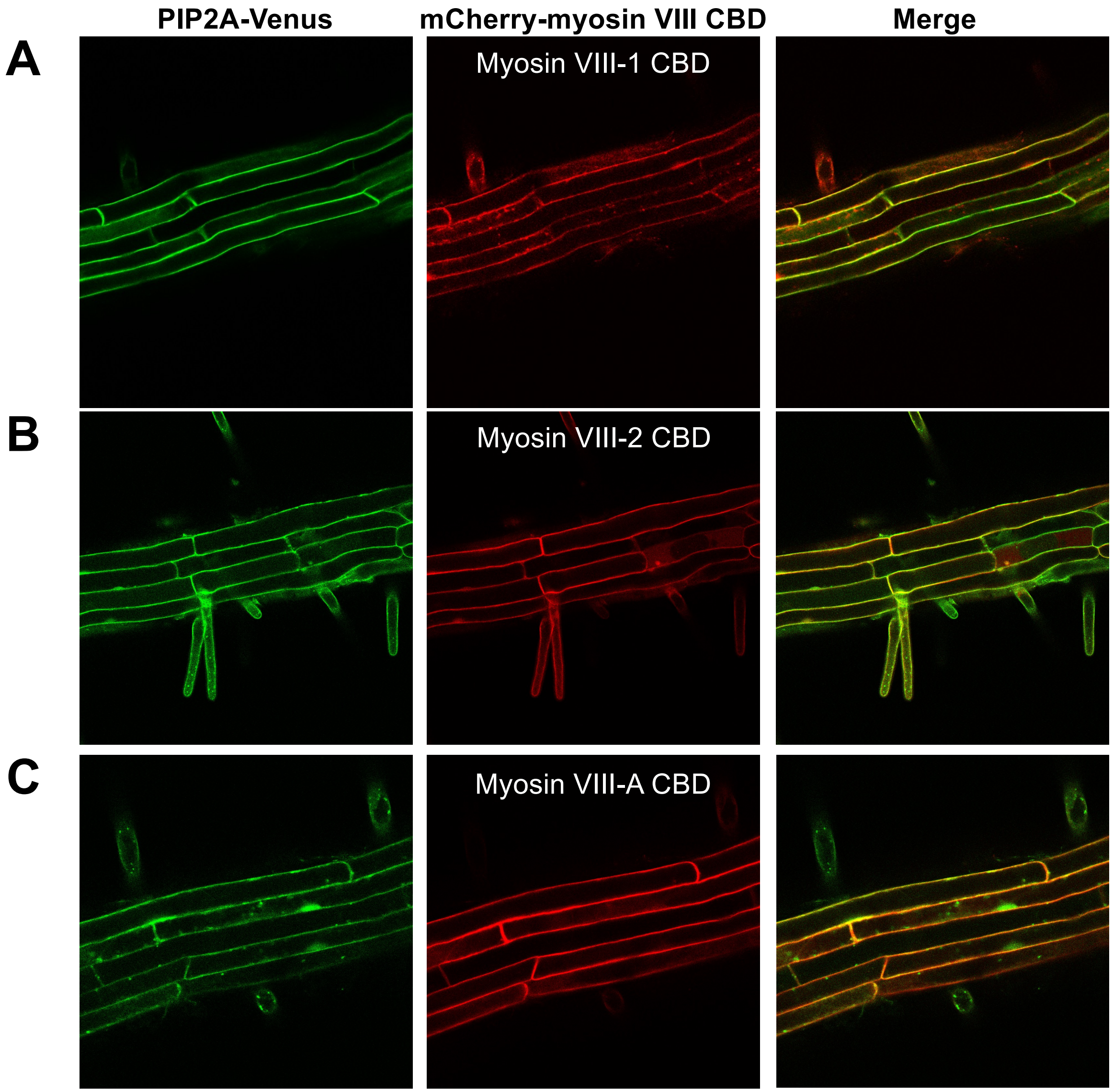


**Supplemental Figure S9. Myosin VIII CBDs colocalize with the plasma membrane marker PIP2A. (A)** Transgenic *Arabidopsis* plant roots that inducibly express an mCherry-myosin VIII-1 CBD and constitutively express PIP2A-Venus. **(B)** Transgenic *Arabidopsis* plants roots that inducibly express an mCherry-myosin VIII-2 CBD and constitutively express PIP2A-Venus. **(C)** Transgenic *Arabidopsis* plants roots that inducibly express an mCherry-myosin VIII-A CBD and constitutively express PIP2A-Venus. A total of 10 independent transgenic lines were treated with β-estradiol for 8 hr, and representative images were taken using a Zesis LSM 880 Upright Confocal Plan-Apo microscope with a 20×/0.8 objective.

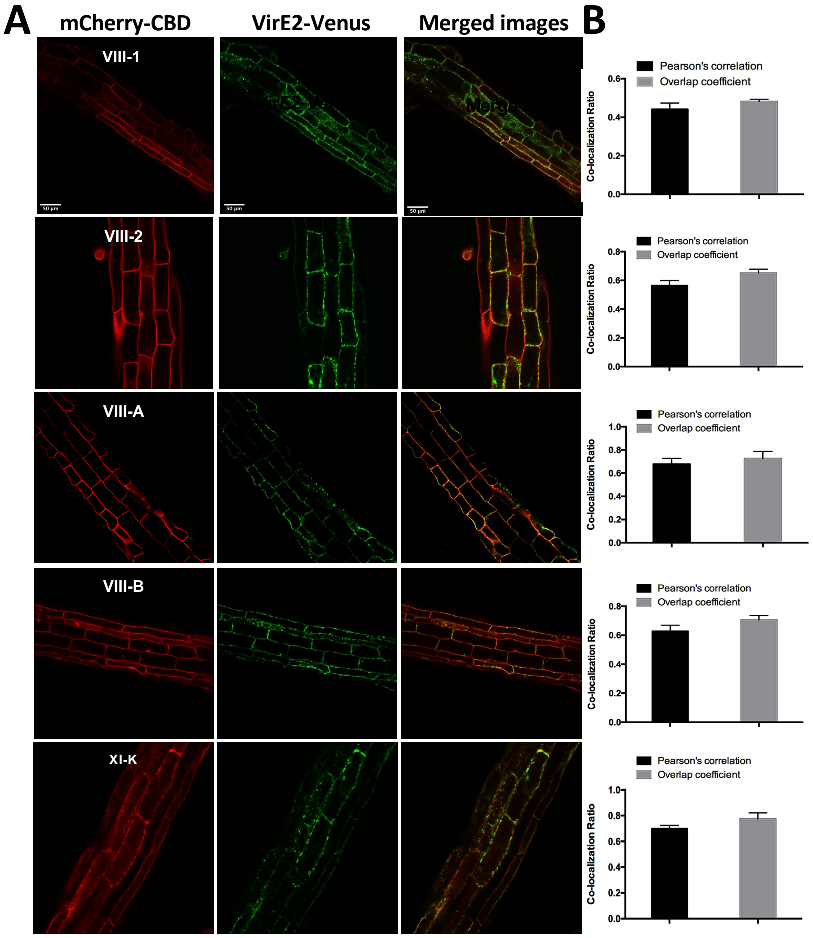


**Figure S10**. **Co-localization of VirE2-Venus and mCherry-myosin CBDs in *Arabidopsis* roots. (A)** Roots of transgenic *Arabidopsis* plants containing VirE2-Venus and mCherry-myosin CBD genes, each under the control of a β-estradiol-inducible promoter, were treated with β-estradiol for 8 hr and examined by confocal microscopy. Bars indicate 50 μm. **(B)** Pearson's correlation and overlap coefficient of Venus and mCherry fluorescence. A total of 10-15 transgenic lines and >50 cells per construct were used for quantitative analysis using Image J. Values >0.5 indicate co-localization of the two proteins for both methods of analysis. The red line indicates a co-localization ratio of 0.5.

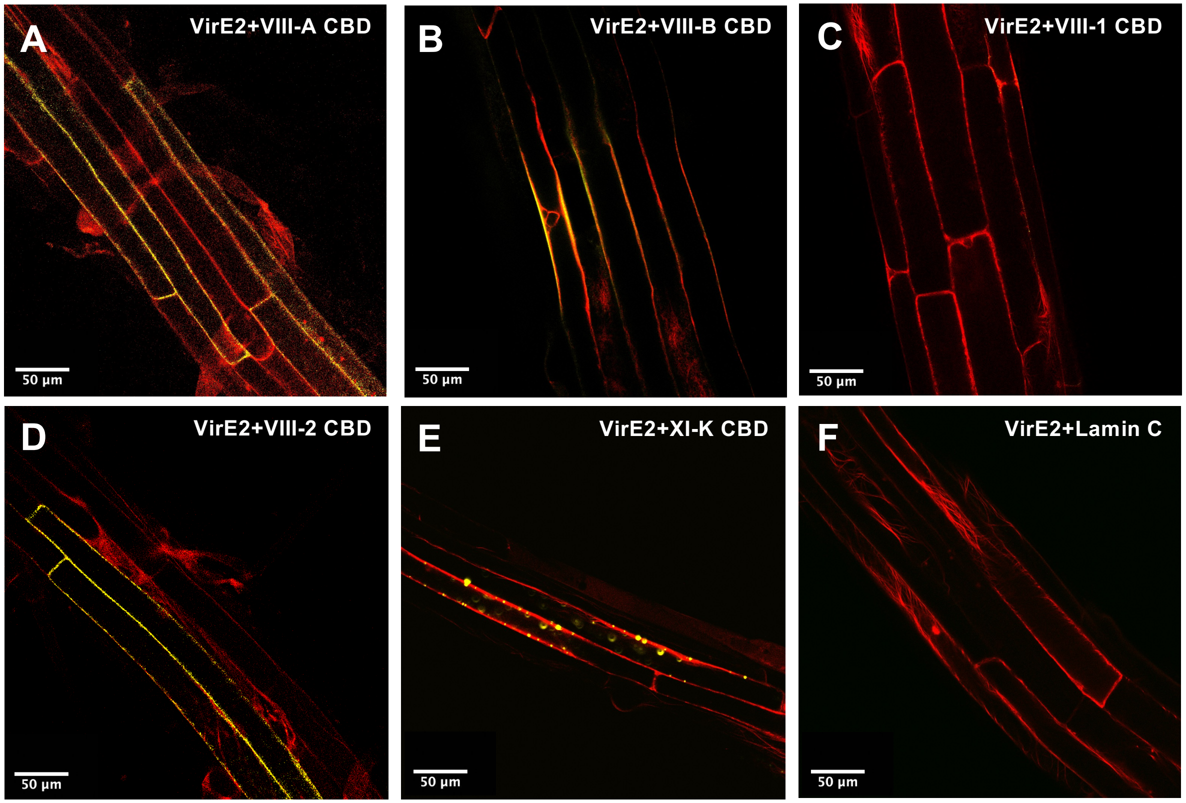

A
VirE2+VIII-A CBD
50 µm
B
VirE2+VIII-B CBD
50 µm
C
VirE2+VIII-1 CBD
50 µm
D
VirE2+VIII-2 CBD
50 µm
E
VirE2+XI-K CBD
50 µm
F
VirE2+Lamin C
50 µm

**Figure S11**. **Bimolecular fluorescence complementation (BiFC) of VirE2 with various myosin CBDs in *Arabidopsis* roots.** Transgenic *Arabidopsis* plants containing VirE2-cEYFP and the indicated myosin CBDs-nVenus, each under the control of a β-estradiol-inducible promoter, were treated with β-estradiol for 48 hr and the fluorescence signal detected by confocal microscopy. VirE2 interacts with the CBDs of **(A),** myosin VIII-2; **(B),** myosin VIII-A; and **(D),** myosin VIII-B at the plasma membrane. VirE2 interacts with the myosin XI-K CBD at the plasma membrane and within the cytoplasm **(E)**. VirE2 does not interact with the myosin VIII-1 CBD **(C)** or Lamin C **(F)**. Yellow fluorescence indicates interaction of VirE2-Venus with the myosin CBD. Actin filaments are labeled by mCherry-ABD2. The figure shows merged images of the Venus and mCherry signals. Bars indicate 50 μm.

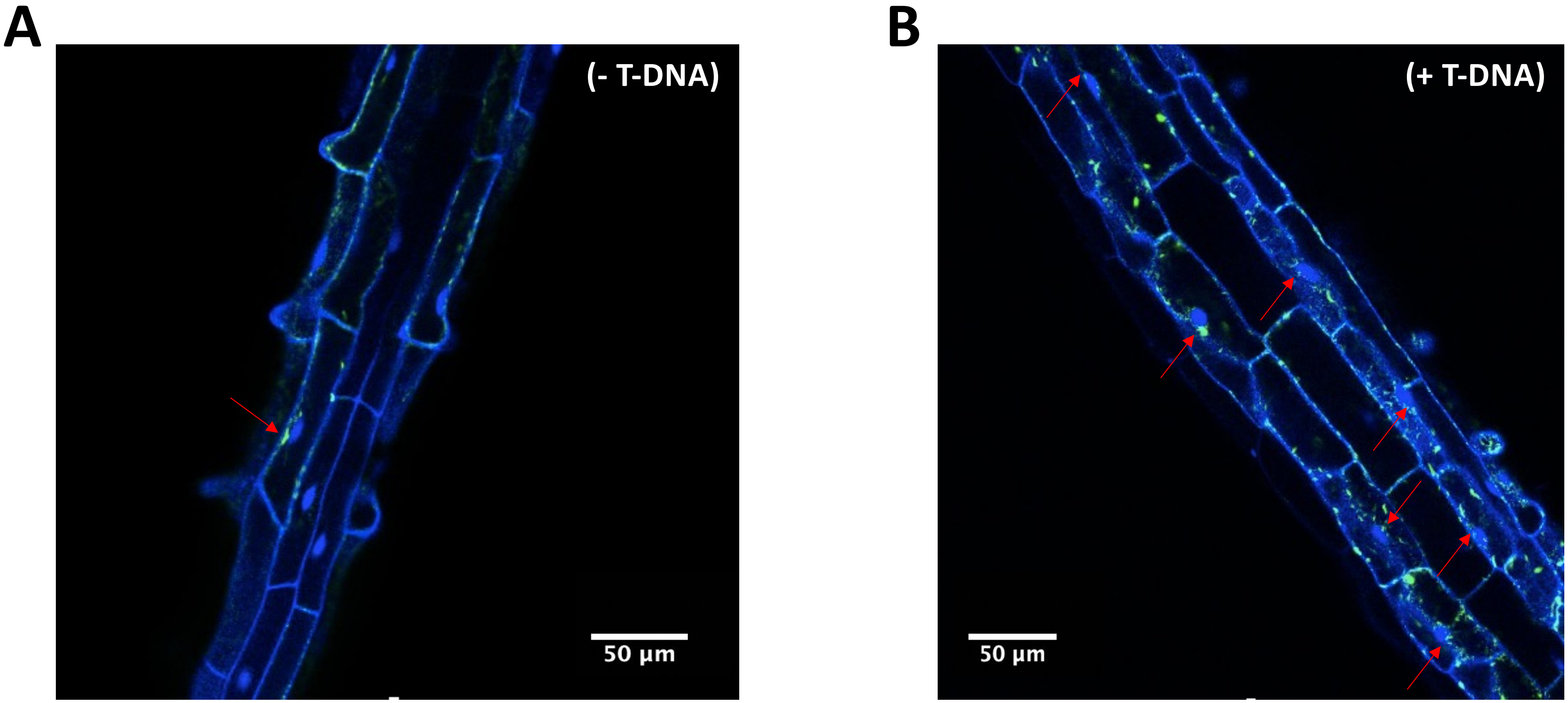


**Supplemental Figure S12. VirE2 re-localizes to the perinuclear area after infection by an *Agrobacterium* strain that can transfer T-DNA.** VirE2-Venus transgenic plants were treated with 5 µM β-estradiol for 24 hr. Root segments were inoculated with $10^8$ cfu/ml of the *virE2* mutant strain *A. tumefaciens* At1872 **(A)** lacking or **(B)** containing a T-DNA region. Images of the elongation zone of the root were taken 8 hr after infection using a Zesis LSM 880 Upright Confocal microscope with a Plan-Apo 20×/0.8 objective. Statistical analysis of VirE2 localization is shown in Figure 11A. Red arrows indicate several examples of perinuclear VirE2-Venus. Bars indicate 50 µm.

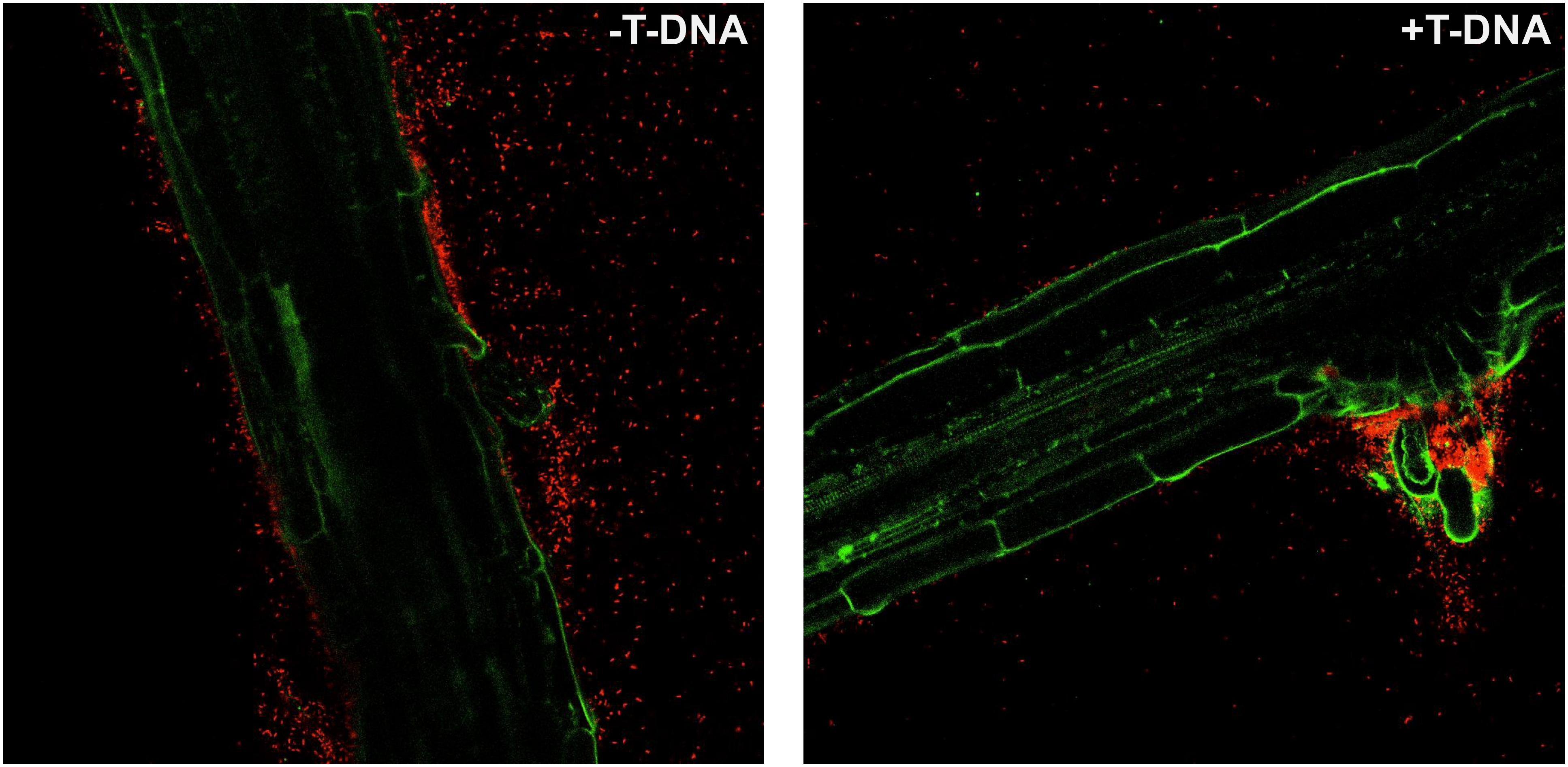


**Supplemental Figure S13. Infection of root cells with an *Agrobacterium* strain capable of transferring T-DNA re-localizes VirE2-Venus from the cellular periphery into the cytoplasm.** Transgenic plants inducibly expressing VirE2-Venus were treated with 5 µM β-estradiol for 24 hr, then inoculated with $10^8$ cfu/ml of the *virE2* mutant strain *A. tumefaciens* At1872 without (left panel) or with (right panel) T-DNA. The *Agrobacterium* cells also expressed an mCherry protein. Z-stack images of cells in the root elongation zone were taken after 8 hr of infection using a Zesis LSM 880 Upright Confocal Plan-Apo microscope using a 20×/0.8 objective.

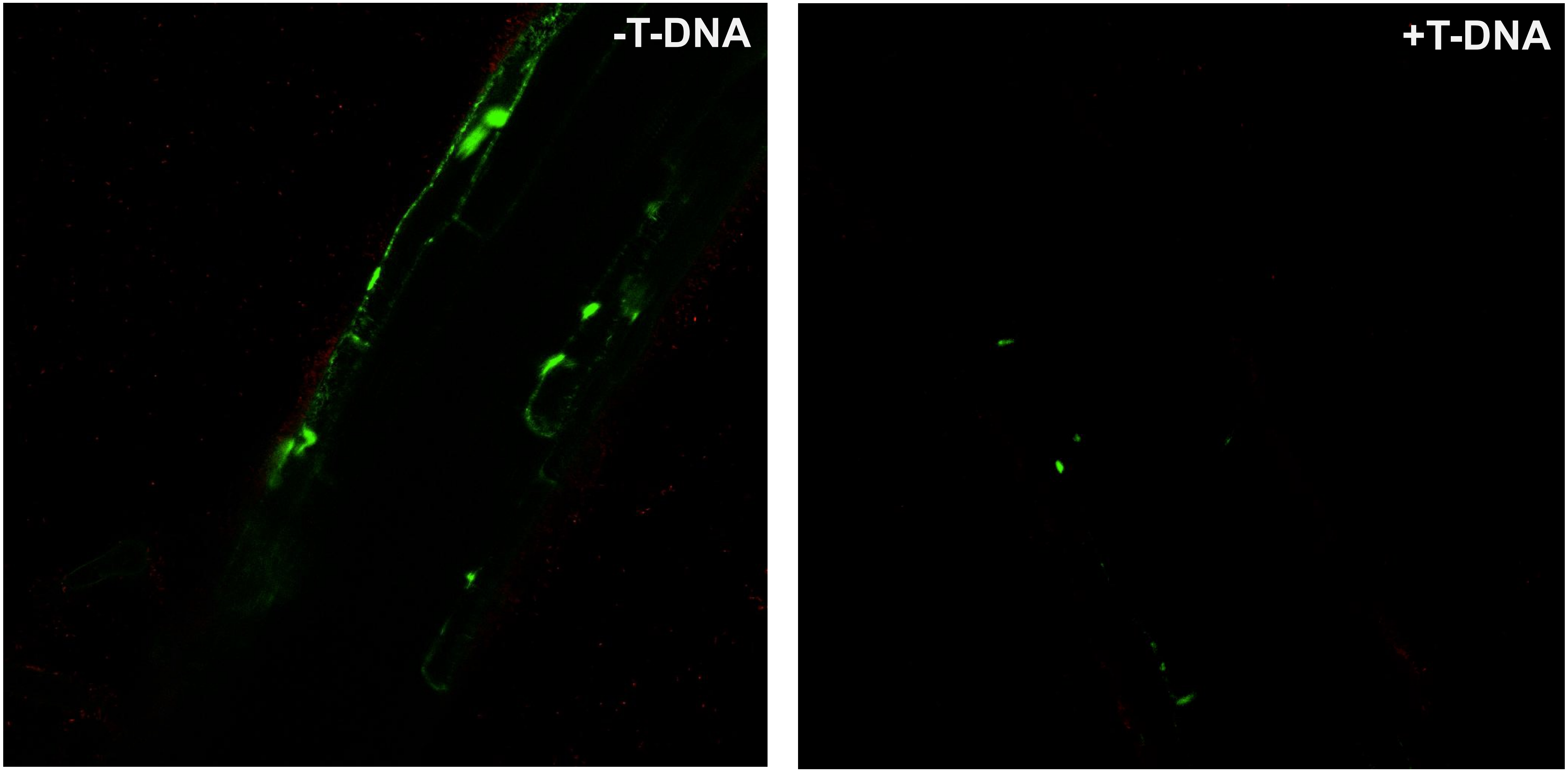


**Supplemental Figure S14. Infection of root cells with an *Agrobacterium* strain capable of transferring T-DNA does not re-localize VirE2-Venus from the cellular periphery into the cytoplasm when the myosin VIII-1 CBD is expressed.** Transgenic plants inducibly expressing the myosin VIII-1 CBD and VirE2-Venus were treated with 5 µM β-estradiol for 24 hr, then inoculated with $10^8$ cfu/ml of the *virE2* mutant strain *A. tumefaciens* At1872 without (left panel) or with (right panel) T-DNA. Z-stack images of cells in the root elongation zone were taken after 8 hr of infection using a Zesis LSM 880 Upright Confocal Plan-Apo microscope using a 20×/0.8 objective.

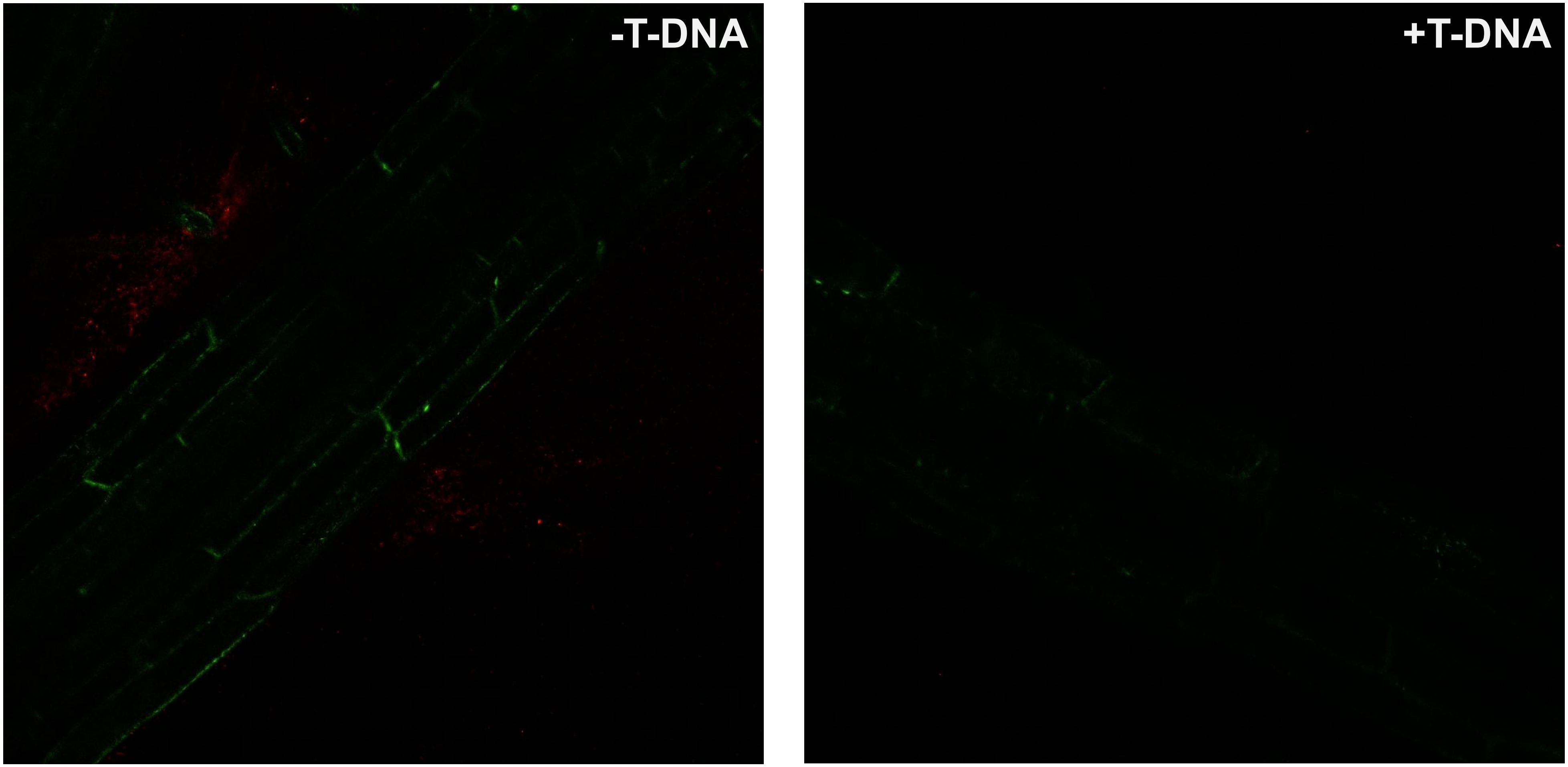


**Supplemental Figure S15. Infection of root cells expressing the myosin VIII-2 CBD with an *Agrobacterium* strain capable of transferring T-DNA re-localizes VirE2-Venus from the cellular periphery into the cytoplasm.** Transgenic plants inducibly expressing the myosin VIII-2 CBD and VirE2-Venus were treated with 5 µM β-estradiol for 24 hr, then inoculated with $10^8$ cfu/ml of the *virE2* mutant strain *A. tumefaciens* At1872 without (left panel) or with (right panel) T-DNA. Z-stack images of cells in the root elongation zone were taken after 8 hr of infection using a Zesis LSM 880 Upright Confocal Plan-Apo microscope using a 20×/0.8 objective.

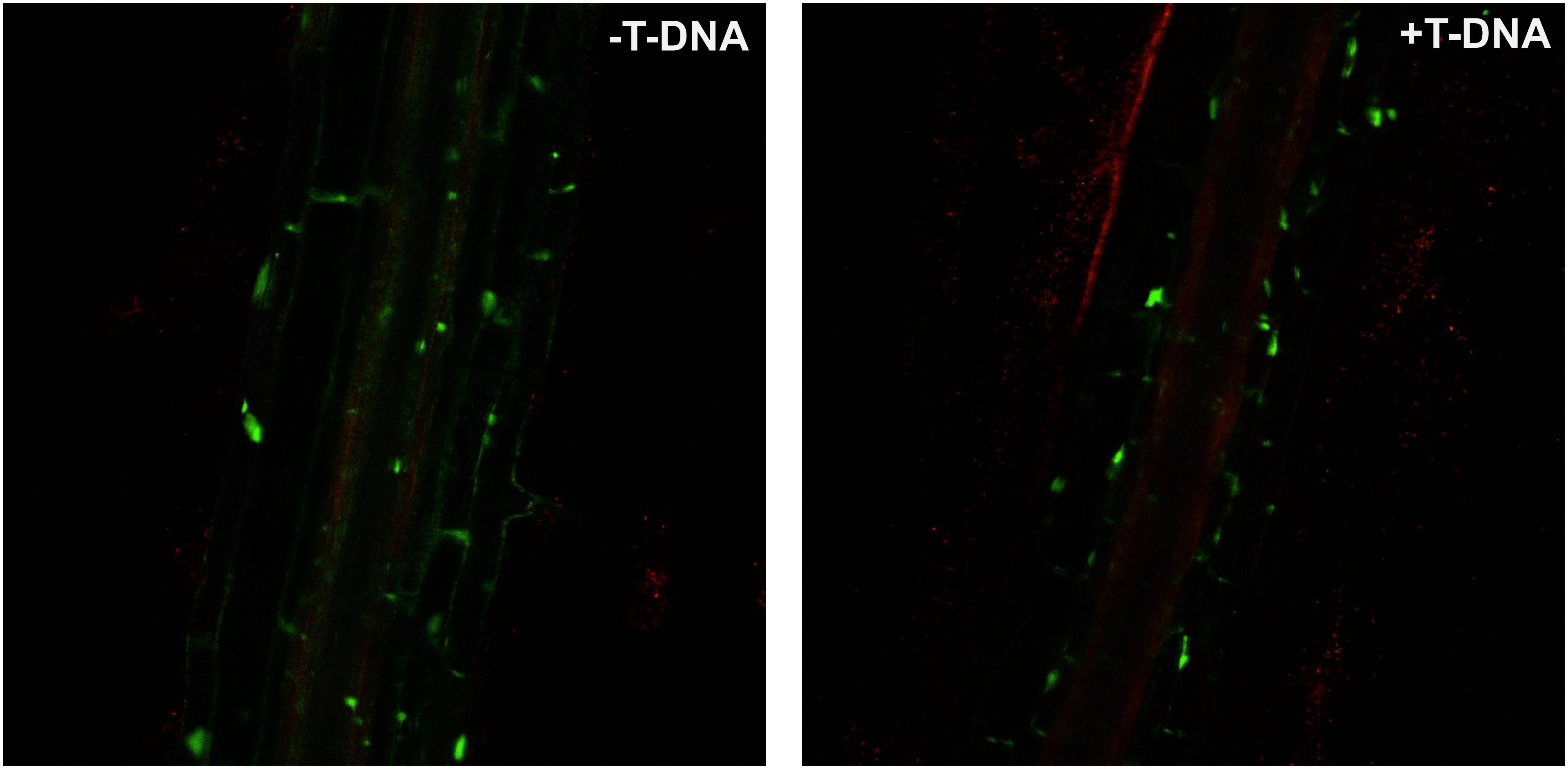


**Supplemental Figure S16. Infection of root cells expressing the myosin VIII-A CBD with an *Agrobacterium* strain capable of transferring T-DNA re-localizes VirE2-Venus from the cellular periphery into the cytoplasm.** Transgenic plants inducibly expressing the myosin VIII-A CBD and VirE2-Venus were treated with 5 µM β-estradiol for 24 hr, then inoculated with $10^8$ cfu/ml of the *virE2* mutant strain *A. tumefaciens* At1872 without (left panel) or with (right panel) T-DNA. Z-stack images of cells in the root elongation zone were taken after 8 hr of infection using a Zesis LSM 880 Upright Confocal Plan-Apo microscope using a 20×/0.8 objective.

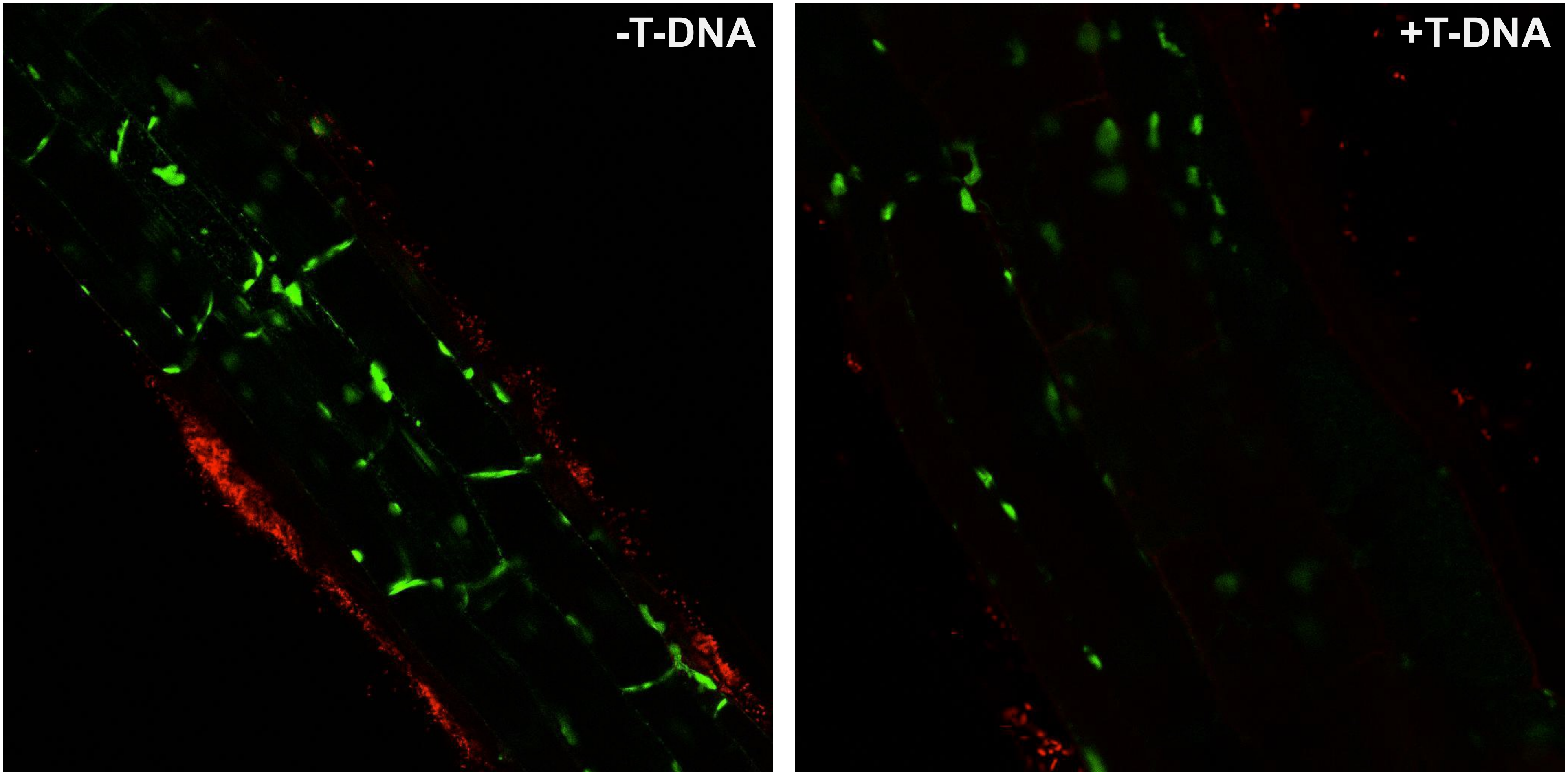


**Supplemental Figure S17. Infection of root cells expressing the myosin VIII-B CBD with an *Agrobacterium* strain capable of transferring T-DNA re-localizes VirE2-Venus from the cellular periphery into the cytoplasm.** Transgenic plants inducibly expressing the myosin VIII-B CBD and VirE2-Venus were treated with 5 μM β-estradiol for 24 hr, then inoculated with $10^8$ cfu/ml of the *virE2* mutant strain *A. tumefaciens* At1872 without (left panel) or with (right panel) T-DNA. Z-stack images of cells in the root elongation zone were taken after 8 hr of infection using a Zesis LSM 880 Upright Confocal Plan-Apo microscope using a 20×/0.8 objective.

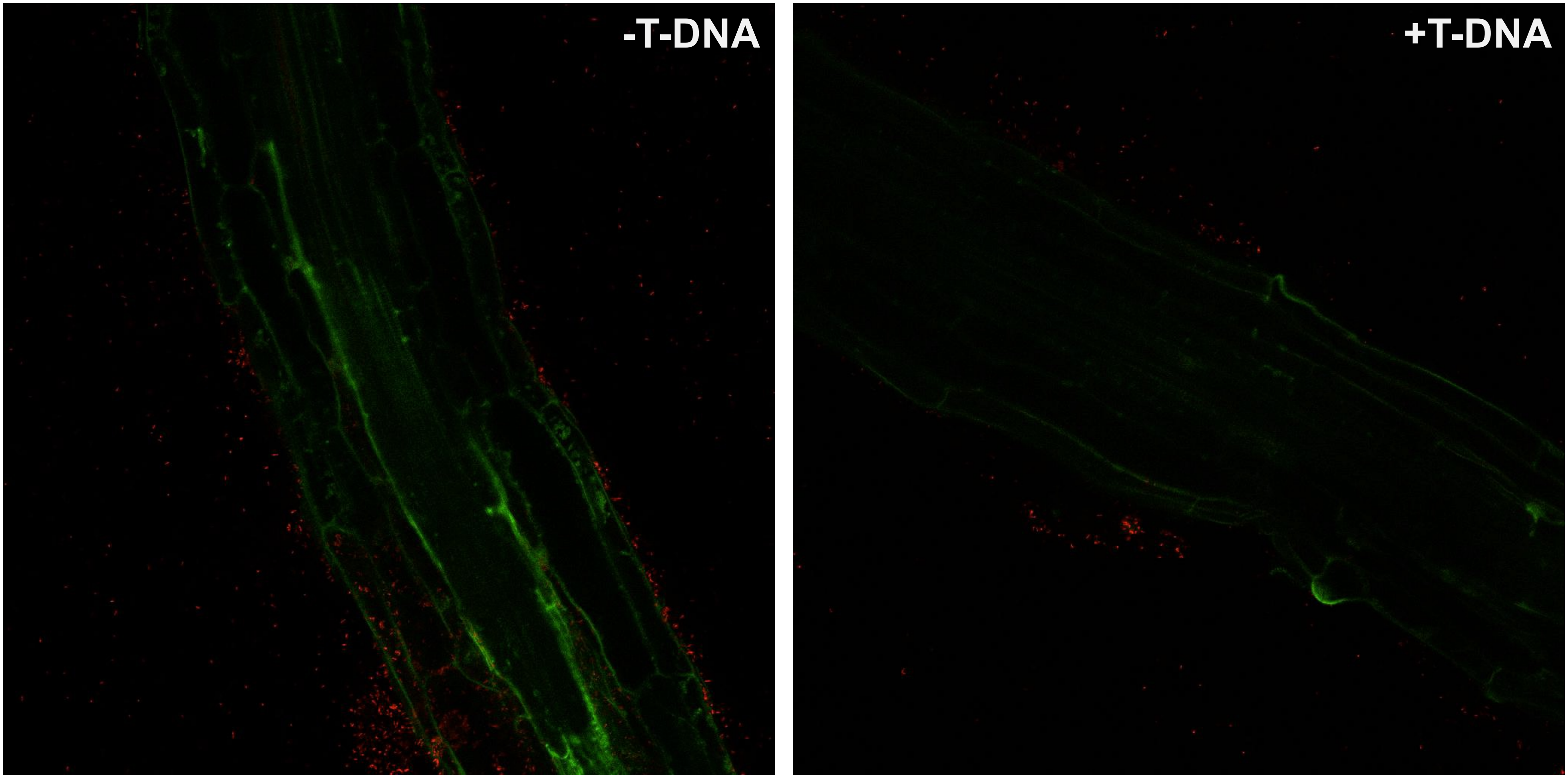


**Supplemental Figure S18. Expression of the myosin XI-K CBD does not affect VirE2-Venus localization upon infection by *Agrobacterium*.** Transgenic plants inducibly expressing the myosin XI-K CBD and VirE2-Venus were treated with 5 μM β-estradiol for 24 hr, then inoculated with $10^8$ cfu/ml of the *virE2* mutant strain *A. tumefaciens* At1872 without (left panel) or with (right panel) T-DNA. Z-stack images of cells in the root elongation zone were taken after 8 hr of infection using a Zesis LSM 880 Upright Confocal Plan-Apo microscope using a 20×/0.8 objective.

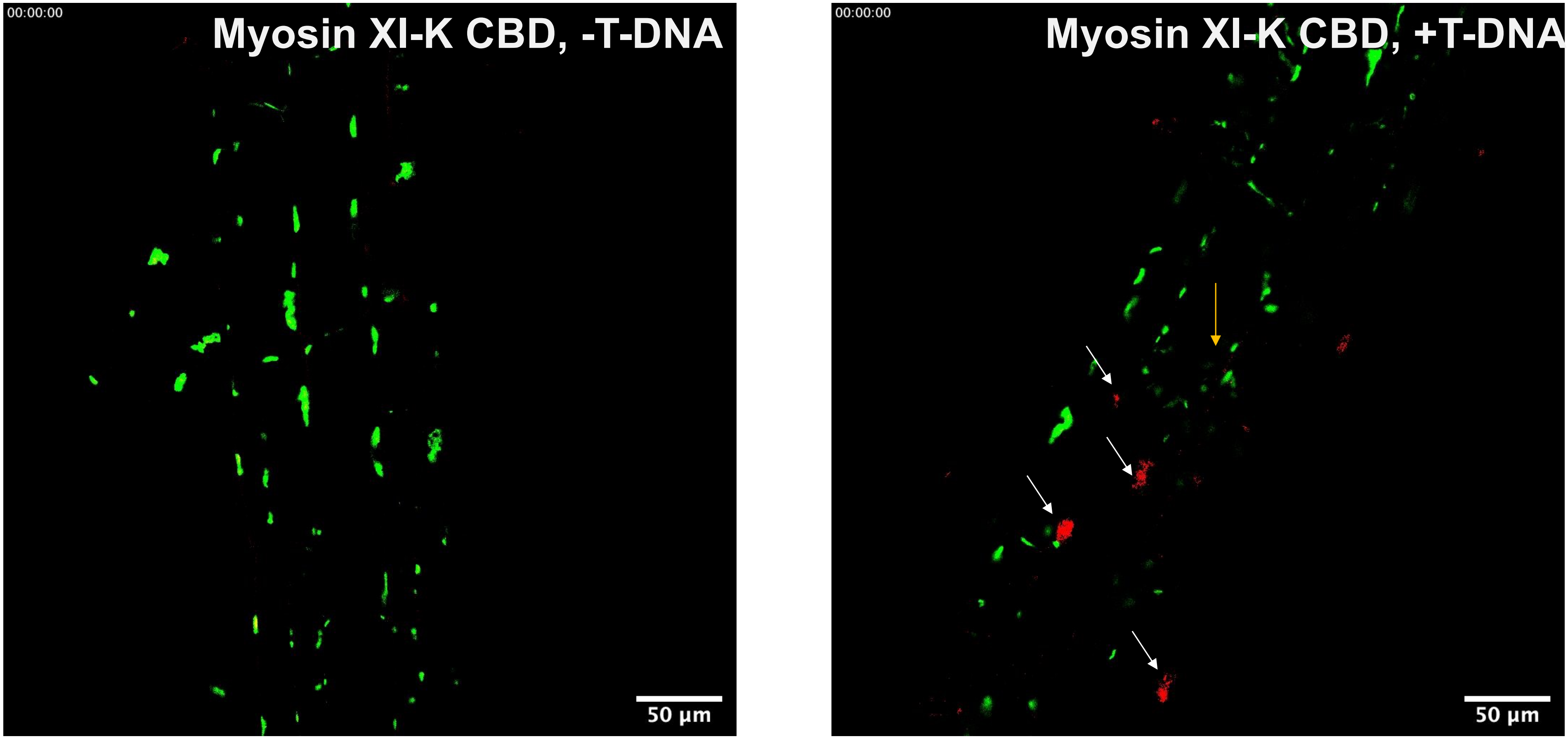


**Supplemental Figure 19. Expression of the myosin XI-K CBD inhibits movement of VirE2-Venus.** Transgenic plants inducibly expressing Myc-myosin XI-K CBD and VirE2-Venus were treated with 5 µM β-estradiol for 24 hr. Root segments were inoculated with $10^8$ cfu/ml of the *virE2* mutant strain *A. tumefaciens* At2405 harboring a T-DNA which constitutively expressed mCherry-NLS (to mark nucleus of transformed cells) (right panel) or without T-DNA (left panel). Time lapse images were taken after 24 hr of infection using a Zesis LSM 880 Upright Confocal Plan-Apo microscope with a 20×/0.8 objective. White arrows indicate nuclei, yellow arrow indicates a non-transformed cell. Bars indicate 50 µm.
